# Positional grammar of transcription factor binding partitions developmental and stress-response regulation in plants

**DOI:** 10.64898/2026.02.05.703842

**Authors:** Abraham Morales-Cruz, Sharon I Greenblum, Peng Wang, Yu Zhang, Lin Yang, Chris G Daum, Jenifer Johnson, Leo A Baumgart, Ronan C O’Malley

## Abstract

Understanding how transcription factor binding site (TFBS) position influences gene regulation remains a fundamental challenge in plants. Here, we integrate conserved multiDAP TFBS maps for 244 transcription factors (TFs) with single-nucleus chromatin accessibility, cell type-resolved gene expression, and hormone-response datasets across Brassicaceae species to determine how TFBS position relates to regulatory function. Although conserved TFBSs are enriched near transcription start sites (TSSs), TSS-proximal accessibility poorly predicts cell type-specific expression. Instead, cell type-specific expression correlates best with conserved TFBSs embedded in cell type-restricted chromatin, with TF family-specific distributions across distal promoters and introns. In contrast, TSS-proximal TFBSs in broadly accessible chromatin are associated with rapid transcriptional responses to abiotic and biotic stress hormones. Coding sequence TFBSs mark a distinct regulatory context in which the same DNA sequence encodes both amino acid sequence and TF motifs, including evidence that CDS-localized ABR1 binding may contribute to repression during hormone response. Finally, distal upstream regions contain conserved multi-family TF clusters with enhancer-like features overlapping rare cell type-specific accessible chromatin and enriched near genes controlling embryonic, meristematic, and hormone-dependent developmental patterning. Together, these results support a positional grammar in which TFBS position and chromatin context jointly partition developmental, stress-responsive, and repressive regulatory output in plants.

## Introduction

Understanding how local regulatory sequences shape gene expression has been a fundamental question in genomics, yet for plants, this relationship remains poorly characterized. In particular, comprehensive accounts of the impact of transcription factor binding site (TFBS) position across plant TF families are lacking^1,2^. This is due in part to challenges in collecting comparable large-scale multi-species datasets, as well as difficulties in differentiating functional cis-regulatory elements (CREs) from binding sites with limited regulatory impact. We recently leveraged our multiplexed DNA affinity purification sequencing (multiDAP) method to produce an atlas containing nearly 3,000 genome-wide TFBS maps across 360 TFs in up to 10 plant species spanning 150 million years of flowering plant evolution^3^. This dataset revealed that while TF binding specificities remain remarkably conserved across evolutionary time, the presence of individual binding sites varies considerably. A subset of TFBSs, however, are conserved across multiple species, and are strongly enriched for functional CREs. Yet whether the position of conserved binding sites relative to gene features is itself conserved, and whether such positional conservation shapes gene expression have not been examined.

It remains unknown, for example, how TF binding position and chromatin accessibility interact to shape cell type-specific gene regulation in plants. Plant tissues are constructed from diverse cell types whose expression programs must be precisely coordinated. Both chromatin accessibility and TFBS enrichment have been clearly linked to plant cell type-specific activity^4,5^. Functional assays also suggest that positional biases of cis-regulatory sequences may modulate the regulatory output of TF binding^1^. Well-established models in mammalian systems further implicate enhancer dynamics in shaping cell type specificity^6–8^. Understanding how this suite of cis-regulatory features shapes gene expression is a central question in genomics, and is particularly unresolved in plants. We test the hypothesis that conserved TFBS position and local chromatin context together define a positional grammar in which TSS-proximal, distal, intronic, and coding-sequence contexts are associated with distinct regulatory outputs.

In this study, we address several fundamental questions about how TFBS position shapes cell type-specific gene regulation in plants (**Fig. 1**). We first examine the positional distribution of conserved TFBSs relative to genes across all major plant TF families and identify TF ‘neighborhoods’. Next, we investigate the relationship between TFBS position and cell type-specific chromatin accessibility by generating a comprehensive multi-tissue, multi-species ATAC-seq dataset at single-nucleus resolution. Together, this resource reveals clear differences in both the relative position and relative accessibility of TFBSs between TF families. We then leverage our previously published high-quality single-cell transcriptomic data^3^ to evaluate the impact of TFBS location and accessibility on cell type-specific gene expression, and further generate a new hormone-treatment time-course RNA-seq dataset to examine the impact of TFBS position in the context of rapid stress response. We also examine coding-sequence TFBSs as a candidate context for transcriptional repression and identify conserved multi-family TFBS clusters in distal upstream regions with enhancer-like developmental signatures. Together, these analyses test a positional grammar model in which TFBS location and chromatin context partition developmental, stress-responsive, and repressive regulatory output.

**Fig. 1.**
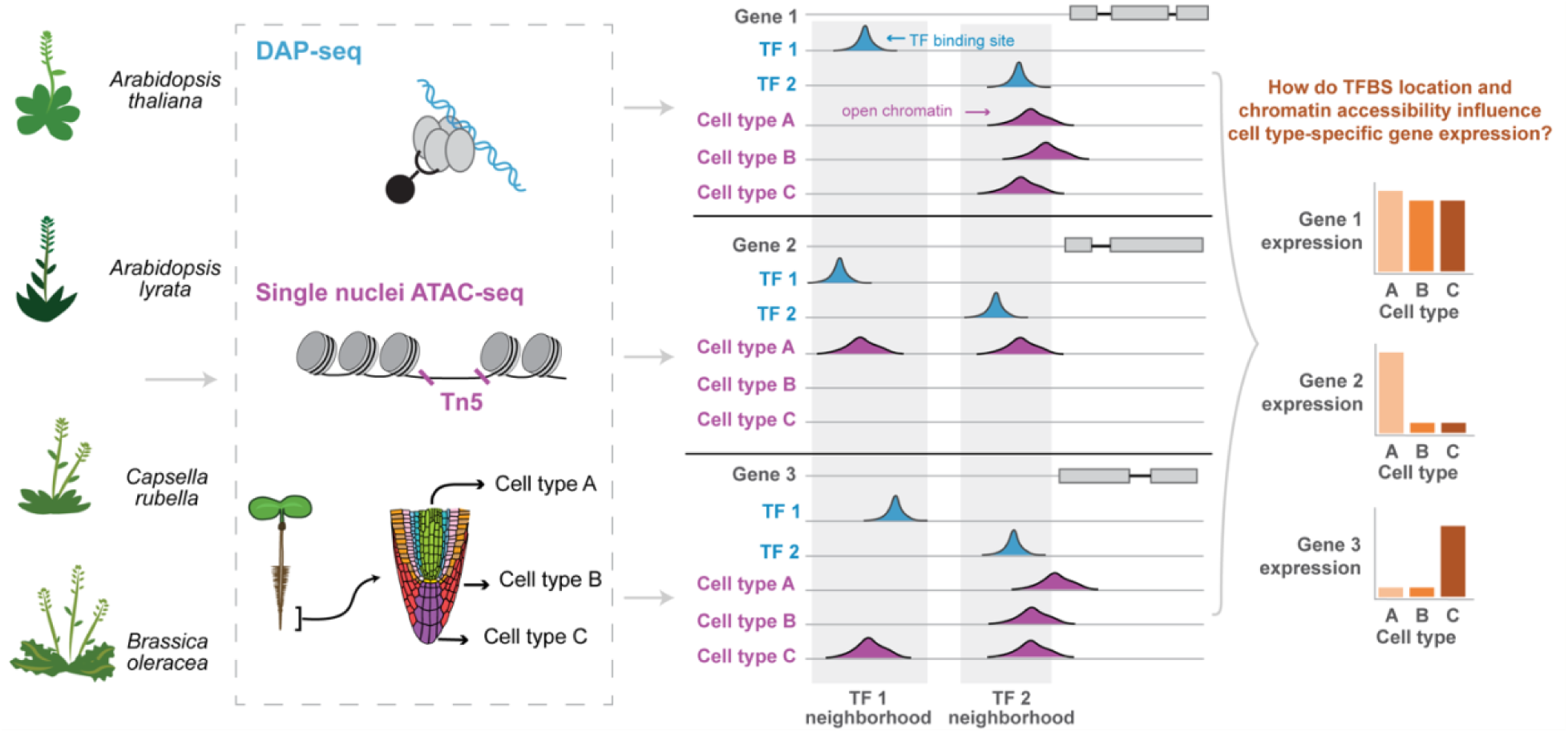
Decoding the impact of TFBS location and chromatin accessibility on cell type-specific expression. Schematic highlighting the cross-species integration of in vitro TF binding profiles, cell type-resolved chromatin accessibility tracks, and cell type-resolved gene expression profiles across Brassicaceae. This integrated workflow serves as the conceptual foundation for exploring how TF-specific positional binding preferences (TF "neighborhoods") and local chromatin accessibility influence cell type-specific gene expression.

## Results

### Profiling conserved cis-regulatory TF neighborhoods

We recently applied multiDAP in plants to investigate the conservation of TFBSs across two evolutionary time scales^3^. The first dataset maps binding sites of 244 TFs across four Brassicaceae species, representing approximately 21 million years of divergence. The second dataset encompasses 108 TFs in 10 angiosperm plant species, spanning both eudicots and monocots, and covers 150 million years of divergence. We found that conserved TFBSs were significantly enriched for features associated with CREs compared to lineage-specific sites, and exhibited patterns consistent with purifying selection within *A. thaliana* populations. We also developed a conservation score (c score) to quantify the number of species for which an orthologous gene promoter contains a binding site for the same TF^3^ (c1 indicates the TFBS was species-specific, c2 indicates a TFBS was conserved in two species, etc.).

In this study, we re-analyzed this data using a refined approach to enable exploration of TFBS position with higher precision, assigning each TFBS to a defined genomic region based on its position relative to a nearby gene (see Methods). We generated a genome-wide overview of TFBS positions by averaging TFBS density across all 244 TFs from the Brassicaceae family (hereafter termed “brassica”) multiDAP dataset relative to either the transcription start site (TSS) (**Fig. 2A**) or gene start codon (ATG) (**Fig. 2B**). In both cases, we normalized the average density of TFBSs to the genomic space available in the *A. thaliana* genome to show normalized values over a random background. We found four key features in these TFBS distributions. First, in agreement with previous genome-wide studies^9,10^, the primary peak of relative TFBS enrichment was situated within the 250 bp region upstream of the TSS. Second, the most conserved binding sites showed the highest enrichment around the TSS and ATG, and displayed densities that dropped off more sharply further upstream. Third, conserved TFBSs were enriched downstream of the start codon, including in coding sequences. Finally, we observed a region of TFBS depletion extending from approximately 50-100 bp upstream of the start codon up to the start codon itself. Interestingly, we did not see this pattern when using the TSS as the point of reference. We also noticed that this TFBS depletion overlaps a region of increased pyrimidine (thymine and cytosine) frequency in *A. thaliana* on the sense strand of the gene (**Fig. 2C**). We also evaluated regions downstream of the stop codon (see Data Availability section). Although prior studies^11^ report regulatory binding events in these 3’ regions, we observed very little evolutionary conservation, with only 4% of downstream peaks achieving a c4 score, prompting us to focus our subsequent analyses on the more prominent 5’ and coding features.

**Fig. 2.**
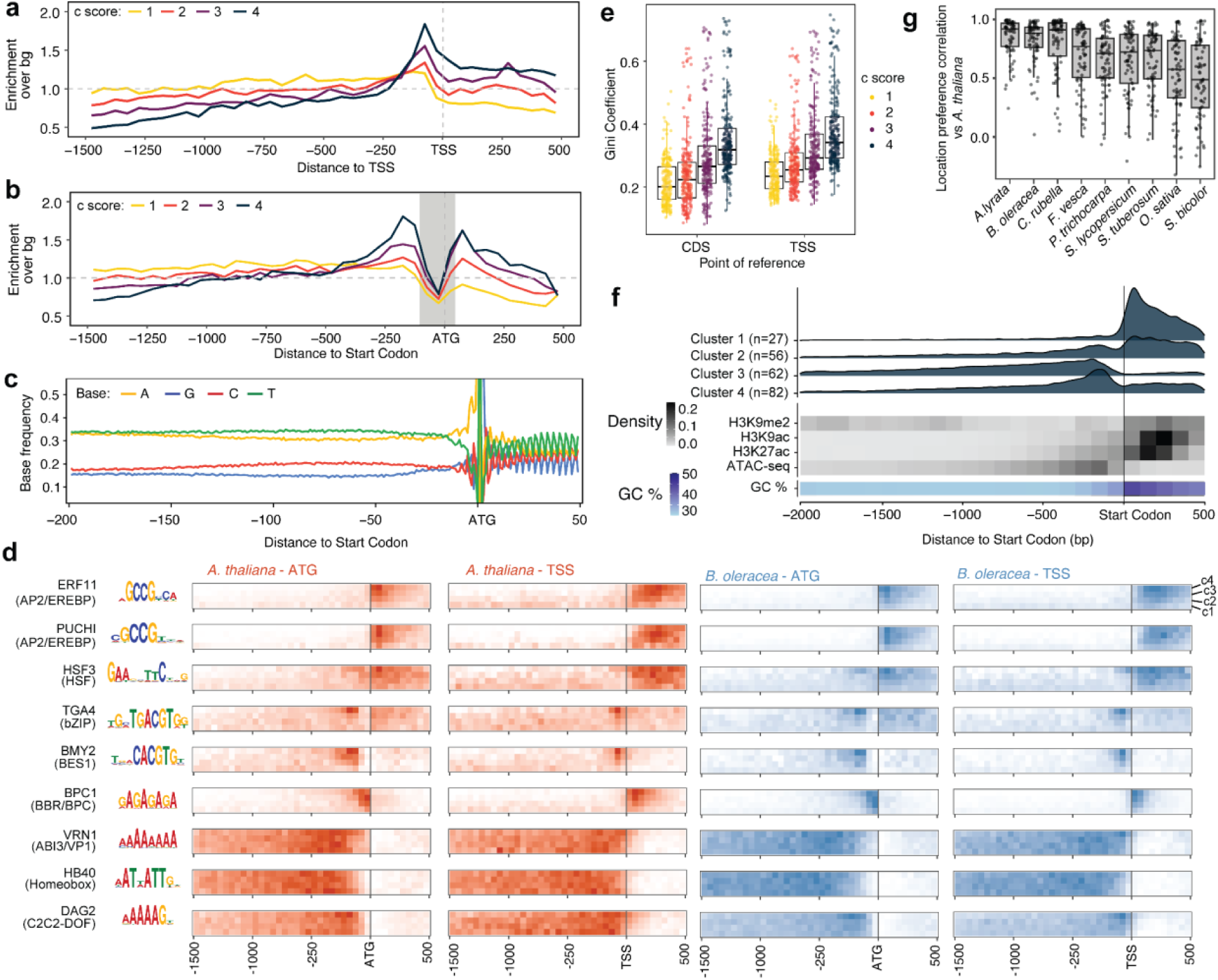
Distribution of transcription factor binding site (TFBS) position, stratified by conservation score across species. Normalized TFBS density at different c scores is shown relative to either the (**a**) transcription start site (TSS), or (**b**) start codon (ATG). Gray box indicates a TFBS-depleted region. (**c**) Nucleotide base frequencies around the TFBS-depleted region. (**d**) Examples of TFs with conserved TFBS position patterns, anchored at either the TSS or the start codon for *A. thaliana* (red, left panels) and *Brassica oleracea* (blue, right panels). Rows of each heatmap show positional distributions for TFBSs binned by c score. (**e**) Gini coefficients for different c scores and reference points, quantifying TFBS concentration around *A. thaliana* promoter regions. (**f**) TFs clustered by their TFBS distributions along promoter regions. (**g**) Pearson correlations of conserved (present in four or more species) TFBS density between *A. thaliana* and nine other plant species, including two monocots (rice and sorghum), across 50 bp windows surrounding the start codon.

To explore in more detail the binding patterns of individual TFs, we created separate TFBS density maps for each TF. We noted that different TFs and TF families displayed striking differences in their TFBS distributions, with most showing enrichment at specific genomic locations, often proximal to start codons and TSSs. We refer to these high-density regions as TF neighborhoods. For example, TGA4, BCP1, and BES1 bound preferentially to neighborhoods immediately upstream of start codons, ERF11 and PUCHI downstream of start codons, and DAG2, VRN1, and HB40 bound to a more diffuse neighborhood across the upstream region (**Fig. 2D**). TFBSs with higher c scores typically showed the strongest enrichment within a neighborhood (measured by the Gini coefficient), whereas c1 TFBSs typically showed minimal enrichment (**Fig. 2E**).

We observed very similar TF-neighborhood associations when mapping TFBSs of *Brassica oleracea* **(Fig. 2D**, blue heatmaps), as well as two additional brassica species (**Extended Data Fig. 1A**). TFs with GC-rich motifs tended to bind to neighborhoods located downstream of the start codon, whereas TFs with AT-rich motifs were more likely to bind to the upstream neighborhood (**Fig. 2D**). The TF BCP1 from the BBR/BCP family was the only TF in our dataset with preferential binding in the first 50 bp upstream of the start codon, a region that is generally depleted of TFBSs from all other tested TFs. Across all TFs, we observed a slight increase in the positional concentration of TFBS when anchored on the TSS vs the start codon (*t*-test, *p* = 0.012) measured by the Gini coefficient (**Fig. 2E**). However, a more focused analysis comparing TFBS density in the 150 bp surrounding the TSS vs the CDS identified distinct groups of TFs for which either the TSS or the CDS was the stronger driver of TFBS position (**Extended Data Fig. 1B**). Using promoter enrichment data from c4 TFBS neighborhoods, we next classified all TFs in the brassica multiDAP dataset into four distinct clusters (**Fig. 2F)**. These clusters capture distinct binding profiles: a strong preference for the gene body (cluster 1) or the upstream region (cluster 3), or more diffuse binding both upstream and downstream, with slight preference for the gene body (cluster 2) or the upstream region (cluster 4). Finally, we compared TFBS distributions of *A. thaliana* TFs within its own genome to their distributions across the other nine species included in our deep evolutionary dataset^3^ (**Fig. 2G, Supplementary Table 1 and Extended Data Fig. 1C**). We calculated pairwise correlations of conserved (c4+) TFBS density within promoter bins, revealing that the positional binding preferences of *A. thaliana* TFs are highly concordant between the *A. thaliana* genome and those of all other species. While median correlation coefficients for *A. thaliana*’s most distant relatives, *Oryza sativa* (rice) and *Sorghum bicolor* (sorghum), were lower than correlations to more closely related species, all correlations remained higher than 0.6, demonstrating that TF binding neighborhoods are a deeply conserved feature in plants (**Fig. 2G**).

### Epigenomic features associated with TF neighborhoods

The regions surrounding genes are tightly packed with epigenetic marks that vary in frequency across promoters and gene bodies, raising the possibility that the neighborhoods identified above influence or are influenced by these features (**Fig. 2F**). To determine if neighborhoods correspond to distinct chromatin environments, we intersected TFBSs with chromatin accessibility^12^, activating histone modifications^13–17^, and repressive epigenetic marks^18,19^ profiled in *A. thaliana* seedlings and quantified enrichment as a function of conservation score and TF family (**Extended Data Fig. 2A,B**). Because these published datasets were generated across different studies, we used them here to define broad chromatin-state context rather than matched measurements of tissue- or condition-specific regulatory dynamics. Across most TFs, highly conserved sites were enriched for accessibility and permissive histone marks, whereas low-conservation sites (c1) overlapped preferentially with silencing marks including DNA methylation- and H3K9me2-marked heterochromatin^18,19^. This supports a strong association between TFBS conservation and local chromatin state, with conserved regulatory elements generally found in permissive chromatin, while sites found only in one species are disproportionately associated with heterochromatin.

TF families also exhibited reproducible epigenetic patterns that mirrored their neighborhoods (**Extended Data Fig. 2A**). TFs with preferred binding in the gene body neighborhood (e.g., AP2/EREBP, LBD, and C2C2-GATA families) showed the highest overlap with permissive chromatin marks associated with gene starts, while families with enriched binding immediately upstream of TSSs (e.g., bZIP and bHLH) were strongly enriched in accessible chromatin. The particularly strong enrichment of GATA TFs with acetylated histones and other permissive marks is notable, as homologous GATA factors in animals regulate gene activity through direct recruitment of histone-modifying complexes^20^, suggesting plant GATAs may use a similar mechanism. In contrast, TF families with TFBSs enriched upstream of the gene, such as C2C2-DOF, Homeobox, and MYB-related, lacked association with accessible chromatin and activating chromatin marks. To quantify the relationship between preferred TF binding location and preferred epigenetic environment, we plotted the median location of c4 binding sites for each TF against their enrichment for each active chromatin mark and found strong correlations for most marks, including H3K36ac (Spearman r=0.84), H3K9ac (r=0.89), and H3K4me2 (r=0.96) (**Extended Data Fig. 2A,B**). Overall, the binding neighborhoods preferentially occupied by different TF families correspond to distinct chromatin states, suggesting that TF binding patterns are closely linked to the local epigenetic environment.

### Single-nucleus ATAC-seq identifies genomic context-dependent TF positioning

Our analysis of published bulk ATAC-seq data^12^ indicated that TF binding neighborhoods near the TSS are associated with accessible chromatin more than binding neighborhoods upstream or in the gene body (**Fig. 2F**). However, chromatin accessibility varies substantially across cell types^4,21–23^ and certain TFs are known to act in a cell type-specific manner^3^. It is thus unclear whether TFs that show little to no association with accessible chromatin in bulk ATAC-seq data (for example, C2C2-DOF, Homeobox, MYB-related families) may actually bind preferentially to chromatin that is accessible in a limited set of cell types. Motivated by this limitation, we generated a multi-species, multi-tissue snATAC-seq atlas of 74,307 high-quality nuclei from *A. thaliana* root and shoot seedling samples, and more than 10,000 high-quality nuclei from each of the other brassica species (**Fig. 3A**, **Extended Data Fig. 3**, **Supplementary Table 2**). We also profiled leaf and flower bud tissues, obtaining an average of 6,000 and 7,284 nuclei, respectively, across the four species. We assigned cell type labels to our snATAC-seq datasets by comparing gene accessibility profiles to gene expression profiles from our previously published snRNA-seq data^3^ (see Methods), resulting in 30 labeled cell types for *A. thaliana* seedlings, and 22-23 cell types for the other brassica species (**Supplementary Table 2**). We identified 30,907 accessible chromatin regions (ACRs) in promoter regions, of which most (n=29,160) were differentially accessible regions (DARs) (p < 0.01, regression test) (**see Data Availability section**). However, only 34.4% of the DARs met our stricter definition of ‘cell type-specific’ (accessible only in 1-3 cell types, **Supplementary Table 3**). Previous work has shown that such regions can work as gatekeepers, restricting binding access to specific TFs in specific cell types, thus playing a key role in shaping cell identity^23–25^. More than 67.9% of the cell type-specific DARs in our *A. thaliana* seedling dataset intersected one or more conserved (c4) TFBSs (**Supplementary Table 3**). For instance, in the promoter of CEPDL1, a gene known to promote nitrate uptake in the phloem^26^, we identified a phloem companion cell-specific DAR overlapping a c4 TFBS for DOF5.8, a key regulator of phloem development^26,27^ (**Fig. 3B**). Consistent with this model of TF-chromatin interactions, we found that the CEPDL1 gene was expressed exclusively in phloem/companion cells in our snRNA-seq atlas.

**Fig. 3.**
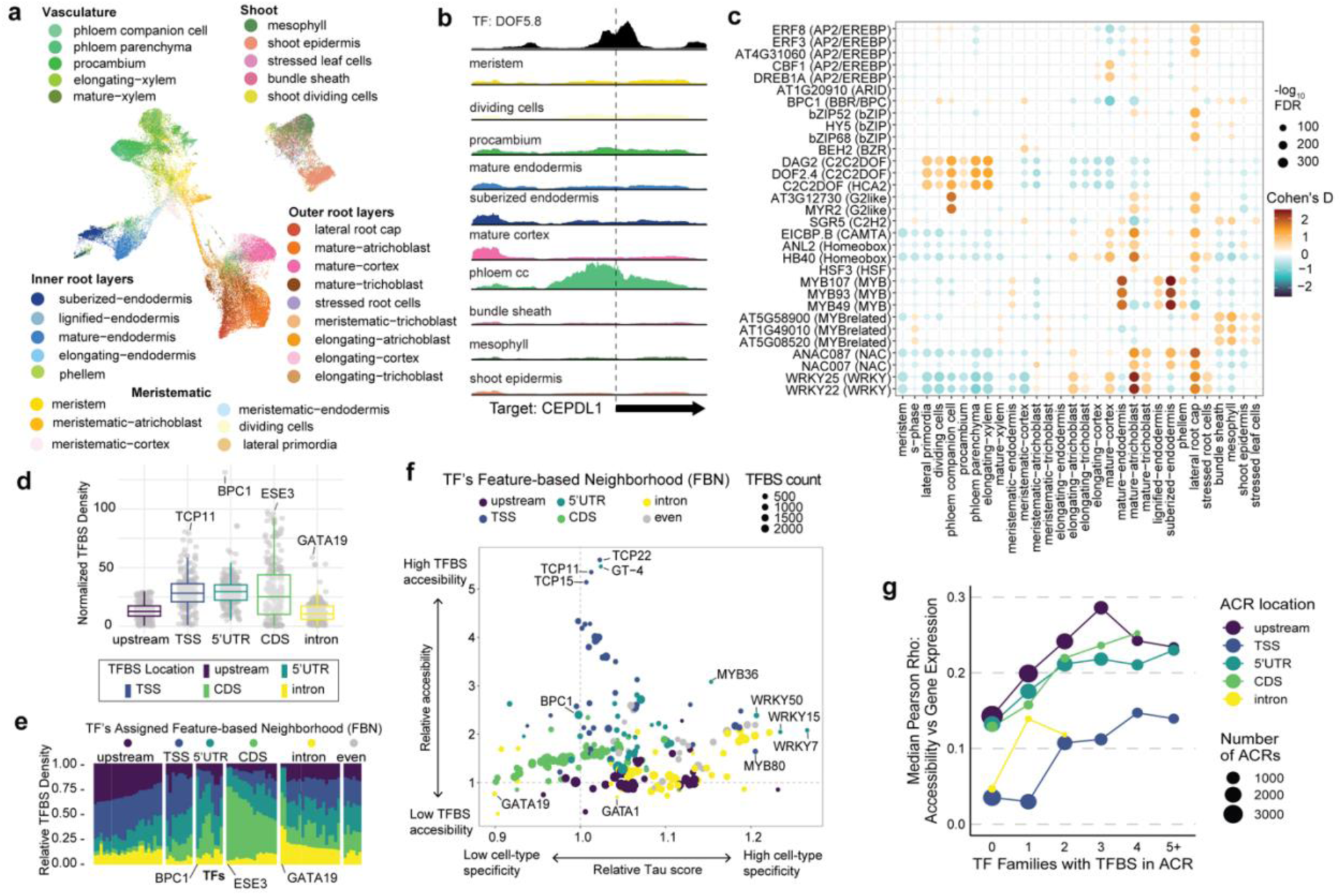
Single-nucleus ATAC-seq reveals TF interactions with open chromatin. (**a**) UMAP representation of the open chromatin profiles of 74,307 high-quality single *A. thaliana* seedling nuclei corresponding to 30 labeled cell types. (**b**) Open chromatin profiles across multiple cell types, around *A. thaliana* gene CEPDL1. The top track shows the binding profile of TF DOF5.8 as assayed by DAP-seq. (**c**) Heatmap showing enrichment or depletion of c4 accessibility for representative TFs (vertical axis) across labeled cell types in the *A. thaliana* seedling snATAC-seq atlas (horizontal axis). Dot size represents Wilcoxon p-value from rank sum test of TF accessibility scores between each cell type versus all other cells. Dot color indicates effect size of enrichment (red) or depletion (blue). Enrichments with FDR-corrected p-value < 0.01 are shown. (**d**) c4 TFBS density in different promoter features. Each dot represents the density of binding sites for a single TF in a given promoter feature, normalized by the TF’s total TFBS count and the total genomic space assigned to each feature. Densities shown were rescaled to represent 1,000 total TFBSs across 1 Mb of genomic sequence for each feature. Features were defined as follows: upstream: >200bp upstream of the TSS and within 2kb from the start codon, TSS: 0-200 bp upstream of the TSS, 5’ untranslated regions (5’ UTR): between the TSS and start codon, coding sequence (CDS): up to 500bp downstream of the start codon, intron: up to 500bp downstream of the start codon. (**e**) Stacked barplot of c4 TFBS density profiles for each TF across promoter features. Each column represents a single TF, and the sum of the TFBS density values are scaled to one. Up to 5 TFs per family are shown, grouped by assigned feature-based neighborhood (FBN). (**f**) Scatterplot showing relative accessibility and specificity for each TF in the brassica DAP-seq dataset labeled by each TF’s FBN. Size of the dot reflects number of TFBSs and colors show the TF’s FBN assigned in **e**. (**g**) Median Pearson correlation (y-axis) between chromatin accessibility in different promoter regions (colors) and gene expression computed across *A. thaliana* seedling cell types, as a function of the number of TF families with c4 TFBSs in the ACR. Point size represents the total number of tested ACRs. Only categories with at least 100 ACRs are shown.

To explore similar associations more systematically, we performed pairwise testing of the overlap between each TF’s c4 TFBSs and cell type-specific accessible chromatin, and recovered many expected TF-cell type associations (**Fig. 3C, Extended Data Fig. 4A**). For example, TFBSs for C2C2-DOFs^28^ were enriched in chromatin accessible in vascular tissue cells, Homeobox ANL2 in epidermal cells^29^, TCP3, TCP24^30^, and MYR2 (G2-like family) in phloem companion cells^31^, WRKYs in mature atrichoblast^32^, and MYB107 in suberized endodermis cells^33^. We extended this enrichment analysis to our other brassica atlases and found many shared instances of c4 TFBSs in cell type-specific accessible chromatin. Notable examples include WRKY40 TFBSs in cortex-accessible ACRs near orthologs of BIR1, a gene linked to root growth and defense^34,35^, and BPC1 TFBSs in atrichoblast-accessible promoters of BIN2, a regulator of atrichoblast differentiation^36,37^ (**Extended Data Fig. 4B**).

We next asked whether TFBS position and broader TF neighborhoods play an important underlying role in the associations we detected above. For this analysis, we adopted a slightly different definition of TF neighborhood, focused on the relative density of each TF’s binding sites within discrete genomic features: upstream regions, TSS-proximal regions, 5’ untranslated regions (5’UTR), coding regions (CDS), and introns (**Fig. 3D**; see caption for region definitions). Accordingly, we assigned each TF to one of six ‘Feature-Based Neighborhoods’ (FBNs) including a category for the 20 TFs with relatively even density across these features (**Fig. 3E**; see Methods). As with our distance-based neighborhoods, we observed clear differences between TF families. For example, the vast majority (42/43) of assayed TFs from the AP2/EREBP family were assigned to the CDS FBN, while the majority (12/15) of TFs from the BZIP family were assigned to the TSS FBN, almost all TFs (7/8) from the G2LIKE family were assigned to the intron FBN, and almost all (10/11) TFs from the Homeobox family were assigned to the upstream FBN.

To understand how these TF-specific associations with gene features relate to chromatin accessibility, we calculated normalized accessibility and cell type-specificity for chromatin regions intersecting each TF’s c4 TFBSs (**Fig. 3F**). At the extremes of the distribution we found TFs that tend to bind to highly accessible regions but do not show high cell type specificity, like multiple members of the TCP family (e.g., TCP22 and TCP11), as well as TFs that tend to bind to highly cell type-specific ACRs like members of the WRKY (e.g., WRKY7 and WRKY22) and MYB families (e.g., MYB80 and MYB93) (**Fig. 3F**). Our findings align with previous studies indicating that TCP transcription factors may be important regulators of chromatin accessibility as they are associated with both high gene expression and 3D genomic architecture^21,38^. TFs in the same FBN groups frequently showed similar relative accessibility and specificity of chromatin surrounding their TFBSs (**Fig. 3F**). TFs in the TSS FBN bound to the most accessible regions (t-test, *p* < 7.3×10⁻¹¹), followed by TFs in the 5′ UTR and CDS FBNs (**Extended Data Fig. 5A and B**). TFs with intronic TFBS enrichment (e.g., WRKY27 and MYB107) showed the highest association with cell type-specific chromatin (t-test, p < 0.013), while TFs with binding sites enriched in the CDS (e.g., PUCHI and ERF10 in the AP2/EREBP family) bound to the least cell type-specific chromatin (t-test, p < 5.9×10⁻⁶). Together, these observations raised the question of whether TSS-proximal accessibility, or accessibility in other regulatory contexts, best predicts cell type-specific expression.

To address this, we assigned each promoter ACR to a specific genomic feature and the closest gene. For each ACR-gene pair, we quantified the strength of association by computing Pearson correlations across cell types between chromatin accessibility and gene expression in our previously published *A. thaliana* seedling snRNA-seq atlas^3^, measured as pseudobulk transcripts per million (TPM). First, we found that, regardless of location, when an ACR contained TFBSs from a larger number of distinct TF families, ACR accessibility was more strongly predictive of gene expression across cell types.(**Fig. 3G**) We noted further that even at ACRs with diverse TF binding patterns, absolute correlation coefficients remained relatively modest (all median Rho values < 0.3), potentially reflecting inherent biological decoupling of chromatin accessibility and gene expression in some contexts. This is consistent with observations from the small set of *A. thaliana* same-cell RNA/ATAC (‘multiome’) studies published to date^39,40^. Grouping ACRs by their assigned feature revealed that the accessibility of chromatin centered in the region 200 bp upstream of the TSS consistently showed the weakest correlation with gene expression, while chromatin accessibility in any other feature was equivalently predictive. In line with this observation, chromatin in TSS regions was generally more accessible than chromatin in any other feature and only moderately cell type-specific (**Extended Data Fig. 5A,B**). We found similar correlations using a non-parametric (Spearman) correlation metric, when restricting correlations to the subset of cell types (n=16) with high overall chromatin accessibility, and when restricting to ACRs with high accessibility in at least one cell type **(Extended Data Fig. 5C)**. However, when limiting instead to ACRs effectively closed in at least one cell type, the correlation strength of TSS chromatin accessibility matched that of other regions. This pattern may arise because widespread constitutive promoter accessibility obscures a subset of dynamic promoters that undergo complete cell type-specific closure and thus preclude expression. Overall however, these results suggest that for many genes, chromatin accessibility near the TSS may not be the limiting factor for cell type-specific gene expression.

### TFBS location influences cell type-specific gene activity

Previously, we used multiDAP data to infer a TF’s cell type-specific activity from the expression patterns of its conserved target genes^3^. In that analysis, we computed a ‘TF activity score’ for each TF in each cell type by summarizing expression over all genes harboring a TFBS in their promoter. The analyses above reveal that TFs vary widely in binding density along the promoter, raising the possibility that the position of a binding site carries additional information about its regulatory impact. We first hypothesized that certain TFs can best exert regulatory influence when bound to a specific location within the promoter, with areas rich in conserved TFBSs representing a ‘sweet spot’ for the TF’s regulatory activity. If this model applies to cell type-specific regulation, we would expect binding sites in a TF’s FBN to show the strongest impact on gene expression. To test this, we binned each TF’s target genes by location of the TFBS, and computed TF activity scores separately for target genes in each bin. Examining activity scores in cell types where the TF itself was strongly expressed, we found that a TF’s FBN was slightly but significantly less likely to contain c4 TFBSs that impact cell type-specific gene expression (log_2_-fold change = -0.34; chi-sq test p < 0.04) (**Fig. 4A**). We repeated this test for target subsets at all c scores and found similar results. c1 TFBSs showed the strongest depletion of impactful binding sites in the FBN region (log_2_-fold change = -2.02; chi-sq test p < 4.0x10^-6^) (**Fig 4A**), suggesting that species-specific binding sites in the FBN region play little role in shaping cell type identity. We also repeated this test individually for TFs assigned to each FBN region; only TFs with an FBN in the 5’ UTR showed significantly higher activity scores for FBN-located binding sites, and only for conserved (c3 and c4) binding sites (**Extended Data Fig. 5D**). Thus, high TFBS density within a TF’s preferred neighborhood does not necessarily translate to the strongest cell type-specific regulatory impact. Although conserved TFBS density identifies preferred genomic territories for each TF, regulatory impact within those territories appears to depend on chromatin and feature context rather than density alone.

**Fig. 4.**
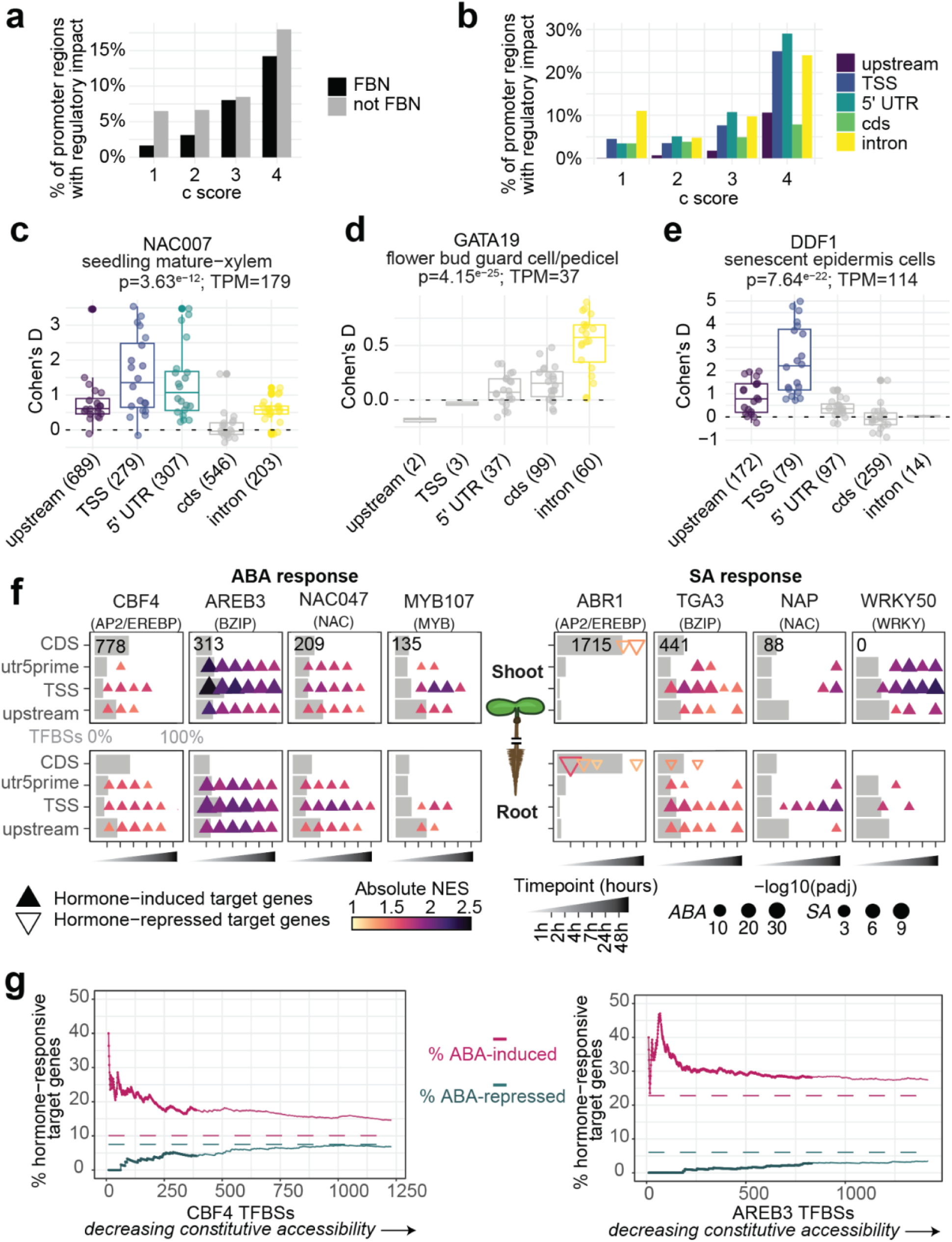
TFBS location carries information for gene expression regulation. (**a**) Barchart showing that TFBSs in each TF’s feature-based neighborhood (FBN) (black bars) are no more likely to impact gene expression than TFBSs in other promoter features (gray bars). The y-axis shows the percent of TF-cell type-feature trios marking genes with significant cell type-specific expression, out of all TF-cell type pairs showing strong TF activity from at least one feature. (**b**) Barchart showing the percent of TF-cell type-feature trios marking genes with significant cell type-specific expression as a function of TFBS location. (**c**) TF activity scores (measured as Cohen’s d; y-axis) representing the magnitude of mature-xylem-specific expression of groups of NAC007 target genes, binned by the location of their c4 TFBSs (x-axis). X-axis labels show the total number of NAC007 target genes harboring c4 TFBSs in each promoter feature. Each dot represents the Cohen’s d value for a randomly-selected subset of 25 genes. The plot title shows the p-value from an ANOVA test of Cohen’s d values as a function of promoter feature, as well as pseudobulk expression (TPM) of the NAC007 TF in mature xylem cells. (**d**) and (**e**) show additional examples as in (c) for two other TFs in two other cell types. (**f**) Gene set enrichment analysis for target genes of hormone-responsive TFs among genes ordered by ABA response (left) or SA response (right) at each timepoint in each tissue (top: shoot, bottom: root). Separate enrichment analyses were conducted for non-overlapping subsets of target genes binned by the location of their TFBSs. Results for target genes with TFBSs in multiple regions are not shown. Gray bars in each plot represent the percent of target genes in each TFBS-location bin, and absolute target gene count for the CDS bin is shown for reference. NES=Normalized Effect Size. (**g**) Enrichment of ABA-responsive genes at the 1 hr timepoint in shoot tissue, among TFBSs ordered by constitutive accessibility for two top ABA-response TFs, AP2/EREBP TF CBF4 (left) and bZIP TF AREB3 (right). All TFBSs in snATAC-seq ACRs are shown along the x-axis; TFBSs in constitutively accessible ACRs are marked with larger points. The y-axis shows the cumulative % of ABA-induced genes (pink) or ABA-repressed genes (teal) encountered as larger subsets of TFBSs are considered. The x-axis begins at a subset size of 10 to mitigate choppiness from low counts.

This unexpected negative association between FBNs and cell type-specific regulation suggested that while high TFBS density may not be indicative of a regulatory ‘sweet spot’, a TFBS’s location may still carry important regulatory information. To explore this, we conducted an ANOVA test for each TF-cell type pair described above, asking whether variance in target gene expression could be explained by position of the TFBS. This test was significant (FDR-adjusted p<0.05) for 91% of the tested pairs (208/227), suggesting that while the FBN might not explain differences in TFBS regulatory impact, differences do indeed exist. We next asked whether TFBSs in a specific feature showed high regulatory impact generally, when summarised across all TFs. We found that as a whole, c4 TFBSs in introns, TSS regions, and 5’ UTRs all showed a relatively higher impact on developmental regulation compared to TFBSs further upstream of the TSS or in coding sequence (**Fig. 4B**). This trend was repeated for TFBSs at every c score, as well as when increasing the stringency of the threshold for regulatory impact. In both cases, *A. thaliana*-specific (c1) TFBSs in introns specifically remained notably more impactful than c1 TFBSs in other regions. While the regulatory role of introns has been previously observed^41,42^, our findings further suggest that intronic TF binding sites may be an important source of regulatory novelty.

The results above indicate that TFBS position carries TF-specific regulatory information. To illustrate this specificity, we examined cases where target genes with TFBSs in different promoter regions showed markedly different regulatory impact. We started with TFs previously observed to have the clearest cell type-specific activity^3^. For example, both the target genes of NAC007 and the TF itself were strongly and specifically expressed in the mature-xylem. Yet examining subsets of target genes from different promoter regions revealed that unlike TFBSs in all other regions, genes harboring TFBSs in coding sequence did not show significant expression in this cell type, despite the fact that more TFBSs were found in this region than three others (**Fig. 4C**). A second example, GATA19, a TF known to regulate flower and shoot apical meristem development, specifically showed the strongest regulatory influence from TFBSs in intronic regions in the two cell types in which the TF was strongly expressed (both floral cell types) (**Fig. 4D**). This finding echoes previous results^1^ noting the importance of binding sites downstream of the TSS for GATA TFs, and indeed we found vanishingly few conserved GATA19 binding sites upstream of the TSS. Here we further show that among downstream TFBSs, intronic binding sites specifically may confer the strongest cell type-specific regulation. A third example (AP2/EREBP TF DDF1, a TF involved in stress response, expressed specifically in senescent leaf epidermis cells) showed an opposite pattern, whereby TFBSs in both intronic and CDS regions showed little impact on target expression, while TFBSs in the TSS region were by far the most impactful, followed by TFBSs further upstream (**Fig. 4E**).

Taken together, these observations suggest that complex dynamics may govern interactions between the location of TF binding and target gene transcription. Despite the complexity, we wondered whether these dynamics were preserved across closely related species. To test this, we calculated TF activity scores for the same set of cell types and promoter regions, using TFBSs and snRNA-seq atlases from three other brassica species^3^. Comparing activity scores from each species to *A. thaliana*, we found significant Pearson correlations in each case, with coefficients ranging from 0.51 in *C. rubella* to 0.59 in *A. lyrata*. To test the contribution of TFBS location specifically, we then shuffled *A. thaliana* activity scores across promoter regions within the same TF-cell type pair (**Extended Data Fig. 5E**) and re-calculated each pairwise correlation. We found that the coefficient dropped by half for each of the three species (coefficient log2-fold change = -1.1, -1.0, -1.1 for *A. lyrata*, *C. rubella*, and *B. oleracea* respectively), strongly suggesting that TFBS location contributes to gene regulation in a conserved, position-dependent manner. This conservation of position-specific regulatory effects across species supports the broader concept of a positional grammar for plant gene regulation.

### Stress response CREs are concentrated in constitutively accessible TSS regions

Although conserved TFBSs were strongly enriched near the TSS, and TSS-region ACRs were among the most broadly accessible chromatin regions across cell types, TSS-proximal accessibility was weakly correlated with cell type-specific gene expression (**Fig. 2A**; **Fig. 3D,F,G**; **Extended Data Fig. 5A,B**). This raised the possibility that many broadly accessible TSS-proximal TFBSs do not primarily define cell identity, but instead function in some other programs.

Consistent with this, several stress-associated TF families, including bZIP, AP2/EREBP, WRKY, and NAC factors, were enriched in TSS-proximal regions (**Fig. 2D,F**; **Fig. 3D,E**). Prior computational analyses have also noted that these stress-associated TF families were enriched near the TSS, and ChIP-seq studies of abiotic and biotic stress-response TFs have shown TSS-proximal binding enrichment in stress-regulated networks^9,10,43,44^. We therefore asked whether genes with conserved TFBSs in broadly accessible TSS-proximal regions are preferentially associated with hormone-responsive stress programs. To test this, we treated *A. thaliana* plants with phytohormone regulators of abiotic and biotic stress-response programs: abscisic acid (ABA) and salicylic acid (SA), respectively ^45,46^. We conducted treatment time-courses for each hormone and profiled gene expression changes via bulk RNA-seq in both root and shoot tissues at multiple timepoints. Principal component analysis confirmed that each hormone treatment initiated a clear and differentiable genome-wide transcriptional response throughout the time-course (**Extended Data Fig. 6A**). We first examined the results from treatment with ABA, a master regulator of responses to abiotic stressors including drought, cold, and soil salinity^45^. As expected, we found that known ABA-responsive AP2/EREBP TFs such as CBF4, DREB1A, and RAP2.1 were among the top TFs exhibiting rapid ABA-induced expression (**Extended Data Fig. 6B**), and likewise conserved target genes of many AP2/EREBP TFs were enriched for up-regulated expression, along with target genes of bZIP, MYB, and NAC TFs (**Extended Data Fig. 6C**).

To investigate the impact of TFBS location on expression regulation, we grouped each TF’s target genes by TFBS position, and conducted a gene set enrichment test to assess whether each group was significantly enriched for hormone-induced or hormone-repressed genes. Across many major ABA-responsive TF families, we found that genes with TFBSs in the TSS region showed the strongest signal of hormone-induced expression (**Fig. 4F, Extended Data Fig. 7A**). Turning instead to the SA response results, this same pattern was largely repeated for a separate set of known SA-responsive bZIP TFs^46^ including TGA1/3/4 which all showed strong early induction of genes harboring TSS-proximal TFBSs (**Extended Data Fig. 7B**), as well as key NAC and WRKY TFs known to be part of the SA signaling cascade (**Fig 4F**). We noted that the location of TFBSs in critical hormone-induced target genes appeared more consistently centered near the TSS than the location of top cell type-specific TFBSs. Indeed, comparison of gene set enrichment scores for different promoter regions revealed that TFBSs in the TSS region were the most enriched for hormone response across all major hormone-responsive TF families, while the location of top enrichments for cell type-specificity were far more variable (**Extended Data Fig. 7C**). Given our findings above that chromatin in the TSS region was unique in its broad accessibility (**Extended Data Fig. 5A**) and weak correlation with cell type-specific expression (**Fig. 3G**), we reasoned that TFs mediating rapid, tissue-wide responses might preferentially operate through constitutively accessible sites.

To test this, we examined the accessibility of TFBSs for two of the top ABA-responsive TFs: CBF4 (AP2/EREBP family) and AREB3 (bZIP family). To specifically differentiate constitutively accessible binding sites, we first looked at the distribution of all ACR accessibility values in each cell type, determined that a mean accessibility of 0.2 formed a robust distinction between accessible and inaccessible sites, and defined ‘constitutively accessible’ ACRs as those with accessibility > 0.2 in at least 29/30 cell types (**Extended Data Fig. 8A**). We noted that a small subset of primarily bZIP TFs, including AREB3, showed a uniquely high proportion of TFBSs located in constitutively accessible chromatin (**Extended Data Fig. 8B**). We then ordered each TF’s c4 TFBSs by accessibility, first including all TFBSs in constitutively accessible ACRs sorted by mean accessibility, followed by all other TFBSs in ACRs, and finally TFBSs not in ACRs. Using this ordered list and hormone-response data from the 1 hr timepoint in shoot tissue (where ABA-response for both TFs was strongest), we found that for both CBF4 and AREB3, ABA-upregulated genes were strongly preferentially distributed at the beginning of the list, in regions with the highest broad chromatin accessibility (**Fig 4G**; GSEA p-values 0.026 and 0.00023 for CBF4 and AREB3 respectively). Together, this supports a model in which one major functional role of conserved constitutively accessible TSS-proximal TFBSs is to mediate rapid global transcriptional responses to environmental stress.

### Coding-sequence TFBSs mark a candidate repressive context

We were intrigued to observe that despite a strong enrichment for conserved AP2/EREBP TFBSs in CDSs, target genes carrying these TFBSs rarely showed evidence of cell type-specific AP2/EREBP regulation. We identified three potential factors that may contribute to this: 1) Coding sequence is subject to strong purifying selection to preserve gene function, and thus our conservation test may have limited ability to distinguish sequence conserved primarily due to regulatory impact in these regions, 2) CDS TFBSs may repress expression, but single-cell transcriptomic data is too sparse to detect this, or 3) as some AP2/EREBP TFs play important roles in stress responses, their activity may be muted in non-stressed conditions.

Our bulk-RNA hormone time course data gave us a unique opportunity to distinguish between these possibilities. We found that target genes harboring AP2/EREBP TFBSs exclusively in coding sequence showed no significant repression in response to ABA and only rarely showed significant up-regulation (**Fig 4F; Extended Data Fig. 7A**). Among the 13 tested AP2/EREBP TFs whose expression was significantly induced by ABA treatment (adjusted DESeq p-value < 0.05, log2-fold change > 1), only two (ERF4 and ERF8) showed significant activation of target genes harboring CDS TFBSs, and this enrichment was only observed in one tissue at a single timepoint (root, 24hr timepoint). Extending our analysis to other ABA-responsive TF families with various enrichment for CDS binding (e.g., bZIP^44^, NAC, MYB)(**Fig 4F)**, we found a similar lack of ABA-induced or ABA-repressed expression for target genes with TFBSs solely in coding sequence. Turning instead to the SA response results, we first examined known SA-activated bZIP TFs^45^ including TGA1/3/4 which all showed strong early target gene induction (**Extended Data Fig. 7B**), as well as key NAC and WRKY TFs known to be part of the SA signaling cascade (**Fig 3F**). For a few bZIP TFs, target genes with CDS TFBSs in coding sequence were moderately repressed, but across key established SA-responsive TFs in these three TF families, the dominant response was from target genes with TFBSs in or near the TSS.

Finally, we examined target genes of a known AP2/EREBP family transcriptional repressor, Abscisic Acid Repressor 1 (ABR1), whose expression was rapidly induced in root tissue by SA in our data (**Extended Data Fig. 6B**). Strikingly, ABR1 target genes with TFBSs exclusively in coding sequence (including ABA-response TF CBF4) showed a strong concurrent drop in expression in root tissue at early timepoints and later in shoot tissue (**Fig 4F**). Taken together, these results are consistent with a model of rapid antagonistic response involving repression via CDS binding. As ABR1 TFBSs downstream of the start codon occur almost exclusively in coding sequence (∼99%; ∼1% intronic), we propose that CDS binding represents a candidate mode through which ABR1 may contribute to transcriptional repression. TFBSs within protein-coding sequence, termed dual-use codons (duons)^73^, were first inferred from genome-wide evolutionary patterns in the human genome, but that study did not assign regulatory function to individual sites. Subsequent studies have linked individual duons in plants^48^ to transcriptional effects at a small number of loci. However, duons have not previously been characterized as a genome-wide regulatory context for any plant transcription factor.

### Distal cis-regulatory modules show hallmarks of developmental enhancers

We next investigated TFBSs in ACRs located in distal genomic regions, defined as the interval starting at 2 kb upstream of the start codon and extending up to 10 kb upstream, while avoiding regions previously assigned to the promoter of other genes (see Methods). We identified approximately 4.4 million TFBSs across all four brassica species mapping to distal regions, and found that 11.1% were conserved across all four species (distal c4 TFBSs). This was notably lower than the observed 30% rate of c4 TFBSs in promoters for the same TFs. We focused on cis-regulatory modules (CRMs)^2,50^, defined here as clusters of distal c4 TFBSs from three or more families within 300 bp of each other (see Methods). This heterotypic composition criterion was designed to capture combinatorial TF input characteristic of enhancer-like elements. We identified between 3,333 and 6,409 CRMs across the four brassica species (**Fig. 5A**). A complete catalog of all 3,333 A. thaliana distal CRMs is provided in **Supplementary Table 4**. In *A. thaliana*, we found that distal CRMs averaged 651 bp in length (**Fig. 5B**) and contained an average of 25.6 TFBSs from 5.3 TF families (**Fig. 5C**), with comparable values across the other species. The average distance to the downstream gene’s start codon ranged from 3,705 bp to 4,936 bp in *A. thaliana* and *B. oleracea*, respectively (**Fig. 5D**). Finally, we found a significant enrichment of H3K27me3 (an epigenomic mark associated with enhancers in plants^51^) in distal CRMs of *A. thaliana* (1.45 fold-change, p < 0.002, permutation test; **Extended Data Fig. 9A**; per-CRM data in **Supplementary Table 4**).

**Fig. 5.**
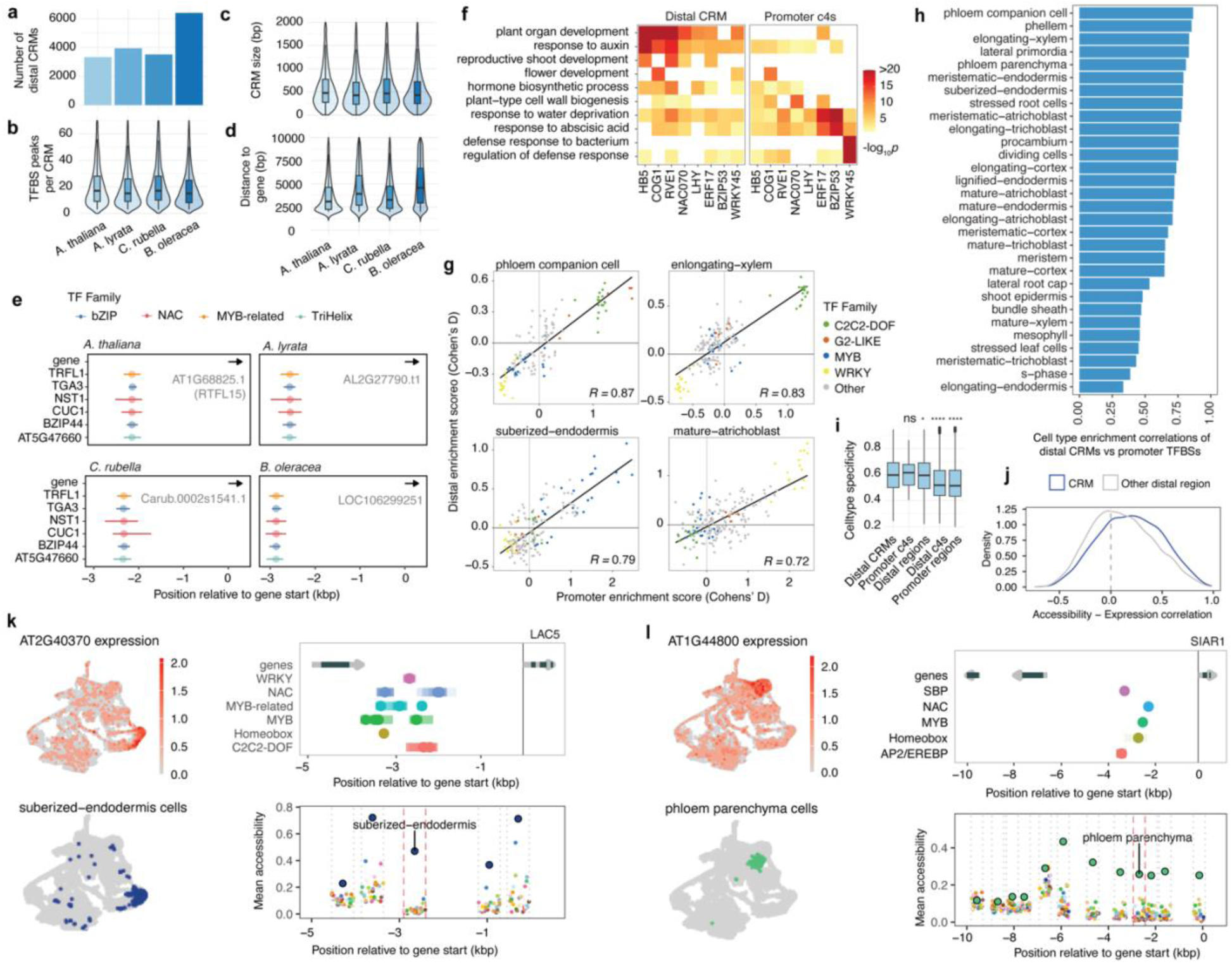
Identification of potential CRMs in distal ACRs across four species. **(a)** CRM count, (**b**) TFBS number, (**c**) genomic length, and (**d**) distance to the closest downstream gene, shown for each of four species. (**e**) GO terms significantly enriched in the genes associated with TFBSs in the promoter region or distal CRMs for a set of representative TFs. (**f**) Example showing conservation of TFBSs from the same TF families in a CRM upstream of four orthologous genes. (**g**) Pearson correlation between open chromatin enrichment for TFBSs in promoter regions vs. distal CRMs. (**h**) Pearson correlation of overall enrichment scores (Cohen’s d) for distal CRMs vs promoter TFBSs within each cell type. (**i**) Cell type specificity (Tau score) of chromatin accessibility in different genomic features. Asterisks indicate statistical significance between distal CRMs and other categories. ns: p > 0.05, *: p ≤ 0.05, **: p ≤ 0.01, ***: p ≤ 0.001, ****: p ≤ 0.0001. (**j**) Distribution of Pearson correlation coefficients measuring concordance of ACR accessibility and gene expression for ACRs overlapping distal CRMs and ACRs in any other distal region. (**k**) Example of a gene (LAC5) showing strong suberized-endodermis-specific expression (UMAPs on the left), harboring a distal CRM specifically accessible in suberized endodermis cells (plots on the right). The top right plot shows a diagram of TFBS locations (x-axis) for different TF families (y-axis) relative to the LAC5 start codon (x=0). Gray and black shaded regions of the arrows on the top line show LAC5 non-coding and coding regions, respectively, and the closest upstream gene. Only c4 TFBSs located in LAC5’s defined distal region (>2 kb upstream of the start codon) are shown. The bottom right plot has the same x-axis, but shows mean chromatin accessibility for genomic windows in different cell types (colors). Genomic windows with a detected ACR are outlined in gray. The genomic window overlapping the CRM is outlined in red, and the most accessible cell type is labeled. (**l**) Same as (k) but showing an example of a gene (SIAR1) expressed predominantly in phloem parenchyma cells harboring a CRM specifically accessible in phloem parenchyma.

We identified conserved CRMs as those with the same TF composition upstream of orthologous genes in all four brassica species. One notable example was a CRM composed of TFBSs from bZIP, MYB-related, NAC, and Trihelix families located upstream of the *A. thaliana* gene RTFL15 (**Fig. 5E**). RTFL15 is a member of the highly conserved DEVIL/ROTUNDIFOLIA-LIKE (DVL/RTFL) gene family involved in plant organogenesis^51,52^. Other highly conserved CRM-associated genes include central regulators of plant development (**Extended Data Fig. 9B**). For example, TREHALOSE-6-PHOSPHATE SYNTHASE 1 (TPS1), the primary enzyme catalyzing trehalose-6-phosphate synthesis in *A. thaliana*, functions as a sucrose-responsive signal linking carbon status with developmental progression^53^.

We next used Gene Ontology (GO) enrichment analysis to characterize the functional roles of these distal CRMs. We found that the 1,736 genes associated with *A. thaliana* distal CRMs were significantly enriched for developmental plant processes including response to auxin (adjusted p = 4.54×10^-25^), plant organ development (adjusted p = 6.86×10^-15^), and floral organ development (adjusted p = 3.81×10^-17^) (**Fig. 5F**). These distal-enriched functions aligned closely with genomic signatures typically associated with enhancers^51,52^ and often differed from the functions enriched among genes harboring c4 promoter TFBSs from the same TF (**Fig. 5F**). Despite these functional differences, 80% of TFs with at least 50 distal CRM target genes showed higher overlap between distal CRM target genes and c4 promoter target genes than expected by chance (adjusted *p* < 1.61×10^-55^, **Supplementary Table 4**). While this overlap was significant, it was still relatively small in magnitude (20.2%), explaining the differences in functional enrichments. Together, this pattern points to TFs associated with distal CRMs acting across partially intersecting but largely partitioned regulatory programs, and aligns with enhancer-like activity for many of the identified distal CRMs^51,52^.

To understand which aspects of plant development and hormone biology are governed by distal CRMs, we analyzed pathway and gene-level enrichment within CRM target sets. Core auxin pathway regulators are strongly overrepresented among CRM target genes, particularly ARF and AUX/IAA transcriptional regulators (ARF Odds Ratio (OR) =10.13, p = 6.9x10⁻⁷; AUX/IAA OR = 11.85, p = 1.56x10⁻^9^, **Supplementary Table 5**). CRM enrichment was concentrated in the core developmental ARFs: four of the five Class A ARFs (ARF5/6/8/19)(**Supplementary Table 6**), TFs central to organogenesis and cell fate specification, together with major patterning ARFs (ARF3/10/16), show strong CRM associations. Additional enrichment for GH3 auxin-conjugating enzymes (OR = 6.43, p = 5.66x10⁻⁴) and SAUR growth-activating genes (OR = 2.81, p = 7.05x10⁻⁴) indicates that CRMs extend beyond core signaling circuitry to both upstream and downstream physiological effectors that shape the auxin response^53^.

We observed that many auxin pathway genes targeted by CRMs exhibited markedly expanded upstream intergenic regions in *A. thaliana*. For example, CRM-bearing AUX/IAA and ARF genes have upstream intergenic regions approximately 7 to 8-fold larger than the genome-wide median and approximately 4-fold larger than non-CRM members of the same families. These expanded intergenic regions coincide with elevated CRM counts at developmental regulatory loci, consistent with the possibility that increased intergenic noncoding space provides the capacity to accommodate additional cis-regulatory content required for more complex regulatory programs. This pattern may be easier to detect in compact genomes such as *A. thaliana* than in larger genomes such as maize, where the larger baseline intergenic space may obscure such a signal. We observed similar functionally-enriched CRMs in atypically large intergenic regions for key regulators of other hormones, including gibberellin (GA) biosynthesis (OR = 14.19, p = 1.27x10^-6^) and GA responses (OR = 8.29, p = 9.3x10^-7^), and cytokinin response regulators^54^ (OR = 5.52, p=5.0x10^-4^, **Supplementary Table 5 and 7**). Across pathways, TF-encoding genes are broadly overrepresented among CRM targets (OR = 4.47; p = 1.08x10^-115^), supporting the conclusion that conserved distal CRMs preferentially target developmental and regulatory control hubs.

Using our snATAC-seq dataset, we evaluated chromatin accessibility at CRMs linked to auxin, GA, and cytokinin genes. Specifically, we quantified the cell type-specificity (tau score) of the DARs intersecting these CRMs. Permutation testing revealed that CRMs associated with target genes for all three hormone pathways show significantly higher cell type specificity than the global genomic background. This enrichment toward localized accessibility is robustly observed for 102 auxin-associated genes (mean tau = 0.58 vs. background mean = 0.53; Cohen’s *d* = 0.41, permutation *p* = 0.0001) and 36 GA-associated genes (mean tau = 0.60; Cohen’s *d* = 0.54, permutation *p* = 0.0004), with a modest but significant shift also observed for 40 cytokinin targets (mean tau = 0.57; Cohen’s *d* = 0.29, permutation *p* = 0.034). Consistent with this localized accessibility profile, these hormone-related CRMs display highly enriched chromatin accessibility within specialized vascular lineages, specifically in elongating xylem and phloem parenchyma cells for auxin and cytokinin and phloem parenchyma for GA.

To prioritize high-confidence distal CRM candidates, we progressively filtered the full catalog of 3,333 *A. thaliana* distal CRMs. First, we identified 740 CRMs located within cell type-specific ACRs (Tau > 0.5) (**Supplementary Table 8**). We observed strong correlations between cell type enrichment scores of the same TFs (c4 TFBSs) in promoters and distal CRMs (**Fig. 5G and Fig. 5H, Extended Data Fig. 9C**), suggesting that distal CRMs can influence cell type-specific promoter chromatin accessibility consistent with enhancer activity. ACRs overlapping distal CRMs or promoter c4 TFBSs showed significantly higher cell type specificity than distal ACRs, distal c4 TFBSs, or promoter ACRs in general (**Fig. 5I**). Furthermore, we noted significantly stronger correlations between per-cell type gene expression and per-cell type accessibility of ACRs overlapping distal CRMs compared to other distal ACRs (1.7 fold change, p < 2.2x10^-16^), suggesting that CRMs play a role regulating cell type-specific gene expression (**Fig. 5J)**. To pinpoint CRMs with the strongest evidence of enhancer function, we focused on distal CRMs that showed strong 1) cell type-specific accessibility (Tau > 0.5), and 2) a match between the cell type with highest CRM accessibility and the cell type with highest expression of the associated gene. In total, we identified 102 high-confidence distal CRM candidates that met these stringent matched accessibility/expression criteria, with top accessibility/expression in 14 distinct cell types (**Extended Data Fig. 9D, Supplementary Table 9**). For example, we found a CRM composed of six TF families upstream of LAC5 (**Fig. 5K**), a gene most strongly expressed in suberized endodermis, located in chromatin that was specifically accessible in suberized endodermis. LAC5 is known to accumulate in the endodermis and is a key component of the lignin biosynthesis pathway, upstream of suberin biosynthesis^58,59^. Another example comprised a CRM accessible predominantly in phloem parenchyma cells upstream of SIAR1, a gene specifically expressed in the same cell type (**Fig. 5l**) where it has a known role in amino acid transport^60,61^. The combination of distal positioning, cell type-specific accessibility matching target gene expression patterns, and enrichment for developmental gene functions collectively support that these CRMs are strong candidates for plant developmental enhancers.

## Discussion

A central challenge in plant regulatory genomics is to move from catalogs of transcription factor binding events to a framework that explains how regulatory information is organized around genes. Our results support TFBS position as a major organizing principle. By integrating conserved multiDAP maps for 244 TFs with single-nucleus chromatin accessibility, cell type-resolved expression, and abiotic and biotic stress-response transcriptomics across Brassicaceae, we show that regulatory output is not determined by TF identity or motif presence alone. Instead, TFBS position and chromatin context appear to partition regulatory output into promoter-proximal stress-response platforms, distal and intronic developmental regulatory regions, and coding-sequence contexts potentially associated with constrained or repressive regulation.

Recent massively parallel reporter assay (MPRA) work showed that plant regulatory sequence activity can depend strongly on position of regulatory sequence relative to the TSS, providing an important functional precedent for position-dependent regulation^1^. However, because the MPRAs used transient, non-integrated reporter constructs in bulk populations of isolated protoplasts, they did not capture endogenous chromatin effects, lacked cell type resolution, and may not fully reflect native cellular behavior. Thus, although this work revealed that TFBS position can shape regulatory output, it remained unclear how positional dependence operates within native plant regulatory architecture. Here, we address this gap by integrating conserved TFBS location with chromatin accessibility, cell type expression, and hormone-response output, showing that position helps partition developmental, stress-responsive, and candidate repressive regulatory programs in a highly TF-specific manner (**Fig. 6**).

**Fig. 6.**
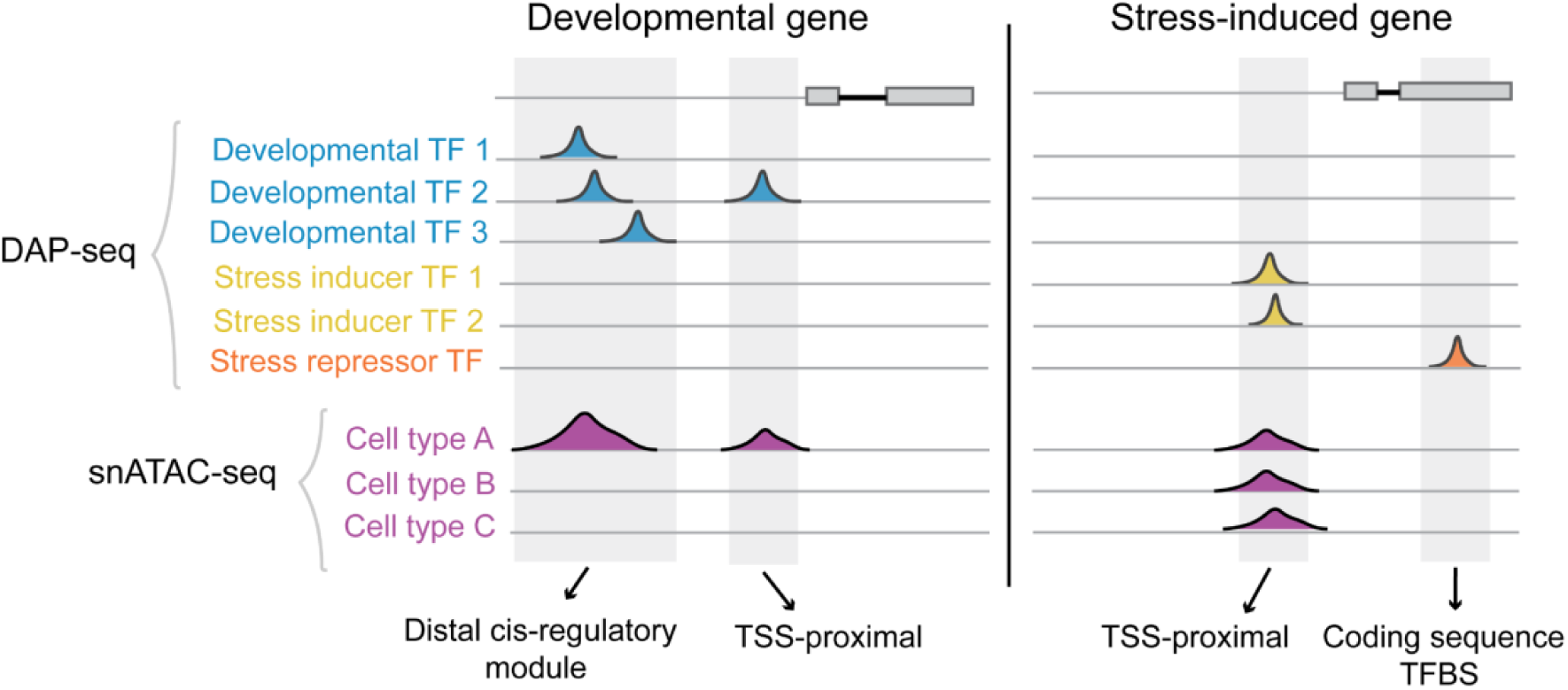
Positional partitioning of developmental and stress-responsive plant gene regulation. Summary model illustrating how TFBS location and chromatin context differentiate regulatory output. Cellular identity programs (developmental cell type-specific regulation) tend to be driven by combinatorial TF binding across both cell type-restricted TSS sites and distal, enhancer-like regulatory modules. Conversely, environmental response networks (stress regulation) leverage globally accessible TSS-proximal platforms to coordinate rapid, widespread transcriptional activation upon stress-induced TF expression.

The first mode is centered on the TSS-proximal region. Conserved TFBSs are enriched near the TSS, and these regions are broadly accessible across cell types. Yet TSS-proximal accessibility is weakly correlated with cell type-specific expression, suggesting that promoter openness often reflects transcriptional competence rather than cell identity. Hormone-response data clarify one major role of this region: stress-associated TF families, including bZIP, WRKY, NAC, and AP2/EREBP factors, are preferentially associated with abiotic and biotic hormone-induced expression through TSS-proximal sites. Constitutively accessible promoters may therefore provide preconfigured platforms through which changes in TF abundance or activity rapidly alter transcription, potentially without requiring extensive chromatin remodeling. This extends stress cis-regulatory code models and promoter-proximal stress TF binding examples^9,10,43,44,62^ by linking TF family identity, conserved position, chromatin state, and response output in one framework.

The second mode involves distal promoter and intronic regions that support developmental and cell type-specific regulation. In contrast to broadly accessible TSS-proximal regions, cell type-specific expression is most strongly associated with conserved TFBSs embedded in cell type-restricted accessible chromatin. These sites occur in TF family-specific patterns, suggesting that different families occupy characteristic regulatory territories rather than a universal preferred position. This places known examples of distal plant regulation, including TB1, Vgt1, KRN4/UB3, and FT, into a broader genome-scale model, and complements single-cell chromatin studies linking cell type-specific ACRs to enhancer activity, developmental state, and trait variation^2,5,21,22,63–71^. The advance is not simply that distal enhancers exist, but that conserved, family-specific TFBS positions embedded in restricted chromatin help explain developmental regulatory output.

A prominent subset of these distal elements consists of conserved multi-family CRMs with enhancer-like features. These CRMs contain heterotypic clusters of conserved TFBSs from multiple families, including Homeobox and other well-known developmental regulators, overlap rare cell type-specific accessible chromatin, and are enriched near genes involved in embryonic patterning, meristem function, auxin signaling, and other hormone-dependent developmental programs. Their association with ARF, AUX/IAA, GH3, SAUR, and transcription factor-encoding genes suggests that they mark developmental control hubs rather than generic distal noncoding space. Because canonical enhancer signatures such as H3K4me1 enrichment or eRNA production have not been uniformly predictive in plants^50,72^, the combination of conservation, heterotypic TFBS clustering, cell type-specific accessibility, and developmental gene association provides a practical route for prioritizing plant enhancer-like elements.

Many CRM-associated developmental genes also have unusually expanded upstream intergenic regions. In the compact Arabidopsis genome, this suggests that some developmental regulatory hubs may retain larger noncoding intervals to accommodate combinatorial TF inputs, although local genome organization and gene annotation effects could also contribute. This interpretation remains speculative, but it provides a useful explanation for why conserved heterotypic CRMs and expanded upstream regions are conspicuous at these loci. While the integration of evolutionary conservation, chromatin accessibility, and cell type-specific expression provides robust computational evidence for these distal CRMs, future *in vivo* functional studies, such as transgenic reporter assays, will be required to definitively validate their enhancer activity.

The third, more tentative mode is coding sequence, where TFBSs lie within protein-coding regions and the same DNA bases may encode both amino acids and regulatory information^73^. CDS-localized transcriptional regulation remains largely underexplored at the genome scale in plants, although prior work has shown that coding-region TFBSs in photosynthetic genes can shape spatial expression by repressing transcription in mesophyll cells, and protoplast TF perturbation experiments have linked individual CDS-localized DAP-seq sites with repressive effects^48,74^. Separately, yeast coding regions harbor latent promoter-like sequences that can drive antisense transcription when chromatin-based suppression is lost^49^. Our results identify a genome-scale candidate example of CDS-localized TFBS-associated repression: target genes with CDS-only ABR1 sites show strong SA-associated repression. Together with prior evidence for dual-use codons and plant duons, the ABR1 pattern suggests that coding-region TFBSs represent a broader, underrecognized regulatory layer in plants.

Together, these findings support a positional grammar for plant gene regulation: TSS-proximal sites provide broadly accessible platforms for rapid inducible responses; distal and intronic TFBSs in cell type-restricted chromatin support cell type-specific developmental regulation, with conserved heterotypic CRMs representing an enhancer-like subset; and coding-region TFBSs mark a potential context for repressive regulation. Although DAP-seq reports *in vitro* binding potential rather than *in vivo* occupancy, the TFBS positional patterns identified here are filtered by cross-species conservation and supported by integrated evidence from single-nucleus chromatin accessibility and expression, hormone-response transcriptomics, and TF family identity. This convergence supports a model in which TFBS position contributes to regulatory specificity. The resulting framework provides a basis for interpreting cis-regulatory variation and regulatory evolution, and may help guide synthetic plant regulatory design.

## Methods

### Genomic feature definition and extraction

We extracted coding sequence (CDS) annotations from genome annotation GFF files for all ten plant species analyzed (1_combine_cds.ipynb). For each gene, we merged individual CDS features to generate a single genomic interval spanning from the start of the first CDS to the end of the last CDS for each transcript. We defined promoter and distal regulatory regions relative to CDS boundaries using strand-aware calculations (2_intergenic_dist.ipynb and 3_extract_features.ipynb). For each gene, we calculated intergenic distances to neighboring genes based on genomic position and strand orientation. For genes on the positive strand, we defined promoter regions as extending 2,000 bp upstream of the CDS start (or to the end of the upstream gene, whichever was closer) and 500 bp into the gene body. For genes on the negative strand, we extended promoters 2,000 bp downstream of the CDS end (or to the start of the downstream gene) and 500 bp into the gene body.

We defined distal regulatory regions only for genes with intergenic distances >2,000 bp to the neighboring gene. For positive-strand genes, we extended distal regions from 10,000 bp upstream of the CDS start (or the end of the upstream gene) to the start of the promoter region. For negative-strand genes, we extended distal regions from the end of the promoter region to 10,000 bp downstream of the CDS end (or the start of the downstream gene). We implemented two approaches for refining distal regions to prevent overlap with promoter regions. Script 2_intergenic_dist.ipynb employed a neighbor-based adjustment method where we iteratively checked each distal region for overlap with adjacent gene promoters and CDS features, adjusting region boundaries accordingly. For plus-strand genes, if the distal region overlapped with the upstream gene’s promoter or CDS, we shifted the distal start coordinate to avoid the overlap. For minus-strand genes, we adjusted the distal end coordinate to prevent overlap with the downstream gene’s promoter or CDS. We excluded distal regions that became invalid after adjustment (start ≥ end or start < 0) from analysis. Additionally, we subtracted all promoter regions from distal regions genome-wide. When this subtraction process fragmented distal regions into multiple non-overlapping windows, we retained the largest contiguous segment for each gene and recorded the total number of windows for quality control. We excluded genes with overlapping CDS features from all regulatory region analyses to avoid ambiguous assignments.

### DAP-seq Analysis

#### DAP-seq peak identification

Fastq files were quality filtered and adapters were trimmed using BBTools v38.96 bbduk.sh^75^ with parameters k=21 mink=11 ktrim=r tbo tpe qtrim=r trimq=6 maq=10, and then aligned to the corresponding reference genome using bowtie2 v2.4.2^76^ and parameters --no-mixed --no-discordant. Bam files from negative control wells were merged using the samtools v1.15.1^77^ merge command, to generate a single background file for each 96-well plate. The MACS3 v3.0.0a6^78^ callpeak command was used to generate narrowPeak files, using the merged background as the control, with the parameters --call-summits --keep-dup 1 and –gsize with the total fasta genome size. Motifs were called using MEME suite v5.3.0^79^ command meme with parameters -dna -revcomp -mod anr -nmotifs 2 -minw 8 -maxw 32, and with a background model (-bfile) generated from the entire reference genome file with fasta-get-markov -m 0 -dna.

The resulting datasets were filtered to exclude those TFs and replicates with poor performance in the DAP-seq assay. As a measure of successful binding site enrichment, we calculated the fraction of reads in peaks (FRIP) scores and only used data from DAP-seq assays that generated FRIP scores of at least 0.05. For experiments including multiplexed species, we only considered those assays where all of the multiplexed species produced FRIP scores of 0.05 or greater. We found that multiplexing larger sets of species with varying genome sizes in a single multiDAP experiment sometimes resulted in large differences in read coverage between different TFs and in some cases differential enrichment of species. To mitigate the effect of low coverage resulting in false negatives in the peak-calling step (and thus lowering apparent c scores), we only used datasets that yielded at least 200,000 read pairs (400,000 reads) for each of the multiplexed species.

#### Peak Assignment to Genomic Features

We assigned DAP-seq peaks to genomic features using a multi-step approach (1_assign_peaks.sh). We first assigned peaks to the five nearest mRNA features, as well as to overlapping exon and CDS features using BEDTools^80^ closest v2.31.0 with default parameters. We intersected peak summits with promoter regions and distal regulatory regions using BEDTools intersect with the -f 1 flag to require complete overlap of the summit position.

We then annotated assigned peaks with detailed genomic context information using a custom Python script (2_annotate_peaks.py). This annotation workflow calculated strand-aware distances from peak summits to transcript features, including distance to transcription start sites (TSS), transcription end sites, and CDS boundaries. We classified peaks into genomic regions (promoter, distal, CDS, 5’ UTR, 3’ UTR, intron, upstream, or downstream) based on summit position relative to gene features and regulatory regions. To ensure mutual exclusivity between promoter and distal assignments, we retained peaks assigned to both categories only in the promoter category and removed them from distal assignments.

#### Peak filtering and conservation scores (c score)

We filtered peaks based on fold-change enrichment (minimum threshold of 5-fold) to retain high-confidence binding events (3_filter_mark_cons_n4.ipynb). To assess binding conservation across species, we grouped genes into orthogroups using OrthoFinder v2.5.5^81^ results and excluded organellar genes from conservation analyses. For each transcription factor and species, we calculated the fraction of genes within each orthogroup that the TF bound, enabling comparison of TF-gene regulatory relationships across the four brassica species we studied.

Finally, a conservation score (c score) was assigned to each TF-orthogroup assignment, representing a count of species in which the given TF also targets at least one gene within the same orthogroup. While c4 sites are shared by all species including *A. thaliana*, c1 sites represent binding events unique to any one of the four species. See code availability statement for scripts.

#### Distance-based TFBS density calculation and normalization

We calculated normalized transcription factor binding site (TFBS) density across promoter regions using two reference points: the transcription start site (TSS) and the translation start site (ATG) (density_tss_50bp.ipynb and density_atg_50bp.ipynb). For each approach, we divided promoter regions into 50 bp bins and counted DAP-seq peak summits within each bin. We calculated raw TFBS density as the number of peak summits per bin divided by the total number of peaks analyzed for each TF. To account for background binding preferences, we generated a matched null distribution using random genomic regions. We created random peaks matching the size distribution of actual DAP-seq peaks and calculated their density across the same 50 bp bins in promoter regions. For each TF and each bin, we normalized the observed density by dividing it by the background density, producing fold-enrichment values that account for positional biases in the genome. We computed these normalized densities separately for TSS-relative coordinates (measuring distance from transcription start) and ATG-relative coordinates (measuring distance from translation start), enabling comparison of TF binding patterns relative to both transcriptional and translational regulatory elements.

#### Gini Coefficient Calculation for Binding Specificity

We quantified the evenness of TF binding distributions across target genes using Gini coefficients. For each TF, we calculated the number of binding sites per target gene and computed the Gini coefficient, which ranges from 0 (perfectly even distribution, where all genes have equal numbers of binding sites) to 1 (maximum inequality, where binding is concentrated in very few genes). We used the standard Gini coefficient formula based on the Lorenz curve, ordering genes by their binding site counts and calculating the area between the line of equality and the actual cumulative distribution. This metric enabled us to distinguish between TFs that bind broadly across many targets with similar intensities versus those that show concentrated binding at specific high-affinity targets, providing insights into different modes of transcriptional regulation.

#### Feature-based TF density calculations and assignments

Using our dataset of filtered c4 promoter-region (2kb upstream of the start codon to 500 bp downstream of the start codon) peaks, we calculated the density of TFBSs for each TF in each of five gene-based features: upstream (more than 200 bp upstream of the TSS), TSS (within 200 bp upstream of the TSS), 5’UTR (between the TSS and the start codon, in an exon), coding sequence, and intron. Densities were calculated by assigning each peak’s summit to one of the features, summing peaks assigned to each feature, and normalizing by both the total feature’s sequence length in our defined promoter regions, as well as the total number of peaks for the TF. To assign each TF to a feature-based neighborhood (FBN), we assigned each TF a z-score for each feature, representing TFBS density relative to other TFs. TFs with at least one z-score higher than 0.5 were assigned to the feature with the highest z-score, and other TFs were assigned to the ‘even’ group as they showed little skew toward any specific feature.

### Epigenomic mark enrichment test

To evaluate the association between TF binding and epigenomic mark occurrence, published ChIP-seq data for several histone marks as well as Methyl-seq and ATAC-seq was intersected with the multiDAP data. For histone marks and ATAC-seq, BED files of called peaks were obtained from their respective studies on the NCBI GEO and SRA repositories. Methylated regions were defined from whole genome bisulfite data using a 50 bp sliding window^82,83^; windows containing three or more methylated cytosines were called methylated regions. These methylated regions were then used as BED intervals for the epigenomic mark enrichment test, analogous to ChIP-seq peak and ATAC-seq peak BED files. Seedling ATAC-seq data were from Ref. ^12^ (GSE85203). Histone modification ChIP-seq datasets were obtained from the following sources: flower H3K36ac (SRP066007^14^), leaf H3K27ac (SRP044176^51^), seedling H3K4me2, H3K4me3, and H3K9me2 (DRA005154^16^), leaf H3K9ac (ERP122722^84^), seedling H3K27me3 (ERP005526^85^), and seedling H3K36me3 (SRP049385^15^). DNA methylation data (GSM1085222) was obtained from the 1001 Epigenomes project^83^. For each TF clade, the peaks in each conservation group are downsized to 10 bp flanking peak summit, then the proportion of DAP-seq peaks in each conservation group that overlap with individual epigenomic marks was calculated. To eliminate the background, we used a simulated peaks list which was produced by randomly sampling peaks from -2000 bp to +500 bp region around TSS as the DAP-seq peaks were restricted to. For each TF clade, the same number of simulated peaks were used to get the proportion of peaks overlapping with epigenomic marks. The enrichment of TF and epigenomic mark co-occurrence was evaluated by the ratio of two proportions.

### Single nuclei ATAC-seq

#### Plant growth conditions

*A. thaliana*, *A. lyrata*, *C. rubella*, and *B. oleracea* seeds were surface sterilized before planting vertically on agar plates containing 3mM Ca(NO_3_)_2_, 1.5mM MgSO_4_, 1.25mM NH_4_H_2_PO_4_, 1mM KCl, and 1x micronutrients (Murashige and Skoog Micronutrient Salts 100x, MSP18-10LT), in 0.8% agar with pH 5.7. The seeds were sterilized by soaking in 75% ethanol for 1 minute and 50% bleach for 5 minutes, 25% bleach with 0.2% Triton X-100 (Sigma-Aldrich 93443) for 4 minutes, then rinsed with sterile water 8 times. After sterilization, seeds in the 2 mL Eppendorf tube were wrapped in aluminum foil and put into a 4°C fridge for cold stratification. Seeds were left in the fridge for 3 days for *A. thaliana* and *C. rubella*, or 5 days for *A. lyrata*. Cold stratification was not needed for efficient germination of *B. oleracea* seeds. We staggered the planting of different species to account for different germination times of the four plant species, allowing us to harvest seedling root and shoot tissues of plants at the same 5-7 day old growth stage of all species on the same day. *A. lyrata* was planted on the agar plates 3 days before *A. thaliana*, *C. rubella* was planted 1 day after *A. thaliana*, and *B. oleracea* seeds without cold stratification were planted 2 days after *A. thaliana*. The conditions of the growth chamber (Percival Scientific) were set to 16 hours light at 22°C and 8 hours dark at 19°C, with the light turning on at 5am and turning off at 9pm. Root, shoot, and whole seedlings were sampled between 10am and 12pm for all experiments.

In order to enable sampling of mature leaf and flower bud samples on the same day of all four species for snATAC-seq experiments, planting times were staggered and growth conditions were modulated to synchronize growth of all four species. As a result, the sampled tissues represent approximate rather than exact developmental equivalents. For mature leaves, we sampled the youngest fully expanded rosette leaf from plants grown in soil in a plant grow room or cold room. *B. oleracea* mature leaves were harvested from plants that were two months old grown at 13°C under a light cycle of 12 hours light/12 hours dark. *A. lyrata* was three months old at harvest, grown at 4°C with a light cycle of 12 hours light/12 hours dark. *A. thaliana* and *C. rubella* were grown in conditions of 16 hours light at 28°C and 8 hours dark at 23°C and harvested at 2 months old. Mature leaves were sampled between 10am and 12pm.

For floral buds, we sampled young buds on the main stem from plants grown in soil, where the petal was just emerging (∼1 mm visible). All four plant species were grown at 16 hours light at 22°C and 8 hours dark at 19°C when floral buds were collected. *A. thaliana* and *B. oleracea* flowered at about two months old without cold treatment. To induce flowering in *A. lyrata* and *C. rubella*, two month old plants were transferred into a 4°C cold room for two months and then transferred back to conditions with 16 hours light at 22°C and 8 hours dark at 19°C for one month. In this way, all the four plant species were synchronized to flower on the same day and sampled between 10am-12pm.

#### Hormone treatment experiments

*A. thaliana* seeds were sterilized and cold stratified as described above, then planted on nylon mesh in agar plates. Seeds were sown on plates at a density of 3 rows per plate with ∼30 plants per row and plates were placed vertically in a growth chamber under conditions of 16 h light at 22°C, 8 h dark at 19°C for 5 days. On the 5th day, the nylon mesh and plants were transferred using forceps to agar plates containing either 2 μM ABA, 150 μM SA, or the standard growth medium without added hormones as a negative control. Samples were collected at time points of 0, 1, 2, 4, 7, 24, and 48 hours, separating root and shoot tissues with a single razor blade cut at the time of collection. At each time point, five biological replicates per treatment were collected and flash frozen in liquid nitrogen then stored at -80°C. Three biological replicates per treatment were used for bulk RNA-seq.

#### Nuclei isolation and single nuclei library preparation

Freshly harvested samples were used for all snATAC-seq libraries. Nuclei isolation from tissues was performed as follows. Buffer 1 (lysis buffer) consisted of 0.275 M sorbitol (Sigma-Aldrich S6021), 0.1% Triton X-100 (Sigma-Aldrich 93443), 0.01 M MgCl_2_ (Ambion AM9530G), 1x protease inhibitor cocktail (Sigma-Aldrich 4693132001), 0.3 U/μl RNase inhibitor (Roche 03335399001), and 1 mM DTT (Teknova D9750). Buffer 2 (wash and resuspension buffer) consisted of 1x phosphate buffer saline (without Mg and Ca), 1% BSA, 0.4 U/μl RNase inhibitor, and 1 mM DTT. For isolation of nuclei from mature leaves containing high levels of chloroplasts, an extra modified wash and resuspension buffer (buffer 3) was used to eliminate chloroplasts before the final wash and resuspension step by buffer 2. It consisted of 0.275 M sorbitol, 0.2% Triton X-100, 0.2% NP40 (Thermo Fisher Scientific PI28324), 0.01 M MgCl_2_, 1x protease inhibitor cocktail, and 1 mM DTT.

All nuclei isolation steps were carried out in a cold room at 4°C. For each sample, 20-100 mg fresh or frozen tissue was chopped for 3 minutes using a razor blade on a glass plate with 100-200 μl buffer 1, then transferred to a petri dish with 1.5 mL cold buffer 1 and allowed to rest on ice with gentle shaking for 2 minutes. For multiplexed samples, tissues from different species were weighed separately before being combined and chopped together. The samples were then transferred to a 48-well filter plate (25 μm, 4mL, Agilent, 201003-100), which was pre-wet with 1 mL cold buffer 1 before it was placed on the receiving plate attached to a QIAvac96 (Qiagen). Pressure was maintained below 100 bar during filtration. The filtered nuclei solution was centrifuged at 500x g for 10 minutes at 4°C to pellet the nuclei, after which the supernatant was discarded.

For samples with low or no chloroplast content: The pellet was resuspended by gentle flicking and pipetting, washed with 2 mL cold buffer 2, centrifuged at 500x g for 5 minutes, and the supernatant was discarded. For nuclei isolation from mature leaves with high chloroplast content: The pellet was resuspended by gentle flicking and pipetting and 10 mL cold buffer 3 was added, centrifuged at 500x g for 5 minutes, and the supernatant was discarded. The pellet was washed with 2 mL cold buffer 2, centrifuged at 500x g for 10 minutes, and the supernatant was discarded. For both protocols, pellets from the final centrifugation were resuspended in 100 μl cold buffer 2 for snRNA-seq, or cold 1x nuclei buffer (10X Genomics PN-2000207) for snATAC-seq.

Nuclei quality and quantity were assessed by flow cytometry and microscopy before continuing to the single nuclei partitioning and barcoding step. A BD Accuri C6 plus flow cytometer was used to analyze the quality and quantity of nuclei stained by propidium iodide (Sigma-Aldrich P4864), added to a final concentration of 50 μg/mL. The stained samples were incubated on ice in darkness for 5-10 min prior to analysis. The analysis was conducted using light-scatter and fluorescence signals produced by a 20 mW laser illumination at 488 nm. Signals corresponding to forward-angle scatter (FSC), 90° side scatter (SSC), and fluorescence were collected.

Fluorescence signals (pulse area measurements) were screened using the following filter configurations: (a) FL-2, with a 585/40 nm band-pass filter, and (b) FL-3, with a 670 nm long-pass filter. Threshold levels were set empirically at 80,000 for FSC-H to exclude debris commonly found in plant homogenates. Templates for uni- (FL2-A and number of nuclei) and bi-parametric (FL2-A and FL3-A) frequency distributions were established. Upon identifying the region corresponding to nuclei, data was collected to a total count of 5,000–10,000 nuclei. The flow cytometer was operated at the Slow Flow Rate setting (14 μl sample/minute), with data acquisition for a single sample typically taking 3–5 minutes. Different plant species exhibited varying patterns of somatic endoreduplication within most of their tissues and organs. For example, in young *A. thaliana* roots and cotyledons, this variation is reflected in the form of multiple clusters within the distribution, corresponding to nuclei forming a 2C, 4C, 8C, 16C, etc., of the endoreduplicative series^86^. All the nuclei of all the endoreduplicative series were counted and divided by the actual volume analyzed in the flow cytometer to obtain the concentration of the nuclei. An EVOS M5000 microscope was also used for validation of nuclei quality and quantity by analyzing 7 μl diluted nuclei sample stained by 1 µl of diluted SYBR Green dye (Thermo Fisher Scientific S7585 diluted 1:10,000 in water) using a C-Chip Disposable Hemocytometer. For nuclei prepared from fresh tissue samples, most of the nuclei in each sample showed well-resolved edges without obvious evidence of blebbing. In the case of nuclei prepared from frozen samples, where cell debris around the nuclei can interfere with these microscopy observations, nuclei quality was primarily assessed by comparing to fresh nuclei’s distribution and the endoreduplicative series pattern on the flow cytometer.

The nuclei suspension was then diluted to concentration of 700-1000 nuclei/μl using cold buffer 2 for snRNAseq or cold 1x nuclei buffer (10x Genomics PN-2000207) for snATAC-seq. Single nuclei were partitioned and barcoded using a Chromium system (10x Genomics) with the Chromium Next GEM Single Cell 3’ Reagent Kits v3.1 (Dual Index) for snRNAseq or Chromium Single Cell ATAC Reagent Kits v2 for snATAC-seq. Approximately 20,000 nuclei in total were loaded for single species samples, and up to 60,000 nuclei for multiplexed species samples. Library creation was carried out following the manufacturer’s protocol. Sequencing was performed on an Illumina NovaSeq 6000 S4 flow cell or NovaSeq X plus 10B or 25B flow cell, using 2x150 bp paired-end sequencing, targeting 200M fragments per species for RNA-seq samples and 250M fragments per species for mixed-species ATAC-seq samples, with higher coverage for *A. thaliana* seedling samples (900M fragments and 1800M fragments for root and shoot respectively).

### Single Nuclei ATAC-seq Analysis

#### snATAC-seq atlas construction

We used the Cell Ranger-ATAC version 2.1.0^87^ count command to filter and map reads to each reference, and generate the fragments file containing a unique ATAC-seq fragment created by two separate transposition events. We then used SnapATAC2 v2.50^88^ to process the fragment files to downstream analyses from cell filtering to cell type annotation. To keep high-quality cells we required a minimum of 3000 fragments per cell and a 3-fold enrichment around the TSS. We separated the data into 500 bp windows across the genome and created a cell by bin count matrix. We used the scrublet^89^ algorithm to identify potential doublets and remove them afterwards. We then combined replicates (**Supplementary Table 2**) using the MNN-Correct algorithm in SnapATAC2 for batch correction. For visualization purposes we created a UMAP of each atlas. Pseudo-bulk profiles generated from our snATAC-seq data aligned well with published^12^ whole-tissue open chromatin profiles (Pearson r2 0.58).

We used gene accessibility scores as a proxy for gene expression to annotate cells with cell types from our snRNA-seq atlases. We calculated gene accessibility scores using the ‘pp.make_gene_matrix()’ function in snapATAC2. Briefly, this function assigns genomic bins to nearby genes, counts the transposase insertion events associated with each gene, and creates a matrix of accessibility scores for a particular gene and cell. Using the gene accessibility matrix, we integrated the snATAC-seq and the snRNA-seq datasets selecting the top 1000 most variable genes from our snATAC-seq data. We used scVI^90^ integrated in SnapATAC2 to transfer cell type labels from the snRNA-seq atlas to the snATAC-seq atlas. To do this, we trained models using as covariate the batch information, negative binomial distribution (“nb”) as the gene likelihood statistic, and a maximum number of epochs of 1000. After assigning cell type labels, we called peaks using MACS3^78^ integrated into snapATAC2 independently for each cell type. We then merged peaks that were in close proximity (250 bp to either edge) to make comparisons across different cell types possible. To calculate the differentially accessible regions (DARs) we sampled random cells per cell type, excluding the cell type being tested, and combined them to create a background using the regression-based test from snapATAC2 “snap.tl.diff_test()”, to identify regions more likely to be open in a given cell type than the background set of cells. We defined DARs as peaks with an adjusted p< 0.01.

#### DAP-seq and snATAC-seq peak overlap analysis

We integrated DAP-seq TF binding data with single-nucleus ATAC-seq (snATAC-seq) chromatin accessibility profiles from *Arabidopsis thaliana* seedlings (dap_vs_DA_peaks_to_boolean_matrix.ipynb). We first filtered the snATAC-seq dataset to retain only peaks located within promoter regions (defined as regions within gene regulatory boundaries). We then intersected DAP-seq peak summits with these promoter-localized snATAC-seq accessible chromatin peaks using BEDTools. For each transcription factor, we created a binary matrix indicating which snATAC-seq peaks overlapped with DAP-seq binding sites. This produced a TF-by-peak boolean matrix where TRUE values indicated overlap between TF binding and accessible chromatin, enabling systematic comparison of predicted TF binding sites with experimentally accessible regulatory regions.

#### Normalized chromatin accessibility and cell type specificity values

We calculated normalized chromatin accessibility values for each TF across cell types using promoter-localized peaks (c4s_promoter_normalized_accessibility_and_tau_by_TF.ipynb). We first intersected DAP-seq peaks with snATAC-seq data to identify overlapping accessible regions in promoters. For peaks that did not intersect with snATAC-seq accessible regions, we assigned a mean accessibility value of 0. For peaks that did intersect, we handled cases where a single DAP-seq peak overlapped multiple snATAC-seq peaks by taking the mean accessibility across all overlapping regions.

To quantify cell type specificity, we calculated tau (τ) scores for each accessible region using the formula: τ = Σ(1 - x_i_/x_max_)/(n-1), where x_i_ represents accessibility in cell type i, x_max_ is the maximum accessibility across all cell types, and n is the number of cell types. We computed relative tau scores (tau fold-change values) by normalizing against the median tau score within mean accessibility bins to account for the relationship between overall accessibility and specificity. We retained only peaks with maximum accessibility ≥ 0.1 across cell types to focus on robustly accessible regions.

We then computed TF-specific normalized accessibility values by aggregating across all DAP-seq peaks for each TF. To assess statistical significance, we generated a background distribution by creating more than 35,000 random genomic regions matching the size distribution of DAP-seq peaks. We calculated mean accessibility for these random regions and performed bootstrap sampling (1,000 iterations) to generate TF-specific null distributions, sampling with replacement to match each TF’s peak count. We computed empirical p-values by comparing observed TF accessibility to these bootstrapped backgrounds and applied Benjamini-Hochberg false discovery rate correction (α = 0.05).

#### snATAC-based TF activity scoring and cell type enrichment

We calculated snATAC-based TF activity scores for each cell using a module score approach (tf_module_score_and_enrichment.ipynb). For each TF, we identified all snATAC-seq peaks that overlapped with its DAP-seq binding sites to define a TF-specific peak module. We computed module scores by averaging accessibility values across module peaks for each cell and subtracting a matched control score. We selected control peaks from the same mean accessibility bins as module peaks (using 25 bins based on genome-wide accessibility percentiles) to account for technical biases. We dynamically determined the number of control peaks as the minimum of 5,000 or the maximum of 50 and the number of module peaks, with a default control-to-module ratio of 1:1.

To identify cell type-specific TF activities, we performed differential activity testing using the Wilcoxon rank-sum test, comparing module scores between cells of each type versus all other cells. We calculated Cohen’s d effect sizes to quantify the magnitude of TF activity differences, providing a standardized measure of effect size independent of sample size. We applied Benjamini-Hochberg false discovery rate correction (α = 0.05) across all TF-cell type combinations to control for multiple testing. We ranked TFs within each cell type by absolute Cohen’s d values to identify the most cell type-specific TF activities. This approach revealed which transcription factors show the strongest cell type-specific activity patterns through their binding at accessible chromatin regions, providing insights into cell type-specific gene regulatory programs.

#### Correlations between chromatin accessibility and gene expression

Of the 65,093 ACRs detected in our seedling snATAC-seq dataset, 35,424 were located in the promoters of one or more genes expressed in our seedling snRNA-seq dataset. For each of these ACR-gene pairs (n=55,455), we used the R function ‘cor’ with the ‘pariwise.complete.obs’ flag to calculate the correlation between mean accessibility and gene expression (per-cell type pseudobulk measured as TPM) across the 30 seedling cell types labeled in both our snATAC-seq and snRNA-seq seedling atlases. We then assigned each ACR-gene pair to a promoter feature bin based on the location of the center of the ACR, and counted the number of TF families with c4 TFBSs whose DAP peak summit was contained within the ACR. Finally, we binned all ACR-gene pairs by their feature bin and TF family count, and computed the median correlation coefficient (rho). We calculated both Pearson and Spearman correlation coefficients and saw similar results. We also calculated correlations across only the 19 cell types with at least 300 ACRs in which accessibility was highest in that cell type, and across only the 15,481 ACRs with accessibility >=0.5 in at least one cell type.

### Single Nuclei RNA-seq Analysis

#### Computing expression-based TF activity scores for promoter feature bins

For each TF, we identified all target genes with a c4 TFBS in a specific promoter feature, then randomly selected 20 subsets of 25 target genes, and computed expression-based TF activity scores as in ^3^ measured by Cohen’s d for each subset across each seedling, leaf, and flower bud cell types. A t-test was conducted to determine whether the Cohen’s d values for the 20 random subsets were significantly greater than or less than zero [R function t.test(cohens_d, mu = 0)]. This test was only performed when the TF had at least 50 total target genes with a TFBS in the promoter feature. We also performed an ANOVA test (R function aov) for each TF-cell type pair using values from all target gene subsets in all promoter regions, to test whether significant variance was associated with promoter region. The same procedure was used to score TF activity using TFBSs at different c scores and in the other brassica species.

#### Testing TFBS impact on expression as a function of TFBS location

First, TF expression in each cell type was calculated using the previously published snRNA-seq atlases, using per-cell type pseudobulk counts normalized to TPM, then scaled across all cell types. A TF was considered ‘strongly expressed’ in a cell type if its scaled expression value was >0.5. A promoter region was considered to have significant impact on cell type-specific expression if the TF activity scores calculated from selected subsets of target genes were significantly greater than zero (FDR-corrected t-test p-val <0.1), and the median Cohen’s d value across all subsets was greater than 0.5. For each TF, we identified all cell types in which it was strongly expressed, and then further filtered to cell types where at least one promoter feature showed significant impact using TFBSs at any c score. After all TFs were assigned to FBNs (see *Feature-based TF density calculations and assignments*), we summed the number of times a promoter region belonging to a TF’s FBN showed significant impact, as well as the number of times a promoter region outside a TF’s FBN showed significant impact, and then conducted a chi-square test (R function chisq.test) to determine whether there was a significant difference in rates.

### Bulk RNA-seq

#### Hormone treatment experiments

*A. thaliana* seeds were sterilized and cold stratified as described before^3^, then planted on nylon mesh in agar plates with the same growth medium composition as described before^3^. Seeds were sown on plates at a density of 3 rows per plate with ∼30 plants per row and plates were placed vertically in a growth chamber under conditions of 16 h light at 22°C, 8 h dark at 19°C for 5 days. On the 5th day, the nylon mesh and plants were transferred using forceps to agar plates containing either 2 μM ABA, 150 μM SA, or the standard growth medium without added hormones to serve as a parallel, time-matched negative control. Samples for all three conditions (ABA, SA, and mock) were collected at time points of 0, 1, 2, 4, 7, 24, and 48 hours, separating root and shoot tissues with a single razor blade cut at the time of collection. At each time point, five biological replicates per treatment were collected and flash frozen in liquid nitrogen then stored at -80°C. Three of the biological replicates were used for bulk RNA-seq. Sequencing was performed on an Illumina NovaSeq 6000 S4 flow cell, using 2x150 bp paired-end sequencing.

#### Data analysis

Raw reads were processed using BBTools v38.96 bbduk.sh^91^ for 1) removal of artifact sequence, RNA spike-in reads, PhiX reads, and reads containing any N, 2) quality trimming with method set at Q6, and 3) removal of reads with <50 bases, resulting in an average of 74.9M filtered reads per sample. Filtered reads from each library were aligned to the *A. thaliana* TAIR10 reference genome using HISAT2 v2.2.1127 ^92^. featureCounts^93^ was used to generate raw gene counts using Araport 11 gene annotations. Only primary hits assigned to the reverse strand were included (-s 2 -p --primary options). High Pearson correlation of raw gene counts was observed between all pairs of replicates (average correlation 0.992; min. correlation 0.89). Principal component analysis (PCA) and differential expression (DE) analysis were performed using DESeq2 v1.42.0^94^. PCA was performed separately for root and shoot samples on vst-normalized counts from all expressed genes. DE analysis was performed on all samples together, with design = ∼ tissue_treatment_timepoint. Crucially, to control for circadian-driven gene expression changes, DE contrasts were performed against time-matched mock controls. P-values and fold-changes for comparisons between ABA-or SA-treated samples and untreated samples from the same tissue at the exact same time point were extracted and subject to fold-change shrinkage (DESeq2::lfcShrink with type = “ashr”). At each timepoint and for each treatment and tissue, all genes were ranked according to log-fold change (post-shrinkage). Gene set enrichment analyses were then performed (using R package fgsea^95^) for TF target genes harboring TFBSs in each promoter region. Target genes harboring TFBSs in multiple promoter regions were assessed separately.

### Cis-Regulatory Module (CRM) identification and analysis

#### Peak clustering to identify CRMs

We identified CRMs by clustering DAP-seq peaks within distal regulatory regions for each species independently (cluster_*.ipynb scripts for *A. thaliana, A. lyrata, B. oleracea,* and *C. rubella*). We only used c4 peaks to focus on high-confidence binding events. We then merged peak summits that occurred within 300 bp of each other into clusters, reasoning that such proximally located TF binding sites likely represent functional regulatory modules. To ensure we identified bona fide multi-TF regulatory regions rather than isolated binding events, we filtered clusters to retain only those containing binding sites from at least 3 different TF families. For each identified cluster, we calculated summary statistics including genomic coordinates (chromosome, start, end, length), the number of peaks within the cluster, the number of distinct TF families represented, and conservation scores (mean, median, min, max) reflecting binding strength across species.

#### CRM-to-gene assignment

We assigned each CRM to its putative target gene by identifying the nearest downstream gene in a strand-aware manner. For each cluster, we calculated distances to the nearest genes on both the 5’ and 3’ flanking sides and determined which represented the most likely regulatory target based on genomic position and gene orientation. We recorded both the distance to the downstream gene and classified this distance into quartiles to enable categorical analysis of CRM-gene relationships. We annotated each CRM with information about its assigned target gene, including gene strand and distance metrics to both the transcription start site and coding sequence boundaries.

#### Cross-species CRM conservation analysis

We assessed CRM conservation across species by leveraging orthology relationships from OrthoFinder (conserved_crms.ipynb). We combined cluster summary tables from all four species and merged them with orthology information to link CRMs targeting orthologous genes. For each orthogroup-TF family combination, we counted how many species exhibited clusters with the same TF family composition targeting genes within that orthogroup. We created unique identifiers combining TF family composition with orthogroup membership to track conserved regulatory relationships across species. This approach allowed us to identify CRMs that maintain similar TF binding profiles at orthologous loci across divergent brassica lineages.

#### Validation with Chromatin Accessibility Data (A. thaliana only)

For *A. thaliana*, we intersected identified CRMs with single-nucleus ATAC-seq (snATAC-seq) peaks to validate their chromatin accessibility (annotate_ath.ipynb). We used BEDTools to identify clusters with at least 60% reciprocal overlap with accessible chromatin regions and marked these as occurring within differentially accessible regions (DARs). We extracted gene lists for CRMs that overlapped with accessible chromatin to enable downstream functional enrichment analysis.

#### H3K27me3 and CRM Overlap Analysis

To assess the co-occurrence of H3K27me3 chromatin marks at distal cis-regulatory modules (CRMs), we analyzed publicly available A. thaliana H3K27me3 ChIP-seq peak calls from two datasets: flower developmental stages (t0, t2, intermediate, and late; two biological replicates each; GEO series GSE71564; doi:10.3390/epigenomes1020008) and vegetative root and shoot tissues (two biological replicates each; GSE155502; doi:10.1101/gr.273771.120). We restricted the analysis to intergenic regions, excluding gene bodies and proximal promoters (2 kb upstream and 500 bp downstream of transcription start sites, strand-specific). For each CRM, overlap with H3K27me3 peaks was assessed separately for each sample, and an “any-tissue” union metric was defined as overlap in at least one of the 12 samples. To establish statistical significance, we performed permutation testing by randomly repositioning CRMs within intergenic space while preserving their length distribution (n = 500 permutations). For each permutation, we calculated the fraction of randomized CRMs overlapping H3K27me3 peaks and computed empirical p-values as (k + 1)/(n + 1), where k is the number of permutations with overlap fractions greater than or equal to the observed value. Across the any-tissue union, distal CRMs showed 1.45-fold enrichment for H3K27me3 overlap compared with length-matched random intergenic regions (p < 0.002).

### GO enrichment analysis

We used clusterprofiler v4.6.2 to perform gene ontology (GO) term enrichment for different gene sets, for instance, list of target genes bound by each TF in distal region. We first converted the TAIR gene names into the Entrez ids. Then we calculated enrichment values for Biological Processes and filtered for enriched terms with level six to avoid broad GO terms. We then used the “simplify” function to combine terms with a similarity cutoff of 70%.

## Data Availability

The raw snATAC-seq FASTQ sequence data files were submitted to the National Center for Biotechnology Information under BioProject no. PRJNA1464623. The processed snATAC-seq data including single-cell gene expression matrices (.mtx format) for each library and compiled anndata files for each species and tissue, were submitted to the National Center for Biotechnology Information under GEO accession no. GSE332675. Additionally, gene expression data from the *Arabidopsis thaliana* root and shoot hormone treatment time course have been deposited in GEO under accession no. GSE331340. Gene features, tables with DAPseq peaks assigned to distal, promoter and downstream regions, snATAC-seq and snRNA-seq summary tables are available in Zenodo (https://doi.org/10.5281/zenodo.18407667)^96^. Published epigenomic datasets used for chromatin mark enrichment analysis are listed with accession numbers in Methods ("Epigenetic mark enrichment test").

## Code Availability

Scripts used for DAP-seq and single-nuclei analyses are available in a git repository at https://code.jgi.doe.gov/AMoralesCruz/regulatory-neighborhoods

## Acknowledgments

The work conducted by the U.S. Department of Energy Joint Genome Institute (https://ror.org/04xm1d337), a DOE Office of Science User Facility, is supported by the Office of Science of the U.S. Department of Energy operated under Contract No. DE-AC02-05CH11231.

## Author contributions

Conceptualization, AMC, LAB, RCO, SIG; Methodology, AMC, JJ, LAB, PW, RCO, SIG, YZ;

Validation, AMC, LAB, RCO, SIG, YZ; Formal Analysis, AMC, LAB, LY, RCO, SIG; Investigation, PW, YZ; Writing, AMC, LAB, PW, RCO, SIG, YZ; Visualization, AMC, LY, SIG; Supervision, CGD, LAB, RCO.

## Extended Data figures and tables

### Extended Data Figures

**Extended Data Fig. 1.**
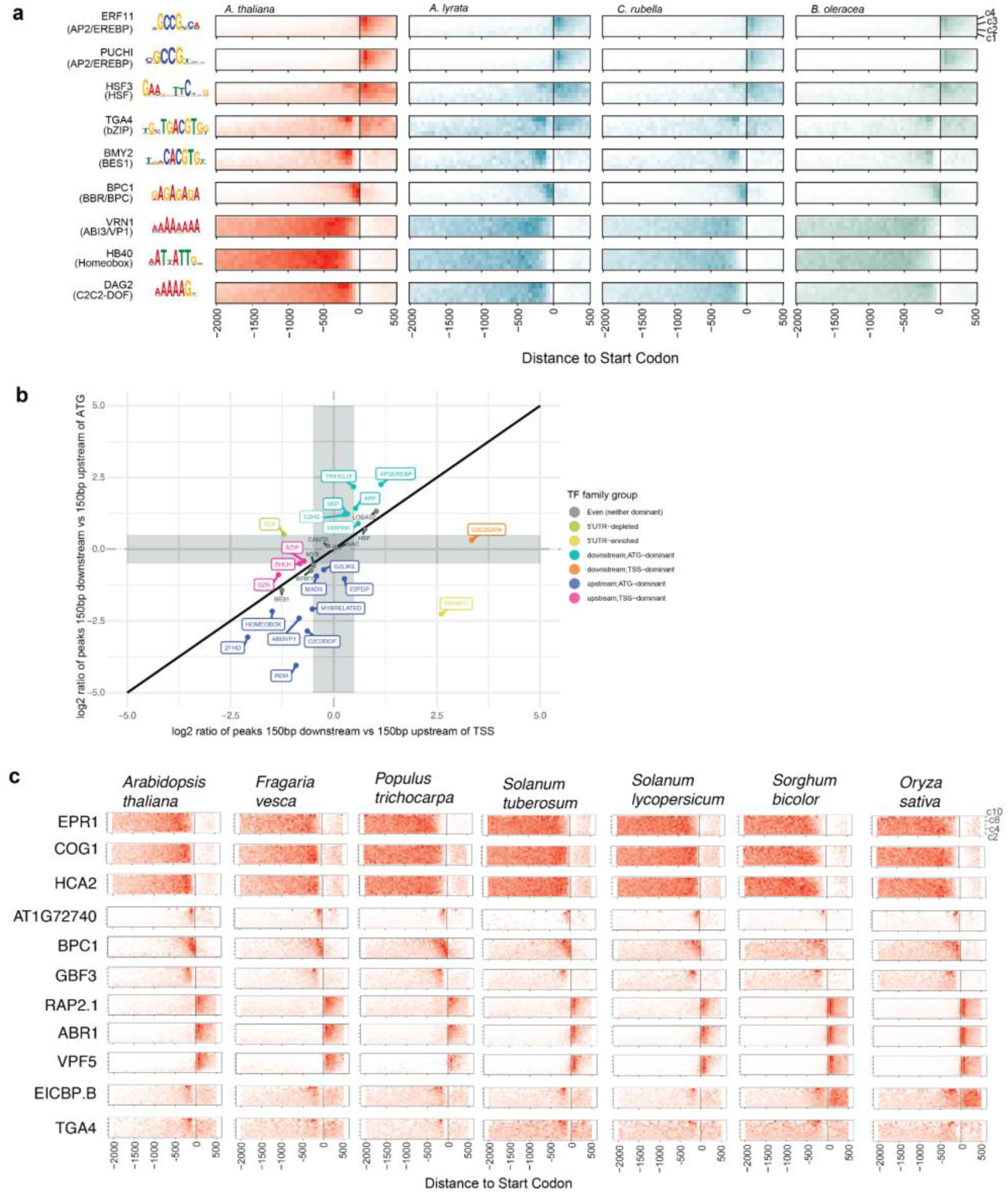
**a)** Examples of TFs with conserved TFBS location patterns across c scores, anchored on either the TSS or the start codon for *A. thaliana*, *A. lyrata*, *Capsella rubella* and *Brassica oleracea,* illustrating conservation of location patterns between species. **b)** Transcription factor binding site enrichment distinguishes TSS-driven and CDS-driven TF positioning. The x-axis shows log2 of the ratio of total DAP peaks associated with each TF family in the region 150 bp before vs. the region 150 bp after the TSS, and the y-axis shows the same for the regions 150 bp before and after the start codon. TF families where the absolute fold change between the ratios was > 0.5 are labeled as either TSS-dominant or ATG-dominant, reflecting a sharper change in TFBS density at the TSS and or ATG respectively. **c)** Locational TFBS conservation patterns anchored on the start codon for seven of the species in the deep conservation dataset.

**Extended Data Fig. 2.**
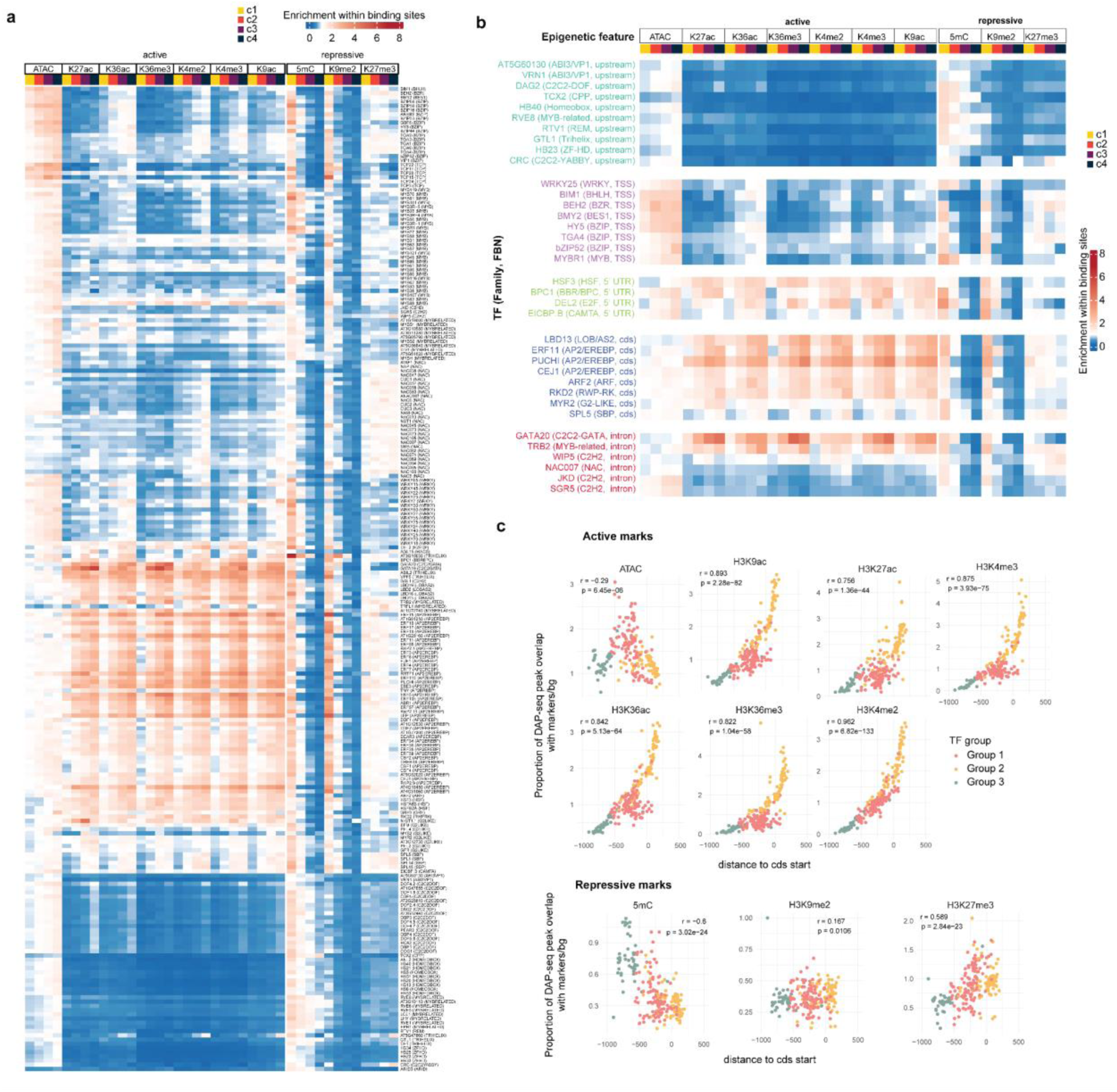
**a)** Enrichment analysis of the degree of overlap between ten epigenetic markers and TFBSs for all TFs tested, categorized by c score (colored bins). The results indicate varying levels of enrichment, from depletion (blue) to enriched (red), providing insights into the relationships between epigenetic modifications and transcription factor activity across the analyzed datasets. **b)** Selected TF from Extended Data Fig. 2a. grouped by TF family and FBN to show differences of epigenomic patterns across groups. **c)** Correlation between median distance to TSS and enrichment of epigenetic marks of all TFs. Colors represent groupings of TFs based on the epigenetic mark overlap pattern seen in Extended Data Fig. 2a.

**Extended Data Fig. 3.**
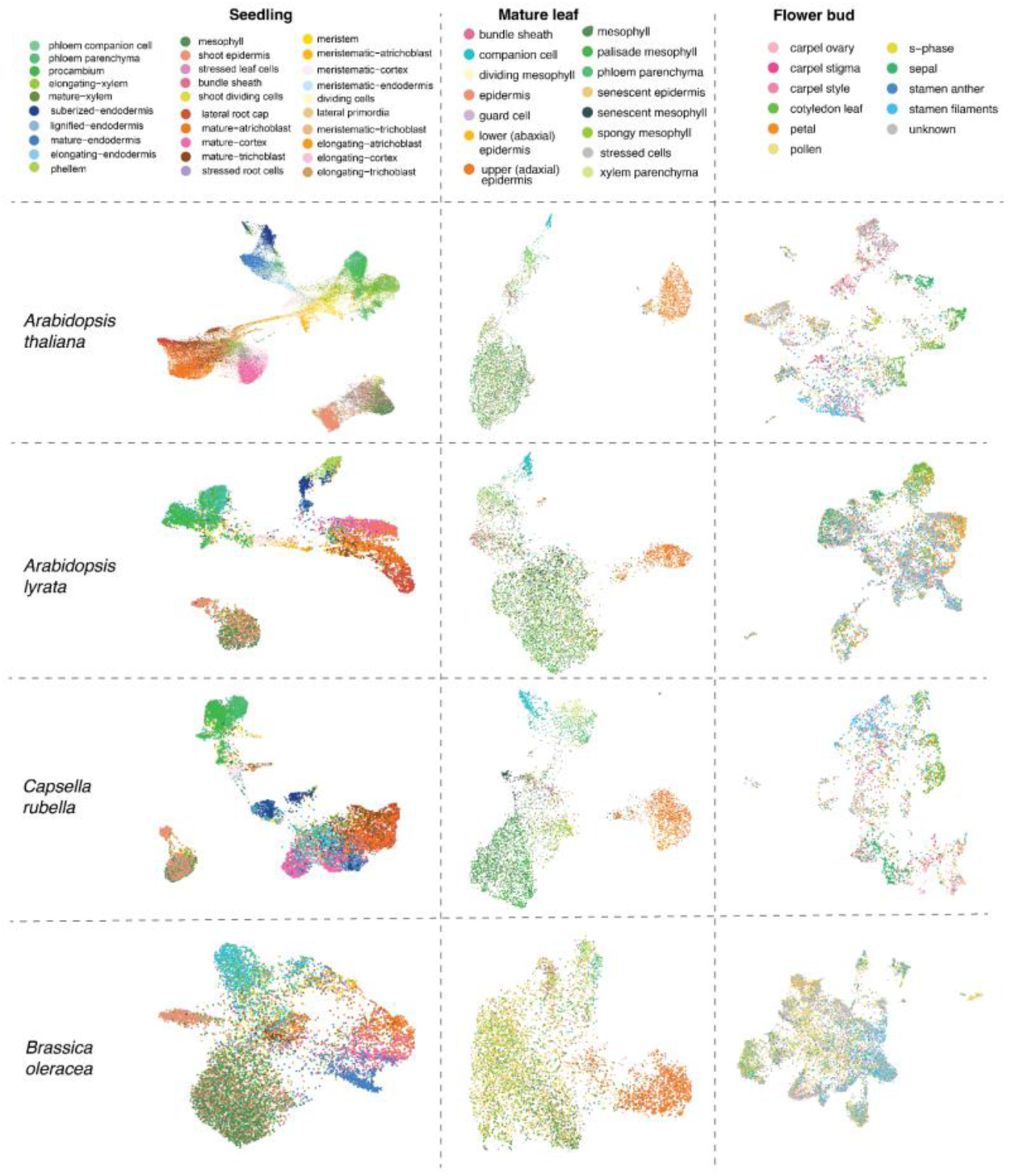
UMAP visualization of single-nuclei ATAC-seq data from four Brassicaceae species and three tissues. Each dot represents an individual cell, with colors indicating the specific cell type assigned to that cell based on clustering analysis. This representation allows for the comparison of chromatin accessibility patterns across different biological contexts, highlighting the diversity and distribution of cell types within the sampled species and tissues.

**Extended Data Fig. 4.**
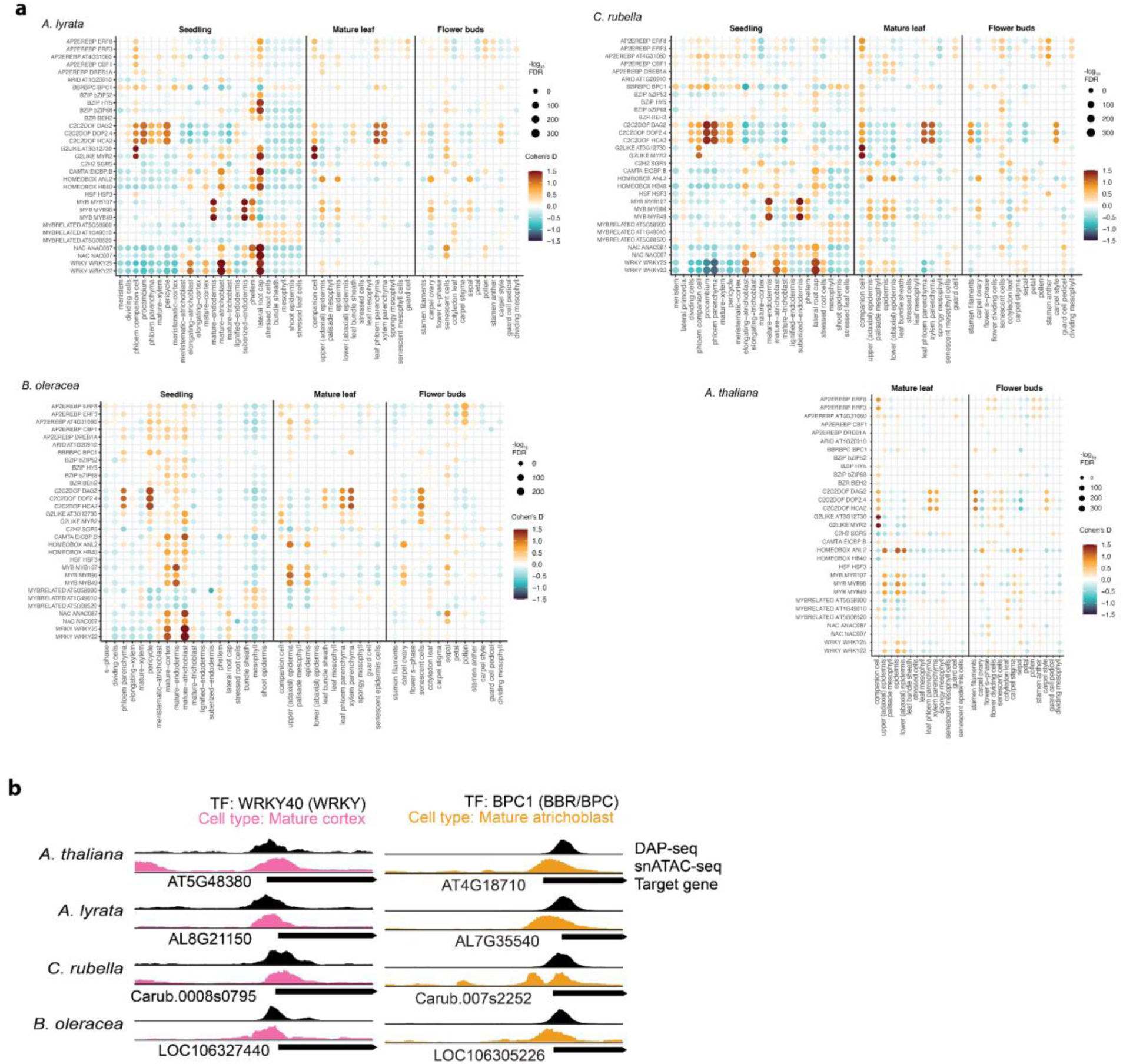
**a)** Bubble heatmaps illustrating the enrichment of transcription factor binding sites (TFBSs) in DARs from multiple tissues and species. The color of each bubble reflects the magnitude of enrichment, quantified using Cohen’s D, while the size of the bubble indicates the significance of the enrichment, represented by -log_10_(FDR p-value). Only selected transcription factors are displayed from a comprehensive analysis involving four different species and three distinct tissues, highlighting patterns of regulatory activity across biological contexts. **b)** Examples of open chromatin regions in species analyzed, showcasing regions with enhanced accessibility and potential regulatory activity.

**Extended Data Fig. 5.**
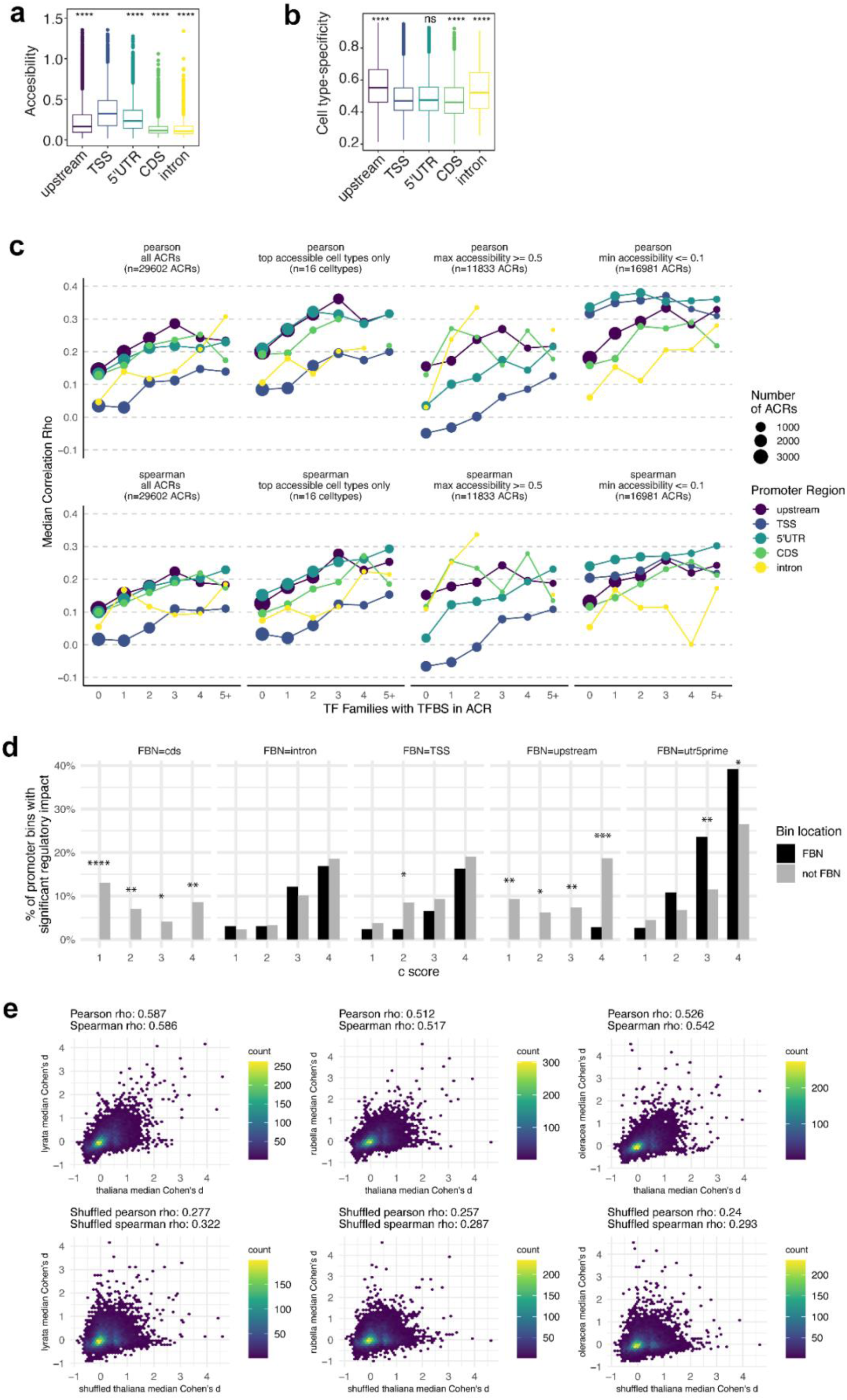
**a)** Mean accessibility and **b)** cell type-specificity of all ACRs classified by promoter region. Asterisks indicate statistical significance from independent, unpaired two-sided *t*-tests comparing each individual feature category against the TSS category (serving as the reference group). ns: *p* > 0.05, * *p* ≤ 0.05, ** *p* ≤ 0.01, *** *p* ≤ 0.001, **** *p* ≤ 0.0001. **c)** Parametric (top row) and non-parametric (bottom row) correlations between accessibility of an ACR and expression of the associated gene across *A. thaliana* seedling cell types. Each point represents the median correlation coefficient for a subset of ACRs binned by location within the promoter (color) and number of overlapping TFBSs from unique TF families (x-axis). The size of the circle represents the number of ACRs considered. In the second column of plots, correlations were calculated only across the 16 cell types with high overall accessibility, in the third column correlations were summarised only for ACRs with high accessibility in at least one cell type, and in the fourth correlations were summarised only for ACRs effectively closed in at least one cell type. **d)** Percent of promoter regions with a significant TF activity score, from all promoters where at least one region (at any c score) had a significant TF activity score. Results are subsetted into different panels for groups of TFs assigned to different feature-based neighborhoods (FBNs), and results for each c score are subsetted into bars representing promoter regions corresponding to the TF’s FBN (black) or regions outside the TF’s FBN (gray). Asterisks represent the significance of the difference between black and gray bars according to a chi-squared test (*p ≤ 0.05, **p ≤ 0.01, ***p ≤ 0.001, ****p ≤ 0.0001). **e**) Each plot represents a comparison of the median TF activity score (for 20 random subsets of target genes) for a given TF/cell type/promoter region in *A. thaliana* (x-axis) vs. the median for the same TF/cell type/promoter region in another brassica species (y-axis). Point density is shown with color, and parametric and non-parametric correlation coefficients are shown at the top. The bottom row of plots shows the same calculations after shuffling the promoter region label for A. thaliana values within the same TF and cell type.

**Extended Data Fig. 6.**
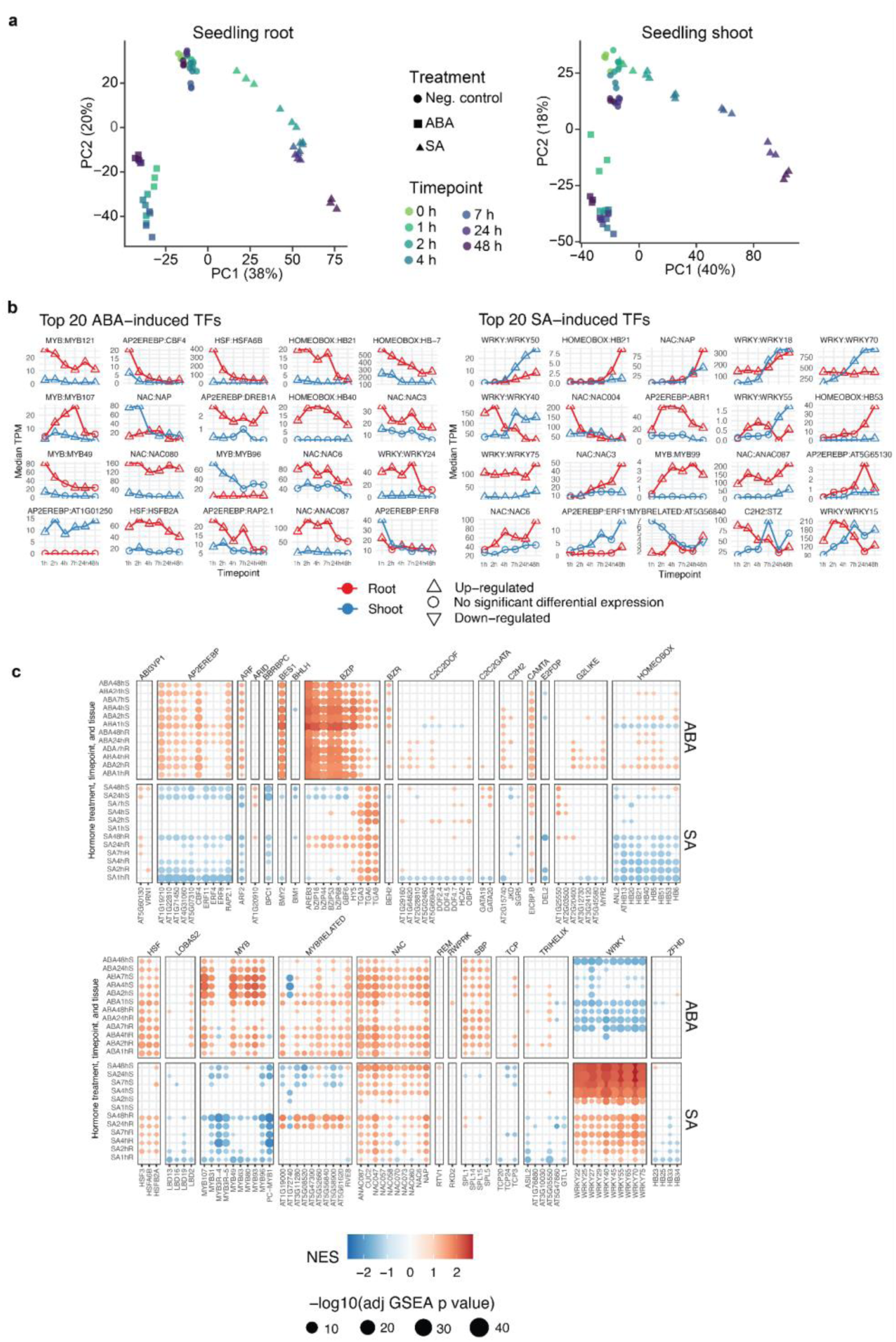
**a)** Principal component analysis of genome-wide transcriptional profile for root and shoot samples from *A. thaliana* plants exposed to a 48-hr treatment time course with either abscisic acid or salicylic acid. **b)** Time course of median TPM values across three replicates for top 20 TFs showing hormone-induced expression in each treatment. **c)** Results of gene set enrichment test for all c4 target genes of each TF among genes ordered by their hormone response in each tissue at each timepoint. NES = Normalized Effect Size.

**Extended Data Fig. 7.**
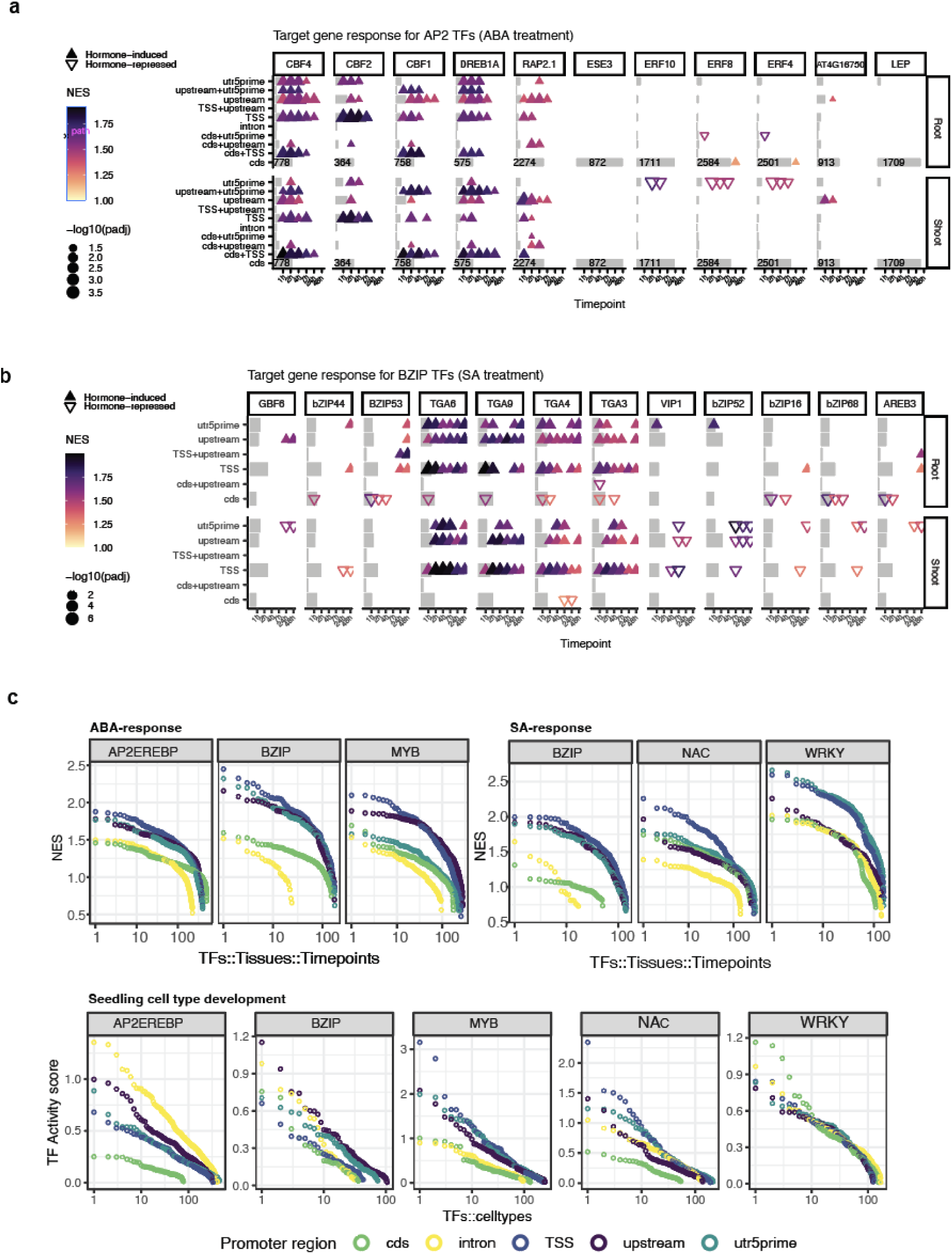
Gene set enrichment analysis for target genes of selected bZIP TFs during ABA treatment (**a**) and SA treatment (**b**). Gray bars represent the proportion of all target genes for the TF that fall into each TFBS location category. **c**) Top NES values for each promoter region across all TFs, tissues, and timepoints for three top-responding TF families for each hormone (top panel), and top TF activity scores representing target gene cell type specificity for each promoter region, for the same set of TF families.

**Extended Data Fig. 8.**
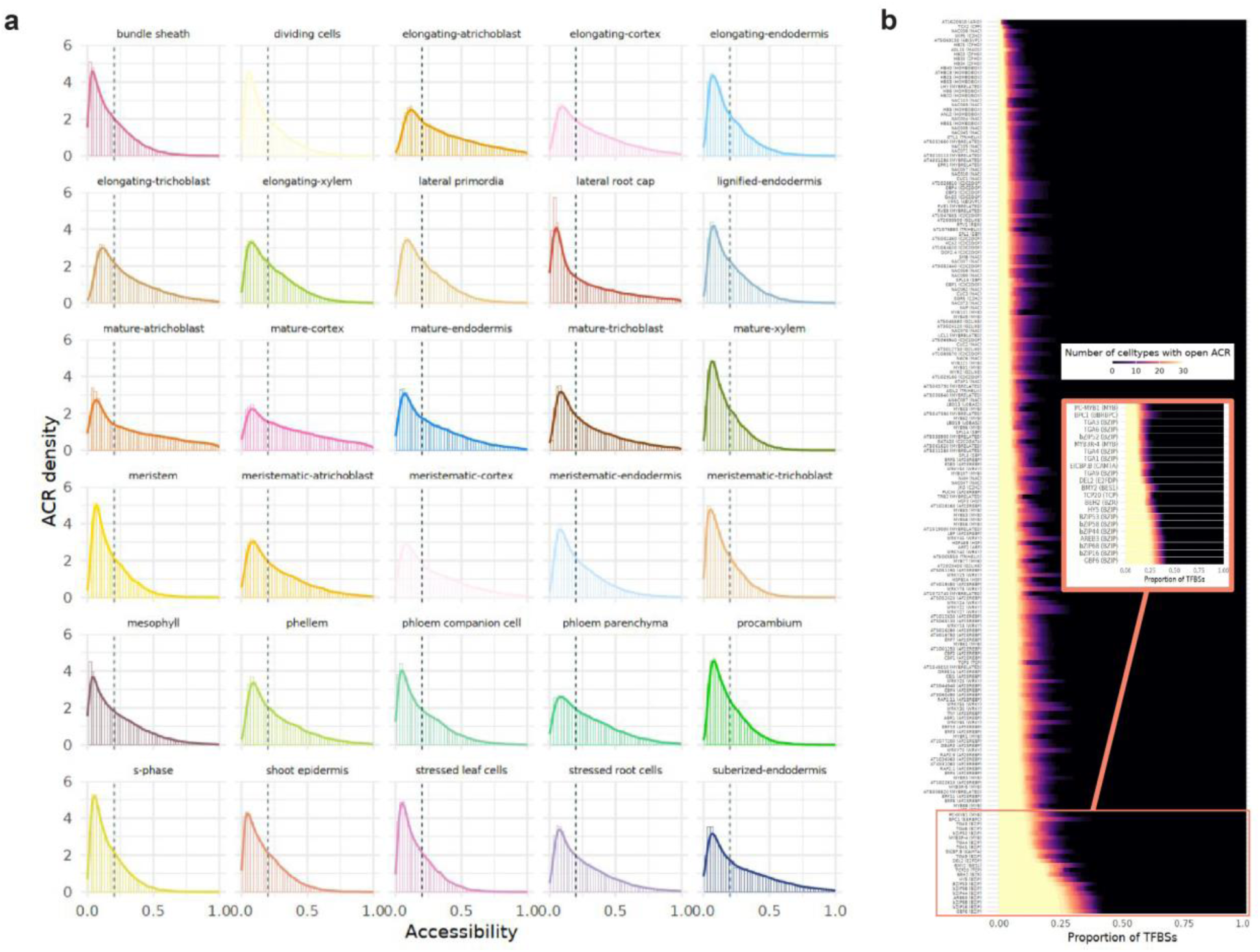
**a**) Distribution of mean accessibility values across all promoter ACRs for each cell type. Dotted line represents the threshold used for defining binary ‘open’ state (mean accessibility = 0.2). **b**) Proportion of TFBSs for each TF that overlap ACRs open in a given number of cell types. TFBSs not overlapping an ACR were coded as 0. Inset shows a zoomed-in view of the set of TFs with the highest proportion of TFBSs in constitutively-accessible chromatin.

**Extended Data Fig. 9.**
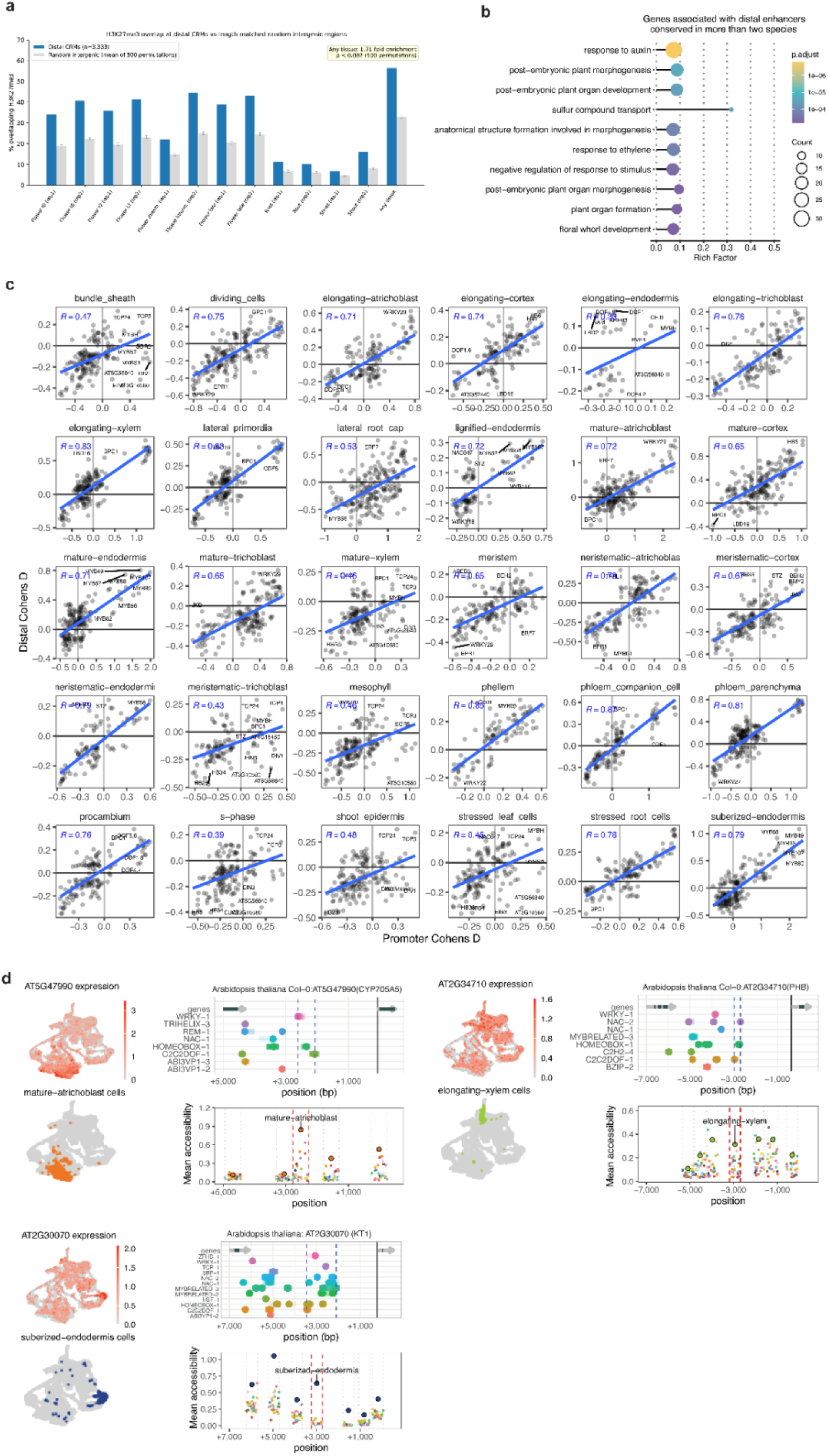
**a)** H3K27me3 enrichment at distal CRMs. Grouped bar chart showing the fraction of distal CRMs (n = 3,333) overlapping H3K27me3 peaks (blue) compared to the mean fraction from 500 length-matched random intergenic region permutations (grey), across 12 H3K27me3 ChIP-seq samples from flower developmental stages (t0, t2, intermediate, late), root, and shoot tissues, plus a union across all tissues ("Any tissue"). CRMs show 1.45-fold enrichment for H3K27me3 overlap relative to random intergenic regions (any-tissue union, empirical p < 0.002). **b)** Significantly enriched (adjusted p < 0.01) GO terms for genes associated with distal CRMs conserved in two species or more. **c**) Scatterplot of TF enrichment magnitudes (Cohen’s d) for cell types from DAP-seq peaks in c4 promoter vs distal CRMs in ACRs. **d**) Other examples of candidate enhancers. The left side shows expression of the gene associated with the candidate enhancer across single *A. thaliana* seedling cells (top) and the location of cells annotated as the cell type with highest overall expression (bottom). The right side shows DAP-seq peaks (top) in and near the candidate enhancer region (dotted lines), and mean accessibility values (bottom) for each cell type (color) for ACRs in the same region. ACRs overlapping the candidate enhancer region are outlined in red, other ACR boundaries are shown with dotted gray lines.

### Supplementary Tables

**Supplementary Table 1**. Classification of TFs by clusters according to the c4 TFBS densities as shown in Fig. 2F.

**Supplementary Table 2**. Quality control metrics of the single-nuclei ATAC-seq samples used in this study.

**Supplementary Table 3**. Cell type-specific (3 cell types or less) differentially accessible regions (DARs) with the maximum conservation (c score) DAP-seq peak intersecting the DAR.

**Supplementary Table 4**. Complete catalog of 3,333 distal cis-regulatory modules (CRMs) identified in A. thaliana, including genomic coordinates, TF family composition, assigned target gene, conservation score, H3K27me3 overlap status, upstream intergenic distance, and functional classification.

**Supplementary Table 5**. CRM functional enrichment analysis. Fisher’s exact test results for hormone pathway and functional group enrichment among CRM-associated genes against genome-wide background.

**Supplementary Table 6**. Per-gene detail for all 23 ARF transcription factors. Phylogenetic class and functional classification per Mutte et al. 2018^96^. Gene annotations from Araport11.

**Supplementary Table 7**: Upstream intergenic length analysis by gene family, including CRM-associated vs non-CRM member comparison and 95th percentile tail enrichment.

**Supplementary Table 8**. Cis-regulatory modules (CRMs) in cell type-specific differentially accessible regions (DARs)

**Supplementary Table 9**. 102 top candidate enhancers, representing distal CRMs where both accessibility and expression of the associated gene was highest in the same cell type.

## References

1. Voichek, Y., Hristova, G., Mollá-Morales, A., Weigel, D. & Nordborg, M. Widespread position-dependent transcriptional regulatory sequences in plants. Nat Genet 56, 2238–2246 (2024).

2. Galli, M. et al. Transcription factor binding divergence drives transcriptional and phenotypic variation in maize. Nat. Plants 11, 1205–1219 (2025).

3. Baumgart, L. A. et al. Recruitment, rewiring and deep conservation in flowering plant gene regulation. Nat. Plants 11, 1514–1527 (2025).

4. Dorrity, M. W. et al. The regulatory landscape of Arabidopsis thaliana roots at single-cell resolution. Nat. Commun. 12, 3334 (2021).

5. Zhang, X. et al. A spatially resolved multi-omic single-cell atlas of soybean development. Cell 188, 550–567.e19 (2025).

6. ENCODE Project Consortium. An integrated encyclopedia of DNA elements in the human genome. Nature 489, 57–74 (2012).

7. Roadmap Epigenomics Consortium et al. Integrative analysis of 111 reference human epigenomes. Nature 518, 317–330 (2015).

8. Visel, A. et al. A high-resolution enhancer atlas of the developing telencephalon. Cell 152, 895–908 (2013).

9. Yu, C.-P., Lin, J.-J. & Li, W.-H. Positional distribution of transcription factor binding sites in Arabidopsis thaliana. Sci. Rep. 6, 25164 (2016).

10. Zou, C. et al. Cis-regulatory code of stress-responsive transcription in Arabidopsis thaliana. Proc. Natl. Acad. Sci. U. S. A. 108, 14992–14997 (2011).

11. Heyndrickx, K. S., Van de Velde, J., Wang, C., Weigel, D. & Vandepoele, K. A functional and evolutionary perspective on transcription factor binding in Arabidopsis thaliana. Plant Cell 26, 3894–3910 (2014).

12. Lu, Z., Hofmeister, B. T., Vollmers, C., DuBois, R. M. & Schmitz, R. J. Combining ATAC-seq with nuclei sorting for discovery of cis-regulatory regions in plant genomes. Nucleic Acids Res. 45, e41 (2017).

13. Cheng, K. et al. Histone tales: lysine methylation, a protagonist in Arabidopsis development. J. Exp. Bot. 71, 793–807 (2020).

14. Mahrez, W. et al. H3K36ac Is an Evolutionary Conserved Plant Histone Modification That Marks Active Genes. Plant Physiol. 170, 1566–1577 (2016).

15. Li, Y. et al. The histone methyltransferase SDG8 mediates the epigenetic modification of light and carbon responsive genes in plants. Genome Biol. 16, 79 (2015).

16. Inagaki, S. et al. Gene-body chromatin modification dynamics mediate epigenome differentiation in Arabidopsis. EMBO J. 36, 970–980 (2017).

17. Luo, C. et al. Integrative analysis of chromatin states in Arabidopsis identified potential regulatory mechanisms for natural antisense transcript production. Plant J. 73, 77–90 (2013).

18. Kawakatsu, T. et al. Epigenomic Diversity in a Global Collection of Arabidopsis thaliana Accessions. Cell 166, 492–505 (2016).

19. Jackson, J. P. et al. Dimethylation of histone H3 lysine 9 is a critical mark for DNA methylation and gene silencing in Arabidopsis thaliana. Chromosoma 112, 308–315 (2004).

20. Grass, J. A. et al. GATA-1-dependent transcriptional repression of GATA-2 via disruption of positive autoregulation and domain-wide chromatin remodeling. Proc. Natl. Acad. Sci. U. S. A. 100, 8811–8816 (2003).

21. Marand, A. P. et al. The genetic architecture of cell type-specific cis regulation in maize. Science 388, eads6601 (2025).

22. Marand, A. P., Chen, Z., Gallavotti, A. & Schmitz, R. J. A cis-regulatory atlas in maize at single-cell resolution. Cell 184, 3041–3055.e21 (2021).

23. Farmer, A., Thibivilliers, S., Ryu, K. H., Schiefelbein, J. & Libault, M. Single-nucleus RNA and ATAC sequencing reveals the impact of chromatin accessibility on gene expression in Arabidopsis roots at the single-cell level. Mol. Plant 14, 372–383 (2021).

24. Maher, K. A. et al. Profiling of accessible chromatin regions across multiple plant species and cell types reveals common gene regulatory principles and new control modules. Plant Cell 30, 15–36 (2018).

25. Kerstens, M. et al. Two deeply conserved non-coding sequences control PLETHORA1/2 expression and coordinate embryo and root development. Plant Commun. 6, 101466 (2025).

26. Ota, R., Ohkubo, Y., Yamashita, Y., Ogawa-Ohnishi, M. & Matsubayashi, Y. Shoot-to-root mobile CEPD-like 2 integrates shoot nitrogen status to systemically regulate nitrate uptake in Arabidopsis. Nat. Commun. 11, 641 (2020).

27. Konishi, M. & Yanagisawa, S. Transcriptional repression caused by Dof5.8 is involved in proper vein network formation in Arabidopsis thaliana leaves. J. Plant Res. 128, 643–652 (2015).

28. Guo, Y., Qin, G., Gu, H. & Qu, L.-J. Dof5.6/HCA2, a Dof transcription factor gene, regulates interfascicular cambium formation and vascular tissue development in Arabidopsis. Plant Cell 21, 3518–3534 (2009).

29. Kubo, H., Kishi, M. & Goto, K. Expression analysis of ANTHOCYANINLESS2 gene in Arabidopsis. Plant Sci. 175, 853–857 (2008).

30. Palatnik, J. F. et al. Control of leaf morphogenesis by microRNAs. Nature 425, 257–263 (2003).

31. Zhao, C., Hanada, A., Yamaguchi, S., Kamiya, Y. & Beers, E. P. The Arabidopsis Myb genes MYR1 and MYR2 are redundant negative regulators of flowering time under decreased light intensity. Plant J. 66, 502–515 (2011).

32. Verweij, W. et al. Functionally similar WRKY proteins regulate vacuolar acidification in petunia and hair development in Arabidopsis. Plant Cell 28, 786–803 (2016).

33. Gou, M. et al. The MYB107 transcription factor positively regulates suberin biosynthesis. Plant Physiol. 173, 1045–1058 (2017).

34. Chen, H. et al. Roles of arabidopsis WRKY18, WRKY40 and WRKY60 transcription factors in plant responses to abscisic acid and abiotic stress. BMC Plant Biol. 10, 281 (2010).

35. Mirabella, R. et al. WRKY40 and WRKY6 act downstream of the green leaf volatile E-2-hexenal in Arabidopsis. Plant J. 83, 1082–1096 (2015).

36. Galway, M. E. et al. The TTG gene is required to specify epidermal cell fate and cell patterning in the Arabidopsis root. Dev. Biol. 166, 740–754 (1994).

37. Balcerowicz, D., Schoenaers, S. & Vissenberg, K. Cell Fate Determination and the Switch from Diffuse Growth to Planar Polarity in Arabidopsis Root Epidermal Cells. Front. Plant Sci. 6, 1163 (2015).

38. Liu, C., Cheng, Y.-J., Wang, J.-W. & Weigel, D. Prominent topologically associated domains differentiate global chromatin packing in rice from Arabidopsis. Nat. Plants 3, 742–748 (2017).

39. Nobori, T. et al. A rare PRIMER cell state in plant immunity. Nature 638, 197–205 (2025).

40. Liu, Q. et al. Multiome in the same cell reveals the impact of osmotic stress on Arabidopsis root tip development at single-cell level. Adv. Sci. (Weinh.) 11, e2308384 (2024).

41. Meng, F. et al. Genomic editing of intronic enhancers unveils their role in fine-tuning tissue-specific gene expression in Arabidopsis thaliana. Plant Cell 33, 1997–2014 (2021).

42. Zhu, X. et al. Molecular dissection of an intronic enhancer governing cold-induced expression of the vacuolar invertase gene in potato. Plant Cell 36, 1985–1999 (2024).

43. Song, L. et al. A transcription factor hierarchy defines an environmental stress response network. Science 354, aag1550 (2016).

44. Birkenbihl, R. P., Kracher, B., Roccaro, M. & Somssich, I. E. Induced genome-wide binding of three Arabidopsis WRKY transcription factors during early MAMP-triggered immunity. Plant Cell 29, 20–38 (2017).

45. Chen, K. et al. Abscisic acid dynamics, signaling, and functions in plants. J. Integr. Plant Biol. 62, 25–54 (2020).

46. Han, Q. et al. Salicylic acid-activated BIN2 phosphorylation of TGA3 promotes Arabidopsis PR gene expression and disease resistance. EMBO J. 41, e110682 (2022).

47. Kobayashi, Y. et al. Abscisic acid-activated SNRK2 protein kinases function in the gene-regulation pathway of ABA signal transduction by phosphorylating ABA response element-binding factors. Plant J. 44, 939–949 (2005).

48. Reyna-Llorens, I. et al. Ancient duons may underpin spatial patterning of gene expression in C4 leaves. Proc. Natl. Acad. Sci. U. S. A. 115, 1931–1936 (2018).

49. Kok, J. Y., Harvey, Z. H., Axelsson, E. & Berger, F. Nucleosome positioning shapes cryptic antisense transcription. PLoS Genet. 22, e1012078 (2026).

50. Beernink, B. M., Vogel, J. P. & Lei, L. Enhancers in plant development, adaptation and evolution. Plant Cell Physiol. 66, 461–476 (2025).

51. Zhu, B., Zhang, W., Zhang, T., Liu, B. & Jiang, J. Genome-Wide Prediction and Validation of Intergenic Enhancers in Arabidopsis Using Open Chromatin Signatures. Plant Cell 27, 2415–2426 (2015).

52. Marand, A. P., Eveland, A. L., Kaufmann, K. & Springer, N. M. Cis-regulatory elements in plant development, adaptation, and evolution. Annu. Rev. Plant Biol. 74, 111–137 (2023).

53. Bao, D., Chang, S., Li, X. & Qi, Y. Advances in the study of auxin early response genes: Aux/IAA, GH3, and SAUR. Crop J. 12, 964–978 (2024).

54. Kroll, C. K. & Brenner, W. G. Cytokinin signaling downstream of the his-asp phosphorelay network: Cytokinin-regulated genes and their functions. Front. Plant Sci. 11, 604489 (2020).

55. Guo, P. et al. Comparative analysis of the RTFL peptide family on the control of plant organogenesis. J. Plant Res. 128, 497–510 (2015).

56. Alarcia, A., Primo-Capella, A., Perpiñán, E., Rossetto, P. & Ferrándiz, C. DEVIL peptides control cell growth and differentiation in different developmental processes. bioRxiv (2023) doi:10.1101/2023.09.13.557675.

57. Fichtner, F. et al. Functional features of TREHALOSE-6-PHOSPHATE SYNTHASE1, an essential enzyme in Arabidopsis. Plant Cell 32, 1949–1972 (2020).

58. Rojas-Murcia, N. et al. High-order mutants reveal an essential requirement for peroxidases but not laccases in Casparian strip lignification. Proc. Natl. Acad. Sci. U. S. A. 117, 29166–29177 (2020).

59. Blaschek, L., Murozuka, E., Serk, H., Ménard, D. & Pesquet, E. Different combinations of laccase paralogs nonredundantly control the amount and composition of lignin in specific cell types and cell wall layers in Arabidopsis. Plant Cell 35, 889–909 (2023).

60. Kim, J.-Y. et al. Distinct identities of leaf phloem cells revealed by single cell transcriptomics. Plant Cell 33, 511–530 (2021).

61. Ladwig, F. et al. Siliques are Red1 from Arabidopsis acts as a bidirectional amino acid transporter that is crucial for the amino acid homeostasis of siliques. Plant Physiol. 158, 1643–1655 (2012).

62. Liu, S., Kracher, B., Ziegler, J., Birkenbihl, R. P. & Somssich, I. E. Negative regulation of ABA signaling by WRKY33 is critical for Arabidopsis immunity towards Botrytis cinerea 2100. Elife 4, e07295 (2015).

63. Clark, R. M., Wagler, T. N., Quijada, P. & Doebley, J. A distant upstream enhancer at the maize domestication gene tb1 has pleiotropic effects on plant and inflorescent architecture. Nat. Genet. 38, 594–597 (2006).

64. Salvi, S. et al. Conserved noncoding genomic sequences associated with a flowering-time quantitative trait locus in maize. Proc. Natl. Acad. Sci. U. S. A. 104, 11376–11381 (2007).

65. Adrian, J. et al. cis-Regulatory elements and chromatin state coordinately control temporal and spatial expression of FLOWERING LOCUS T in Arabidopsis. Plant Cell 22, 1425–1440 (2010).

66. Studer, A., Zhao, Q., Ross-Ibarra, J. & Doebley, J. Identification of a functional transposon insertion in the maize domestication gene tb1. Nat. Genet. 43, 1160–1163 (2011).

67. Liu, L. et al. KRN4 controls quantitative variation in maize kernel row number. PLoS Genet. 11, e1005670 (2015).

68. Zicola, J., Liu, L., Tänzler, P. & Turck, F. Targeted DNA methylation represses two enhancers of FLOWERING LOCUS T in Arabidopsis thaliana. Nat. Plants 5, 300–307 (2019).

69. Yan, W. et al. Dynamic control of enhancer activity drives stage-specific gene expression during flower morphogenesis. Nat. Commun. 10, 1705 (2019).

70. Mendieta, J. P. et al. Investigating the cis-regulatory basis of C3 and C4 photosynthesis in grasses at single-cell resolution. Proc. Natl. Acad. Sci. U. S. A. 121, e2402781121 (2024).

71. Yan, H. et al. A single-cell rice atlas integrates multi-species data to reveal cis-regulatory evolution. Nat. Plants 11, 2050–2071 (2025).

72. McDonald, B. R. et al. Enhancers associated with unstable RNAs are rare in plants. Nat. Plants 10, 1246–1257 (2024).

73. Stergachis, A. B. et al. Exonic transcription factor binding directs codon choice and affects protein evolution. Science 342, 1367–1372 (2013).

74. Brooks, M. D. et al. Network Walking charts transcriptional dynamics of nitrogen signaling by integrating validated and predicted genome-wide interactions. Nat. Commun. 10, 1569 (2019).

75. BBMap. SourceForge https://sourceforge.net/projects/bbmap/ (2022).

76. Langmead, B. & Salzberg, S. L. Fast gapped-read alignment with Bowtie 2. Nat. Methods 9, 357–359 (2012).

77. Li, H. et al. The Sequence Alignment/Map format and SAMtools. Bioinformatics 25, 2078–2079 (2009).

78. Zhang, Y. et al. Model-based analysis of ChIP-Seq (MACS). Genome Biol. 9, R137 (2008).

79. Bailey, T. L., Johnson, J., Grant, C. E. & Noble, W. S. The MEME suite. Nucleic Acids Res. 43, W39–49 (2015).

80. Quinlan, A. R. & Hall, I. M. BEDTools: a flexible suite of utilities for comparing genomic features. Bioinformatics 26, 841–842 (2010).

81. Emms, D. M. & Kelly, S. OrthoFinder: phylogenetic orthology inference for comparative genomics. Genome Biol. 20, 238 (2019).

82. Lister, R. et al. Highly integrated single-base resolution maps of the epigenome in Arabidopsis. Cell 133, 523–536 (2008).

83. Schmitz, R. J. et al. Patterns of population epigenomic diversity. Nature 495, 193–198 (2013).

84. Ageeva-Kieferle, A. et al. Nitric oxide coordinates growth, development, and stress response via histone modification and gene expression. Plant Physiol. 187, 336–360 (2021).

85. Cui, X. et al. REF6 recognizes a specific DNA sequence to demethylate H3K27me3 and regulate organ boundary formation in Arabidopsis. Nat. Genet. 48, 694–699 (2016).

86. Galbraith, D. W. Simultaneous flow cytometric quantification of plant nuclear DNA contents over the full range of described angiosperm 2C values. Cytometry A 75, 692–698 (2009).

87. Zheng, G. X. Y. et al. Massively parallel digital transcriptional profiling of single cells. Nat. Commun. 8, 14049 (2017).

88. Zhang, K., Zemke, N. R., Armand, E. J. & Ren, B. A fast, scalable and versatile tool for analysis of single-cell omics data. Nat. Methods 21, 217–227 (2024).

89. Wolock, S. L., Lopez, R. & Klein, A. M. Scrublet: Computational identification of cell Doublets in Single-cell transcriptomic data. Cell Syst. 8, 281–291.e9 (2019).

90. Lopez, R., Regier, J., Cole, M. B., Jordan, M. I. & Yosef, N. Deep generative modeling for single-cell transcriptomics. Nat. Methods 15, 1053–1058 (2018).

91. Bushnell, B. BMap: A Fast, Accurate, Splice-Aware Aligner. in 9th Annual Genomics of Energy & Environment Meeting (2014).

92. Kim, D., Paggi, J. M., Park, C., Bennett, C. & Salzberg, S. L. Graph-based genome alignment and genotyping with HISAT2 and HISAT-genotype. Nat. Biotechnol. 37, 907–915 (2019).

93. Liao, Y., Smyth, G. K. & Shi, W. featureCounts: an efficient general purpose program for assigning sequence reads to genomic features. Bioinformatics 30, 923–930 (2014).

94. Love, M. I., Huber, W. & Anders, S. Moderated estimation of fold change and dispersion for RNA-seq data with DESeq2. Genome Biol. 15, 550 (2014).

95. Korotkevich, G. et al. Fast gene set enrichment analysis. bioRxiv (2016) doi:10.1101/060012.

96. Mutte, S. K. et al. Origin and evolution of the nuclear auxin response system. Elife 7, e33399 (2018).

