## Supplemental Figures for "Positional grammar of transcription factor binding partitions developmental and stress-response regulation in plants"

### Extended Data Figures

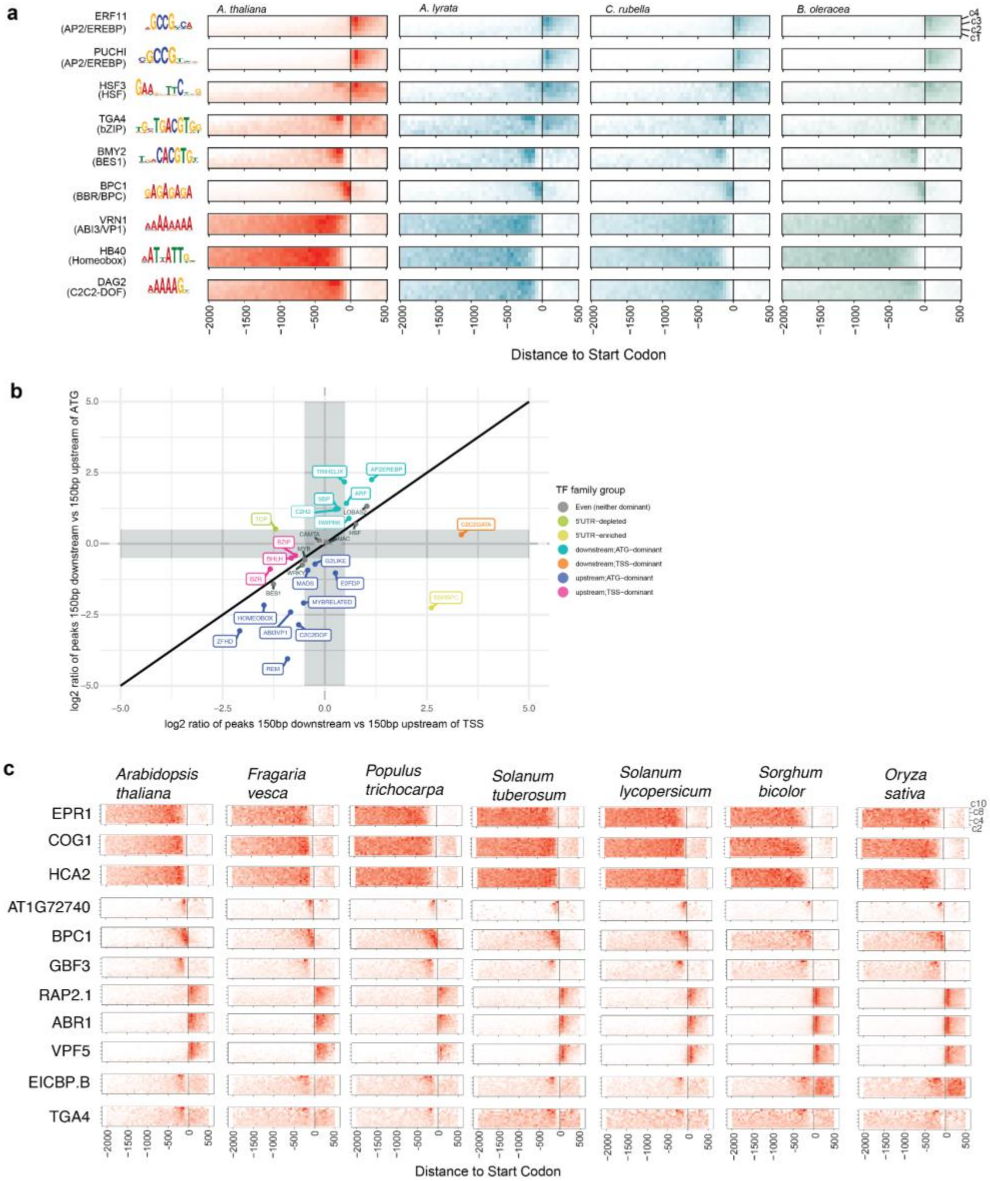


**Extended Data Fig. 1. a)** Examples of TFs with conserved TFBS location patterns across c scores, anchored on either the TSS or the start codon for *A. thaliana*, *A. lyrata*, *Capsella rubella* and *Brassica oleracea,* illustrating conservation of location patterns between species. **b)** Transcription factor binding site enrichment distinguishes TSS-driven and CDS-driven TF positioning. The x-axis shows log2 of the ratio of total DAP peaks associated with each TF family in the region 150 bp before vs. the region 150 bp after the TSS, and the y-axis shows the same for the regions 150 bp before and after the start codon. TF families where the absolute fold change between the ratios was > 0.5 are labeled as either TSS-dominant or ATG-dominant, reflecting a sharper change in TFBS density at the TSS and or ATG respectively. **c)** Locational TFBS conservation patterns anchored on the start codon for seven of the species in the deep conservation dataset.

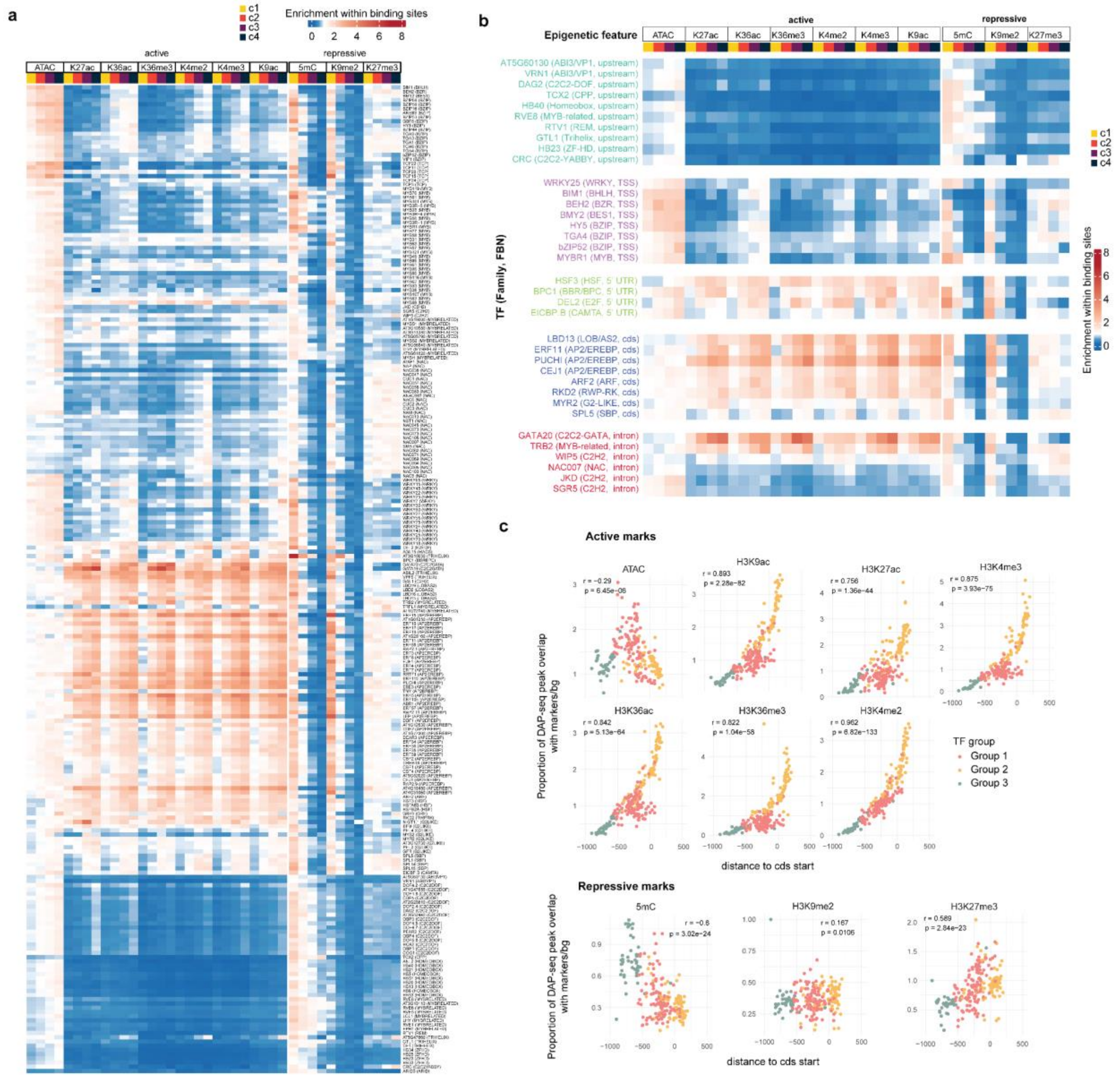


**Extended Data Fig. 2. a)** Enrichment analysis of the degree of overlap between ten epigenetic markers and TFBSs for all TFs tested, categorized by c score (colored bins). The results indicate varying levels of enrichment, from depletion (blue) to enriched (red), providing insights into the relationships between epigenetic modifications and transcription factor activity across the analyzed datasets. **b)** Selected TF from Extended Data Fig. 2a. grouped by TF family and FBN to show differences of epigenomic patterns across groups. **c)** Correlation between median distance to TSS and enrichment of epigenetic marks of all TFs. Colors represent groupings of TFs based on the epigenetic mark overlap pattern seen in Extended Data Fig. 2a.

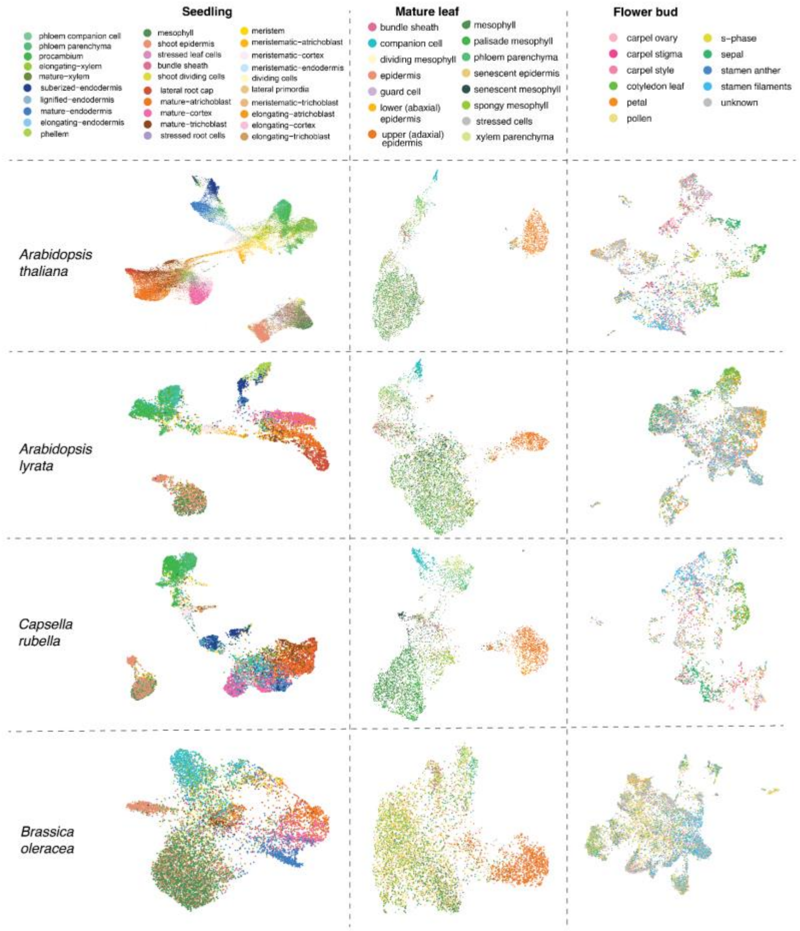


**Extended Data Fig. 3**. UMAP visualization of single-nuclei ATAC-seq data from four Brassicaceae species and three tissues. Each dot represents an individual cell, with colors indicating the specific cell type assigned to that cell based on clustering analysis. This representation allows for the comparison of chromatin accessibility patterns across different biological contexts, highlighting the diversity and distribution of cell types within the sampled species and tissues.

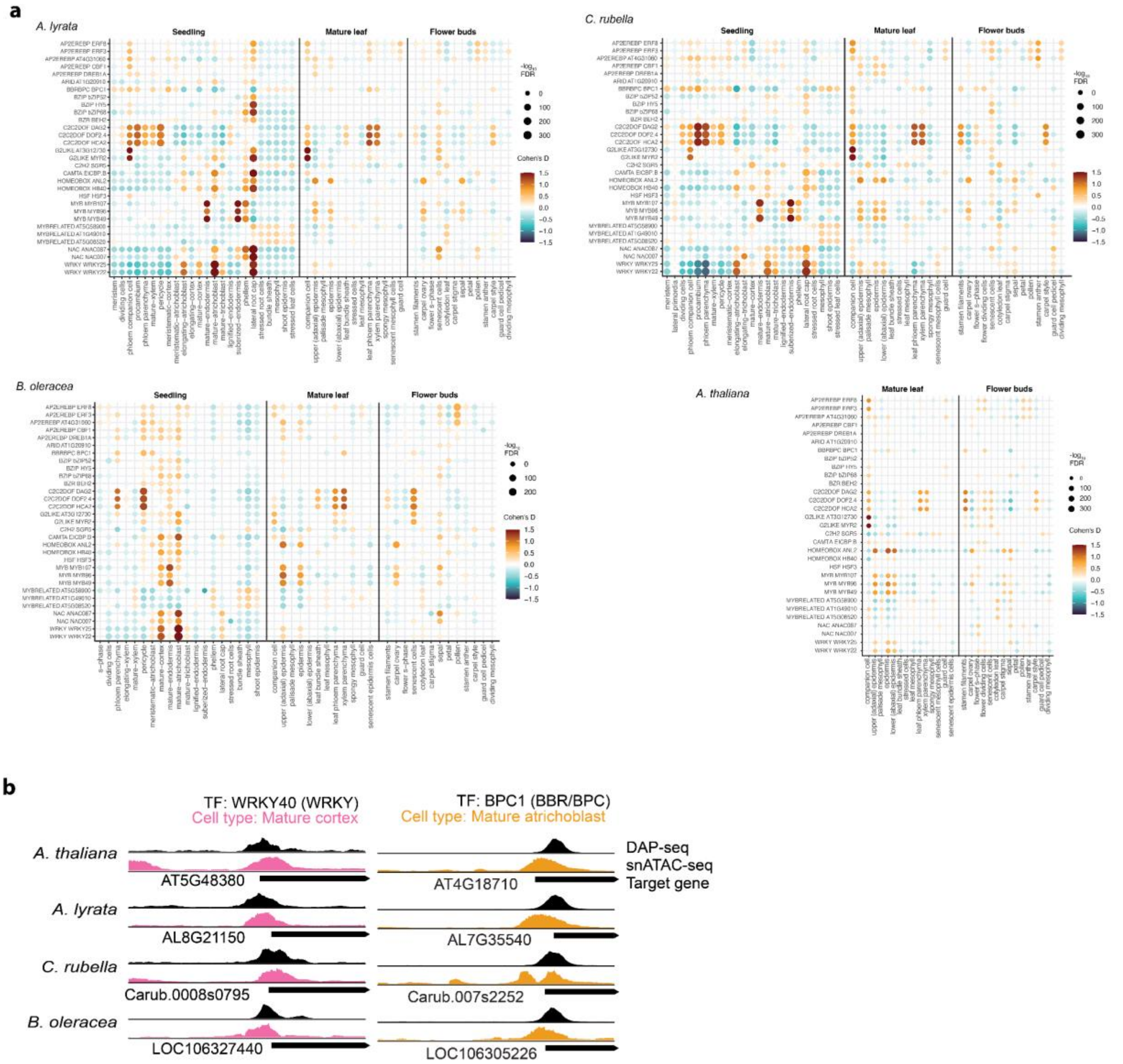


**Extended Data Fig. 4. a)** Bubble heatmaps illustrating the enrichment of transcription factor binding sites (TFBSs) in DARs from multiple tissues and species. The color of each bubble reflects the magnitude of enrichment, quantified using Cohen's D, while the size of the bubble indicates the significance of the enrichment, represented by -$\log_{10}$(FDR p-value). Only selected transcription factors are displayed from a comprehensive analysis involving four different species and three distinct tissues, highlighting patterns of regulatory activity across biological contexts. **b)** Examples of open chromatin regions in species analyzed, showcasing regions with enhanced accessibility and potential regulatory activity.

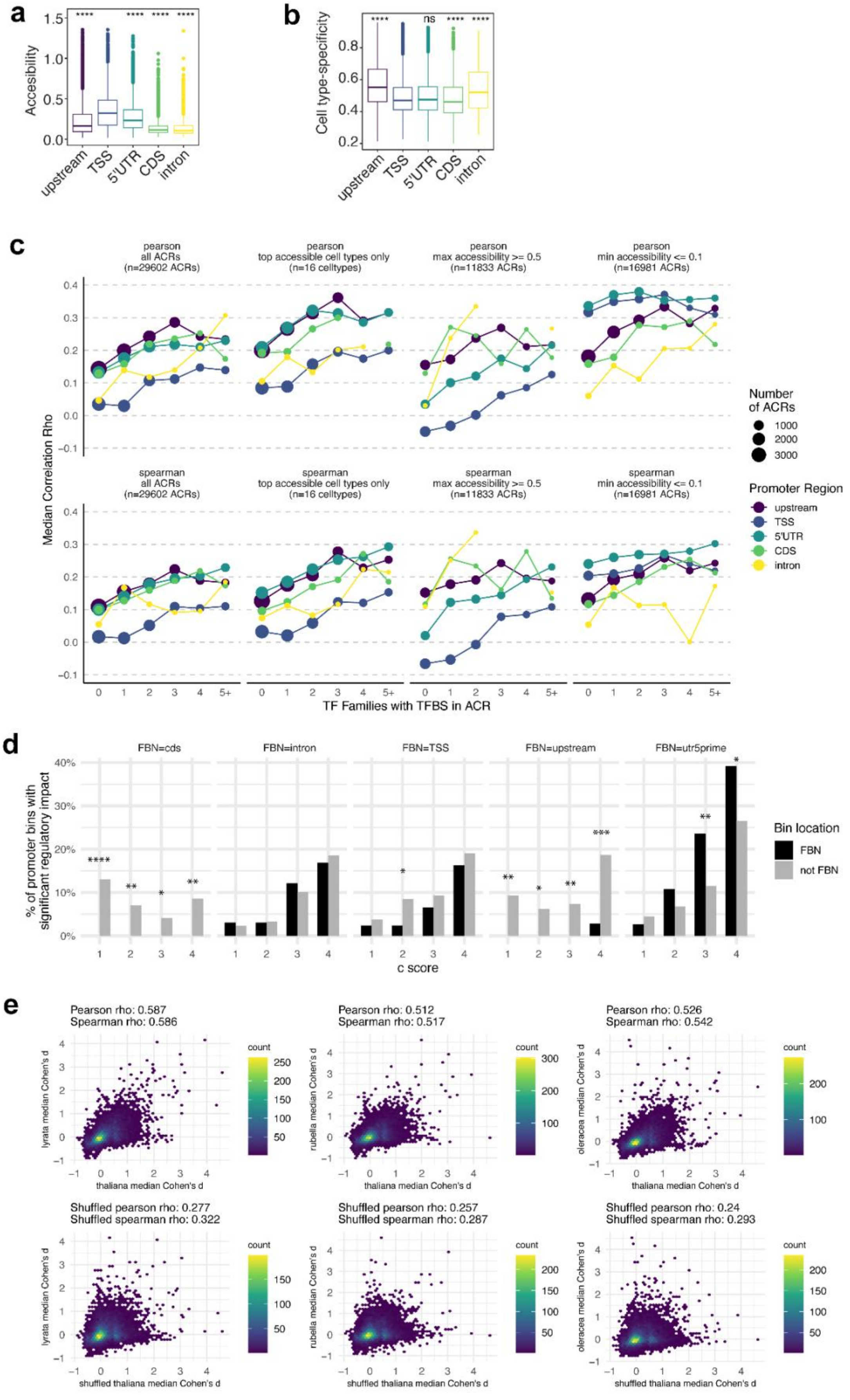
a
Accesibility
upstream
TSS
5'UTR
CDS
intron
b
Cell type-specificity
ns
c
pearson
all ACRs
(n=29602 ACRs)
top accessible cell types only
(n=16 celltypes)
max accessibility >= 0.5
(n=11833 ACRs)
min accessibility <= 0.1
(n=16981 ACRs)
spearman
Median Correlation Rho
TF Families with TFBS in ACR
Number
of ACRs
Promoter Region
d
FBN=cds
FBN=intron
FBN=TSS
FBN=upstream
FBN=utr5prime
% of promoter bins with
significant regulatory impact
c score
Bin location
FBN
not FBN
e
Pearson rho: 0.587
Spearman rho: 0.586
Pearson rho: 0.512
Spearman rho: 0.517
Pearson rho: 0.526
Spearman rho: 0.542
lyrata median Cohen's d
rubella median Cohen's d
oleracea median Cohen's d
thaliana median Cohen's d
count
Shuffled pearson rho: 0.277
Shuffled spearman rho: 0.322
Shuffled pearson rho: 0.257
Shuffled spearman rho: 0.287
Shuffled pearson rho: 0.24
Shuffled spearman rho: 0.293
shuffled thaliana median Cohen's d

**Extended Data Fig. 5. a)** Mean accessibility and **b)** cell type-specificity of all ACRs classified by promoter region. Asterisks indicate statistical significance from independent, unpaired two-sided *t*-tests comparing each individual feature category against the TSS category (serving as the reference group). ns: $p > 0.05$, * $p \leq 0.05$, ** $p \leq 0.01$, *** $p \leq 0.001$, **** $p \leq 0.0001$. **c)** Parametric (top row) and non-parametric (bottom row) correlations between accessibility of an ACR and expression of the associated gene across *A. thaliana* seedling cell types. Each point represents the median correlation coefficient for a subset of ACRs binned by location within the promoter (color) and number of overlapping TFBSs from unique TF families (x-axis). The size of the circle represents the number of ACRs considered. In the second column of plots, correlations were calculated only across the 16 cell types with high overall accessibility, in the third column correlations were summarised only for ACRs with high accessibility in at least one cell type, and in the fourth correlations were summarised only for ACRs effectively closed in at least one cell type. **d)** Percent of promoter regions with a significant TF activity score, from all promoters where at least one region (at any c score) had a significant TF activity score. Results are subsetted into different panels for groups of TFs assigned to different feature-based neighborhoods (FBNs), and results for each c score are subsetted into bars representing promoter regions corresponding to the TF's FBN (black) or regions outside the TF's FBN (gray). Asterisks represent the significance of the difference between black and gray bars according to a chi-squared test (*p ≤ 0.05, **p ≤ 0.01, ***p ≤ 0.001, ****p ≤ 0.0001). **e**) Each plot represents a comparison of the median TF activity score (for 20 random subsets of target genes) for a given TF/cell type/promoter region in *A. thaliana* (x-axis) vs. the median for the same TF/cell type/promoter region in another brassica species (y-axis). Point density is shown with color, and parametric and non-parametric correlation coefficients are shown at the top. The bottom row of plots shows the same calculations after shuffling the promoter region label for A. thaliana values within the same TF and cell type.

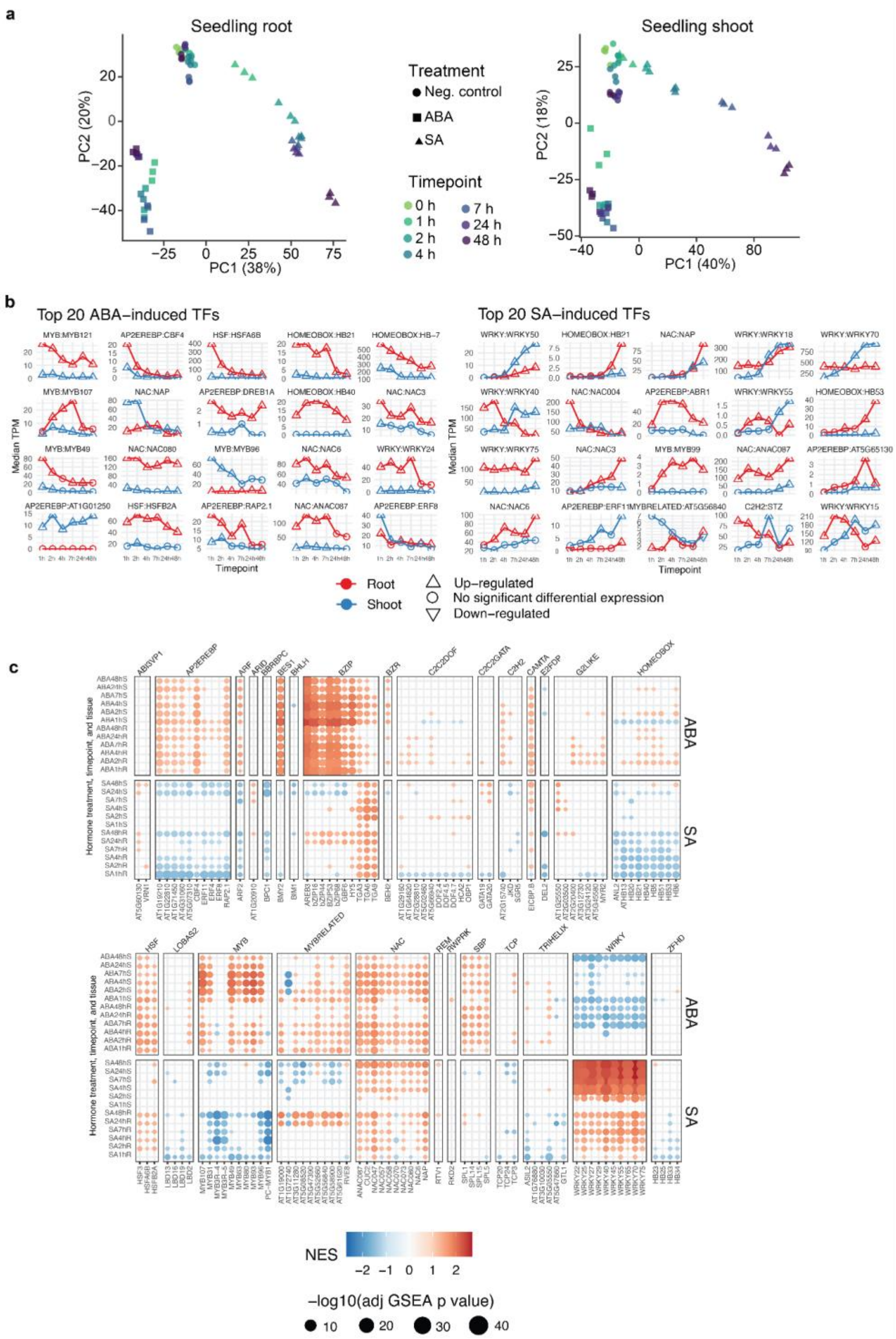


**Extended Data Fig. 6**. **a)** Principal component analysis of genome-wide transcriptional profile for root and shoot samples from *A. thaliana* plants exposed to a 48-hr treatment time course with either abscisic acid or salicylic acid. **b)** Time course of median TPM values across three replicates for top 20 TFs showing hormone-induced expression in each treatment. **c)** Results of gene set enrichment test for all c4 target genes of each TF among genes ordered by their hormone response in each tissue at each timepoint. NES = Normalized Effect Size.

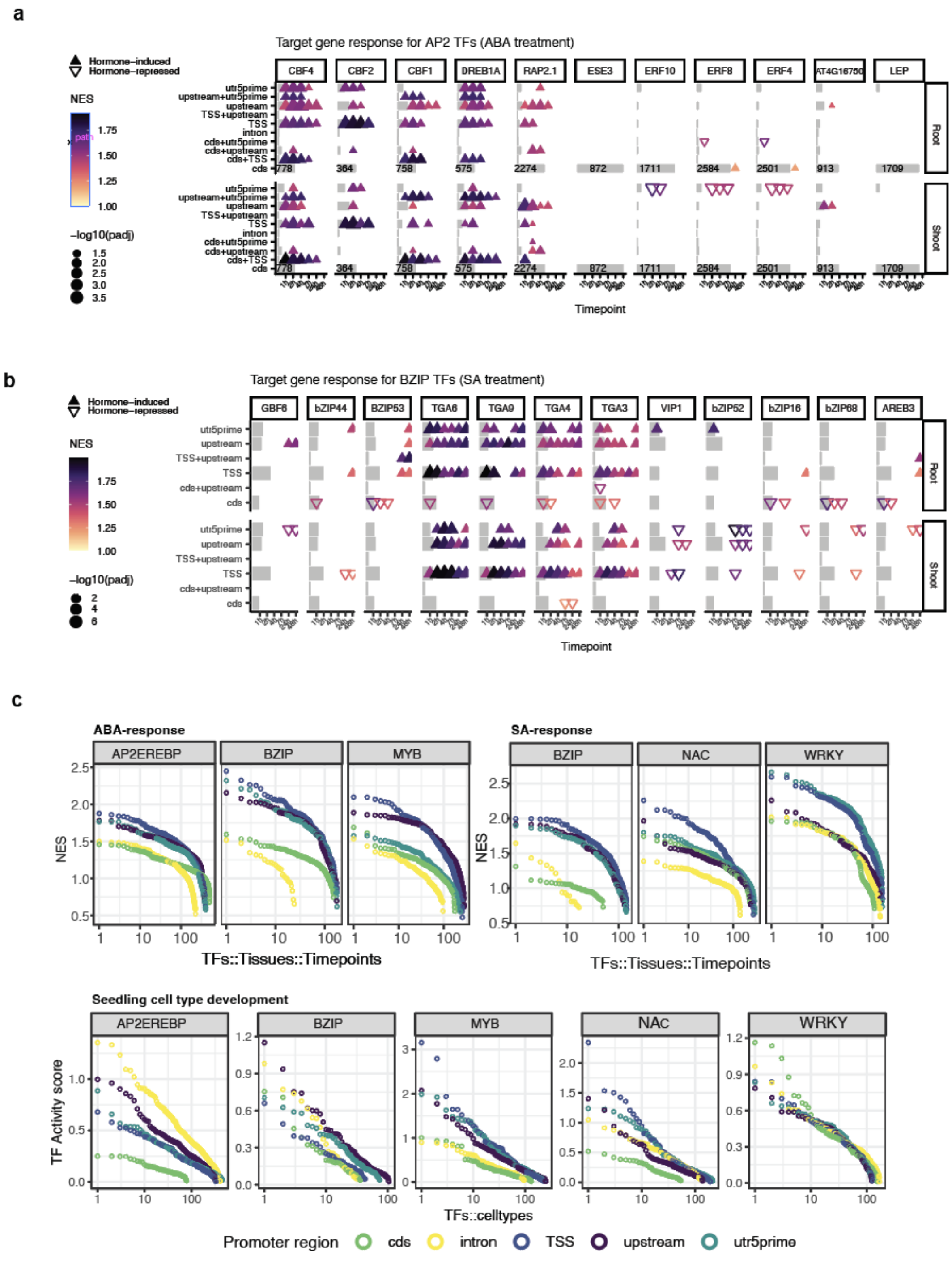


**Extended Data Fig. 7.** Gene set enrichment analysis for target genes of selected bZIP TFs during ABA treatment (**a**) and SA treatment (**b**). Gray bars represent the proportion of all target genes for the TF that fall into each TFBS location category. **c**) Top NES values for each promoter region across all TFs, tissues, and timepoints for three top-responding TF families for each hormone (top panel), and top TF activity scores representing target gene cell type specificity for each promoter region, for the same set of TF families.

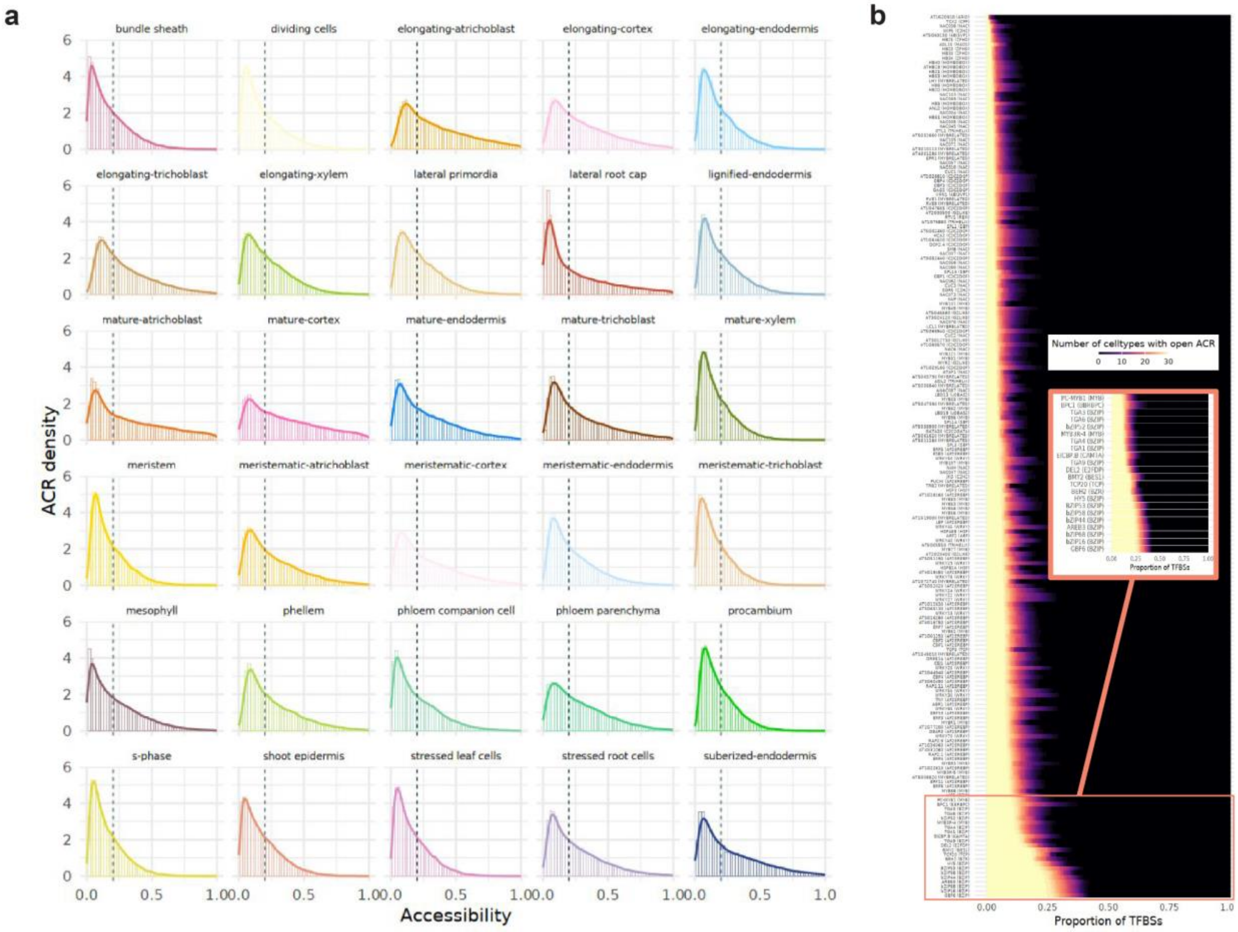


**Extended Data Fig. 8. a**) Distribution of mean accessibility values across all promoter ACRs for each cell type. Dotted line represents the threshold used for defining binary ‘open’ state (mean accessibility = 0.2). **b**) Proportion of TFBSs for each TF that overlap ACRs open in a given number of cell types. TFBSs not overlapping an ACR were coded as 0. Inset shows a zoomed-in view of the set of TFs with the highest proportion of TFBSs in constitutively-accessible chromatin.

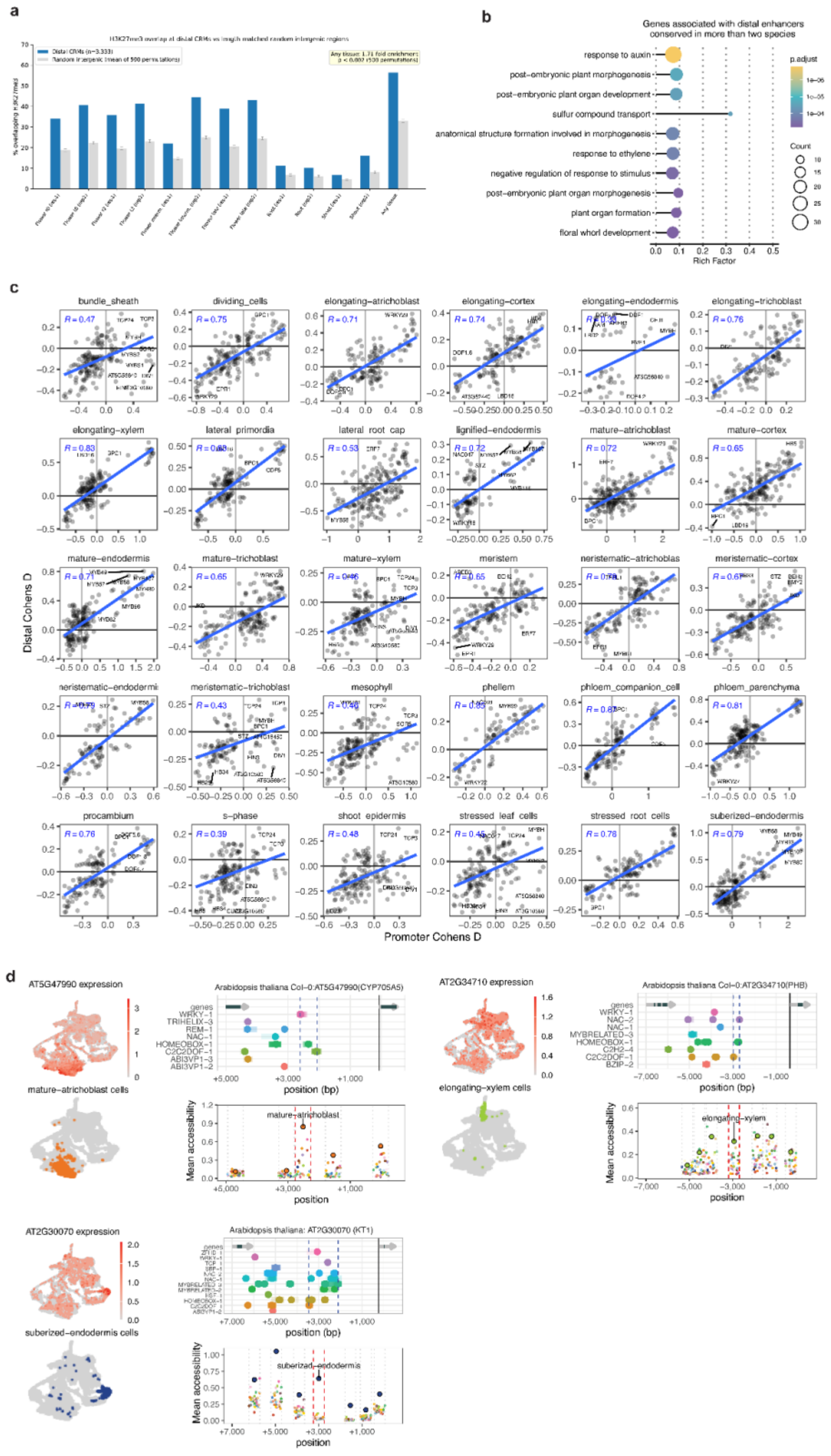

**Extended Data Fig. 9. a)** H3K27me3 enrichment at distal CRMs. Grouped bar chart showing the fraction of distal CRMs (n = 3,333) overlapping H3K27me3 peaks (blue) compared to the mean fraction from 500 length-matched random intergenic region permutations (grey), across 12 H3K27me3 ChIP-seq samples from flower developmental stages (t0, t2, intermediate, late), root, and shoot tissues, plus a union across all tissues ("Any tissue"). CRMs show 1.45-fold enrichment for H3K27me3 overlap relative to random intergenic regions (any-tissue union, empirical $p < 0.002$). **b)** Significantly enriched (adjusted $p < 0.01$) GO terms for genes associated with distal CRMs conserved in two species or more. **c**) Scatterplot of TF enrichment magnitudes (Cohen's d) for cell types from DAP-seq peaks in c4 promoter vs distal CRMs in ACRs. **d**) Other examples of candidate enhancers. The left side shows expression of the gene associated with the candidate enhancer across single *A. thaliana* seedling cells (top) and the location of cells annotated as the cell type with highest overall expression (bottom). The right side shows DAP-seq peaks (top) in and near the candidate enhancer region (dotted lines), and mean accessibility values (bottom) for each cell type (color) for ACRs in the same region. ACRs overlapping the candidate enhancer region are outlined in red, other ACR boundaries are shown with dotted gray lines.
